# Transient architecture of the embryonic pancreas determines endocrine mass

**DOI:** 10.64898/2026.06.30.735678

**Authors:** Neha H. Ahuja, Tyler Bierschenk, Christopher Chaney, Tuli Pramanik, Austin Mills, Peter M. Luo, Mitzy A. Cowdin, Jinlong Lin, Jun Tsunezumi, Kevin M. Dean, Denise K. Marciano, Thomas J. Carroll, Ondine Cleaver

**Author notes:** Corresponding author: Ondine Cleaver, Department of Molecular Biology, University of Texas Southwestern Medical Center 5323 Harry Hines Blvd., NA8.300, Dallas, Texas 75390-9148, USA.

## Abstract

During organogenesis, epithelial tissues undergo extensive three-dimensional (3D) remodeling while simultaneously generating specialized cell types. Whether these transient architectural states actively instruct lineage allocation remains unclear. Here we identify a morphogenetic stage in which resolution of epithelial stratification is required for lineage allocation and establishment of endocrine cell mass. We show that loss of the Hippo pathway regulator Merlin disrupts lumen morphogenesis and prevents formation of the transient 3D epithelial architecture that characterizes normal pancreas development. Failure to establish this architectural state alters lineage allocation, impairing acinar differentiation, markedly reducing adult endocrine cell mass, and disrupting glucose homeostasis. Mosaic analyses reveal that these lineage defects arise non-cell autonomously, demonstrating that epithelial architecture itself instructs cell fate decisions. Mechanistically, Merlin coordinates PI3K-regulated polarized membrane trafficking required for apical membrane biogenesis and lumen formation. Together, these findings identify Merlin-dependent membrane trafficking as a mechanism coupling epithelial morphogenesis to lineage allocation and demonstrate that transient developmental architectures can determine the cellular composition of mature organs.

## INTRODUCTION

How tissue architecture influences cell fate remains a fundamental question in organogenesis. Increasing evidence suggests that morphogenesis and fate are reciprocally coupled, with mechanical forces feeding into gene regulatory networks that influence cell identity.^1,2^ The developing pancreas provides a powerful model to study this relationship because perturbations in epithelial remodeling profoundly alter lineage allocation.^3,4^ During pancreas formation, a transient, stratified epithelium remodels into a branched organ that generates endocrine, acinar, and ductal lineages.^5^ This transition is driven by *de novo* lumen formation, whereby nascent microlumens coalesce into an epithelial plexus.^6^ Because endocrine cell production during development establishes the endocrine reserve of the adult pancreas,^7^ understanding how epithelial remodeling influences lineage allocation has important implications for lifelong organ function. Whether this transient architectural state merely accompanies differentiation or actively instructs lineage allocation is unknown.

In epithelial model systems, lumen formation begins at an Apical Membrane Initiation Site (AMIS) at the interface of two adjacent cells.^8–11^ At this site, polarity complexes and polarized membrane trafficking cooperate to establish the nascent apical domain and initiate lumen formation. These events are accompanied by junctional remodeling that stabilizes the emerging apical surface. Although AMIS formation and trafficking pathways have been characterized in *in vitro* epithelial cyst models, these processes *in vivo* during organogenesis have received less attention.^3,11–13^

Epithelial morphogenesis is increasingly recognized as being coupled to mechanotransduction pathways that sense and respond to tissue architecture. Studies from 3D culture of pancreatic progenitors demonstrate that lumen formation is highly sensitive to mechanical forces, including lumen pressure.^14^ The Hippo signaling pathway has been identified as a key sensor of physical cues, linking cytoskeletal tension and cell-cell adhesion to transcriptional control of growth and differentiation.^15–19^ In addition, PI3K signaling regulates pancreatic epithelial organization, intracellular transport, and mechanotransduction.^20^ These observations raise the possibility that mechanotransduction and membrane trafficking cooperate to build transient architectures that instruct lineage allocation.

Here, we investigate the role of the upstream Hippo pathway component MERLIN (encoded by *Nf2*) in coordinating epithelial remodeling and lineage specification during pancreatic development. We show that MERLIN localizes to nascent lumens and is required for proper lumen formation and epithelial remodeling. Pancreas-specific deletion of *Nf2* disrupts lumen morphogenesis, prevents epithelial destratification, and alters lineage allocation. Single-cell transcriptomic analysis reveals upregulation of both Hippo and PI3K signaling pathways in mutant epithelia, and genetic deletion of Yap1/Taz or pharmacological inhibition of PI3K partially rescues these defects. Together, our findings identify a morphogenetic stage in which MERLIN-dependent epithelial remodeling establishes a transient 3D architecture that determines lineage allocation and adult endocrine mass.

## RESULTS

### Loss of Merlin impairs pancreatic growth and function

To investigate the role of MERLIN in pancreas formation, we generated pancreas-specific Nf2 knockout mice (*Nf2^f/f^;Pdx1^Cre^*, or *Nf2^PancKO^*). Although *Nf2^PancKO^* mice survived to adulthood (**Extended data 1. A**), they exhibited marked pancreatic hypoplasia (**Fig. 1A, B** and **Extended data 1. B**). *Nf2^PancKO^* mice also displayed reduced body weights beginning at P14; however, pancreatic mass was disproportionately reduced relative to body weight (**Fig. 1C, D**). Pancreatic dysplasia was evident by embryonic day (E)14.5, but not E11.5, when epithelial organization and progenitor cell morphology appeared normal (body and cap cells were unaffected) (**Extended data 1. C-K.i’**). Smaller pancreas size was not associated with detectable changes in proliferation (phospho histone H3, pHH3) or apoptosis (Cleaved Caspase 3, CC3), at either E14.5 or P1 (**Extended data 1. L-V**). Rare apoptotic cells were observed adjacent to cystic lumens in mutant pancreata (**Extended data 1. 1U, U’**-**W**).

**Fig.1.**
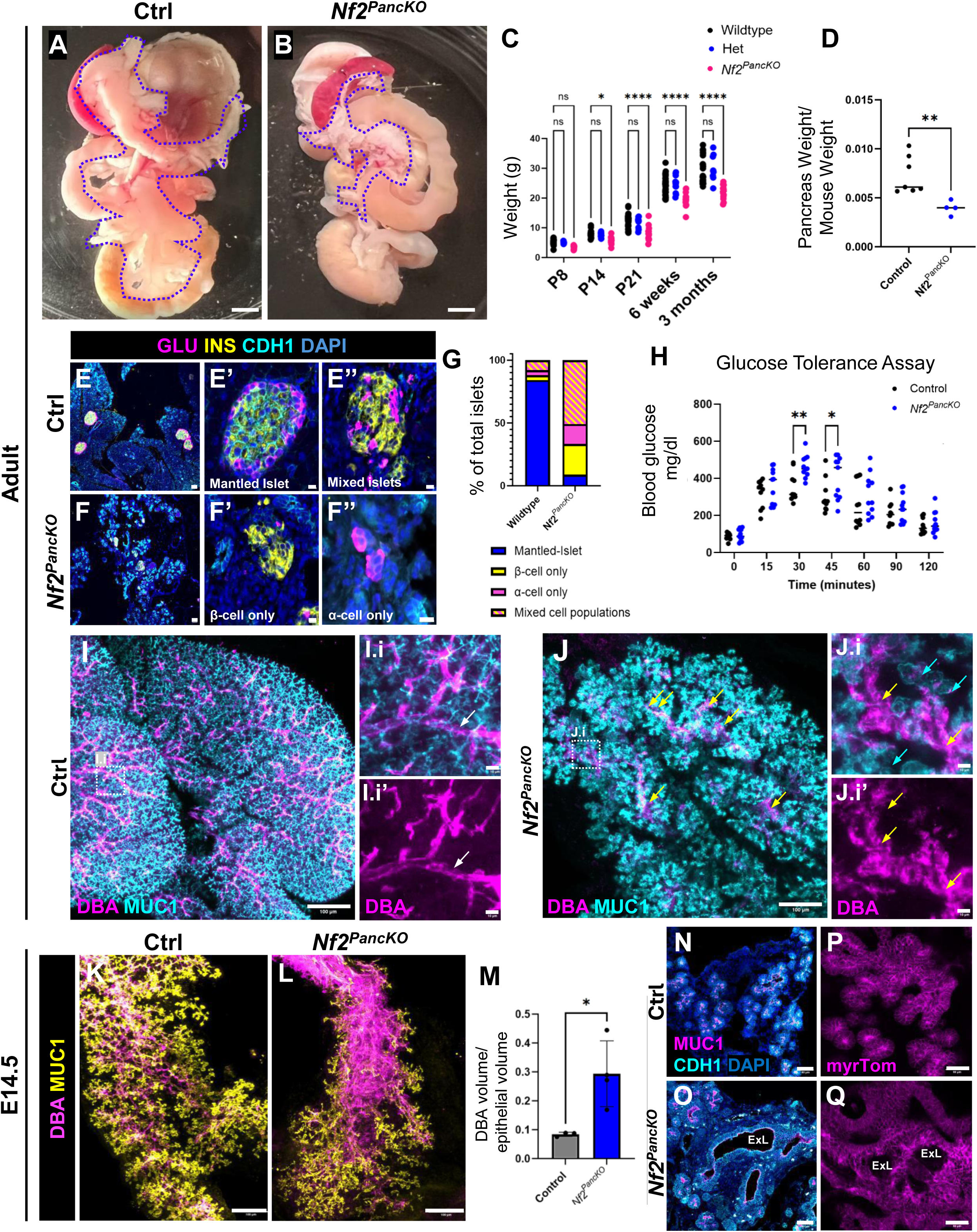
Loss of Merlin impairs pancreas growth, differentiation, and function. **A, B)** Gross morphology of digestive tract from ∼16 weeks old control (A) and *Nf2^PancKO^* (B) mice. Pancreata are outlined with blue dashed line. **C)** Body weights of control, heterozygous, and *Nf2^PancKO^* mice from P8 to adulthood. Data analyzed by 2-way ANOVA followed by Dunnett’s multiple comparisons test (ns indicates p>0.05, * indicates p <0.05, **** indicates p < 0.0001). **D)** Pancreatic weights normalized to body weight in 16-18 week-old mice. N= 7 ctrl (4 males, 3 females), and 4 mutants (3 males, 1 female). P=0.003 by Welch’s t-test. Immunofluorescence (IF) against GLU, INS, and CDH1 on adult control **(E)** and mutant **(F)** pancreata. **E’)** Representative control islet displaying the typical mantle of glucagon-positive α-cells surrounding inner insulin-positive β-cells. **F’-F’’’)** Representative dysmorphic *Nf2^PancKO^* islets. **G)** Quantification of islet morphology in control and *Nf2^PancKO^.* Each dot represents a single islet (n=17 control, 17 mutant islets evaluated from n=3 ctrl, 3 mutant animals between 16-18 weeks). **H)** Intraperitoneal glucose tolerance test in 15–18-week-old control and *Nf2^PancKO^* mice. N= 10 ctrl (4 male, 6 female), 11 *Nf2^PancKO^* (5 male, 6 female). Analyzed by 2-way ANOVA, with Tukey’s multiple comparison test, p= 0.0022 at 30 minutes, and 0.0333 at 45 minutes. Other time points were not significantly different. **(I-J.i’)** WMIF of DBA and MUC1 of a single branch from the splenic lobe of control (white arrow, large duct) **(I-I.i’)** and mutant **(J-J.i’)** adult pancreas. Yellow arrows, DBA^+^ lesions; cyan arrows, MUC1^+^ dilated lumens. Whole mount immunofluorescence (WMIF) of control **(K)** and *Nf2^PancKO^* **(L)** E14.5 pancreata, stained for MUC1^+^ and DBA. **M)** Quantification of ductal lumen diameter (N = 3 ctrl, 4 mutant pancreata. p=0.03 by Welch’s t-test). Representative images in K,L are shown from E14.5 pancreata; quantification includes pooled E13.5–E14.5 littermate pancreata. Error bars indicate standard deviation (std). **N-Q)** Section IF of control (**N, P)** and mutant (**O, Q)** pancreata at E14.5 (100μm vibratome sections). ExL, expanded lumens. Scale bars: Panels A and B 100 mm; E and F 130μm; E’-F’’ 25μm; I, J, K and L 100μm; I.i, I.i’, J.i, and J.i’ 10μm, and N-Q 50μm.

Because adult *Nf2^PancKO^* mice were smaller, we asked whether pancreas endocrine mass and function were impaired. We assessed endocrine cells by immunofluorescence (IF) for endocrine hormone markers. Control islets displayed the expected organization of peripheral glucagon-positive alpha (α) cells surrounding a core of insulin-producing beta (β) cells^21^ (**Fig. 1E, E’**), although occasional islets with mixed populations were observed (**Fig. 1E’’**). By contrast, *Nf2^PancKO^* islets were dysmorphic, smaller, and often displayed single hormone positivity, or intermingled α- and β-cells (**Fig. 1F-F’’**), with fewer β-cells (**Fig. 1G, Extended data 2. A-C**). Somatostatin-producing delta (δ) were unchanged, and pancreatic polypeptide-producing gamma (γ) cells were only modestly reduced (**Extended data 2. D-I**). Consistent with these islet defects, *Nf2^PancKO^* mice showed impaired glucose tolerance (**Fig. 1H**), with a more pronounced defect in females than in males (**Extended data 2. J, K**).

### Nf2^PancKO^ pancreata display defects in embryonic epithelial architecture

Because endocrine progenitors arise from a proto-differentiated ductal epithelium,^22^ we examined ductal morphogenesis. Whole Mount Immunofluorescent microscopy (WMIF) for the ductal marker Dolichos Bioflorus Agglutinin (DBA) revealed a highly branched ductal network in controls, with large central DBA^+^ ducts connecting to fine, peripheral Mucin-1^+^ (MUC1^+^) lumens (**Fig. 1I-I.i’**). By contrast, *Nf2^PancKO^* pancreata displayed diffuse DBA and dilation of terminal MUC1^+^ ducts (**Fig. 1J-J.i’**). These defects were evident prior to birth at E14.5 (**Fig. 1K-M**), where mutant ducts exhibited markedly expanded lumens (ExL; **Fig. 1O-Q**).

### Loss of Merlin impairs epithelial destratification

During pancreas formation, a transient stratified bud epithelium progressively resolves as the branched ductal network takes shape.^6,23,24^ We previously showed that destratification is driven by *de novo* microlumen formation and lumen extension.^6,25^ Here, we assessed epithelial organization upon loss of MERLIN to test for alteration of the cellular neighborhood during this critical time in development. By E14.5, control pancreata consisted largely of polarized epithelial monolayers surrounding narrow lumens, whereas *Nf2^PancKO^* pancreata retained extensive stratified epithelium (SE) and exhibited expanded lumens (**Fig. 2A-C, and Supplementary Movies 1-4**). Light sheet imaging of WMIF confirmed a significant increase in epithelial cell layers around mutant lumens in 3D (**Supplementary Movies 3 and 4**).

**Fig. 2.**
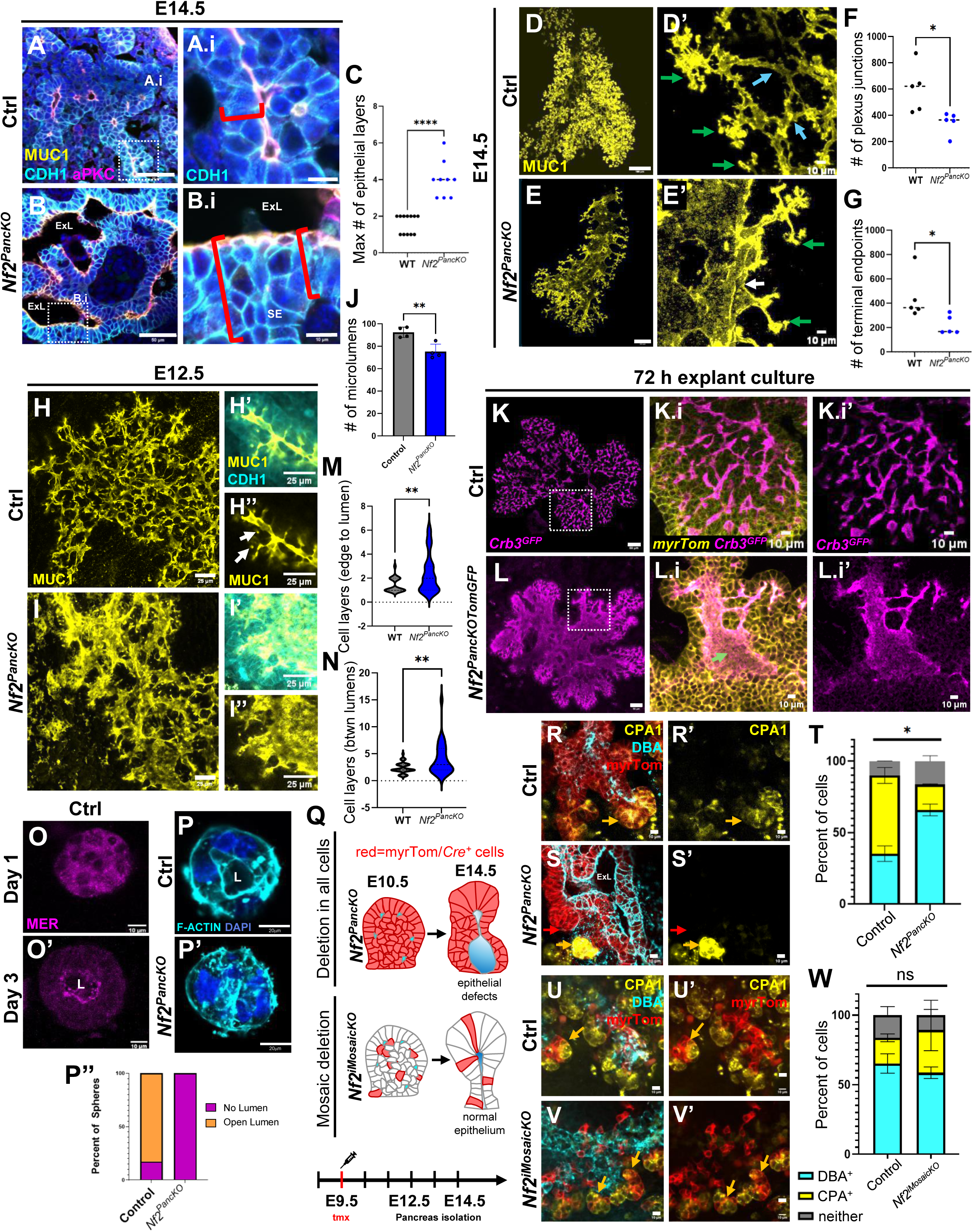
Pancreatic lumen formation and epithelial remodeling require MERLIN. **A-B.i)** Representative 10 µm sections of control (A) and mutant (B) pancreata. Red brackets mark epithelial layers between apical surfaces and epithelial edge. **C)** Quantification of maximum epithelial layers per field of view (FOV) (n=4 control, 3 *Nf2^PancKO^* mutants; ****p < 0.0001, unpaired t-test). **D, E)** WMIF Max Intensity Projections (MIPs) of E14.5 control (D) and *Nf2^PancKO^* (E) luminal networks. **D′, E′)** High magnification substacks highlight plexus junctions (blue arrows) and terminal endpoints (green arrows). **F, G)** Quantification of lumen junctions and terminal endpoints per pancreas (p = 0.0207, 0.0382, unpaired t-test). **H, I)** WMIF at E12.5. **H′, I′′)** High magnification views showing *de novo* microlumens (white arrows). **J)** Quantification of microlumen number per pancreas (p = 0.0071, Welch’s t-test). **K, L)** MIPs of explants after 72 h culture. Boxes indicate regions shown in optical sections in **K.i–L.i′**. Green arrow indicates enlarged lumen. **M, N**) Quantification of epithelial layers from edge to lumen (M) and between lumens (N) (40 control, 30 mutant measurements from 4 control, 3 mutant explants; p = 0.0033, 0.0061, Welch’s t-test). **O)** WMIF shows MERLIN localization in wildtype spheres at Day 1 and Day 3, becoming strongly apical by Day 3. L, lumen. **P)** Pancreatospheres generated from E12.5 control and mutant tissue (96 h culture), stained for F-actin and DAPI. Mutant spheres fail to form central lumens (n = 23 control spheres/3 embryos; 20 mutant/2 embryos). **Q)** Experimental strategy for lineage tracing and mosaic deletion. Tamoxifen (tmx) induction at E9.5 yields ∼10% labeled epithelial cells by E11.5–E14.5, compared to widespread labeling in *Nf2^PancKO^.* **R, S)** Vibratome sections stained for DBA1, CPA1, and myrTom(RFP) show increased ductal (DBA⁺) and reduced acinar (CPA1⁺) fate in *Nf2^PancKO^* (S, S’). Orange arrows, myrTom^+^ cells showing expression of CPA1; red arrows, myrTom^+^ cells showing no CPA1 stain; red arrows (N=2344 control, 2851 mutant cells from 2 embryos/genotype; two-way ANOVA followed by Tukey’s test; P values as follows: DBA+ Control vs *Nf2^PancKO^*, <0.0001; CPA + Control vs *Nf2^PancKO^* <0.0001, Neither Control vs *Nf2^PancKO^*, 0.3463.**U-V’)** Optical sections from vibratome sections of control (U, U’) and *Nf2^iMosaicKO^* (V, V’) pancreata stained for DBA1, CPA1 and (RFP). **W)** Quantification of lineage allocation following mosaic Nf2 deletion (N=715 control, 460 mutant cells evaluated from n=5, 4 embryo respectively. P=0.88 by two-way Anova). For T,W, error bars indicate standard deviation. Scale bars: Panels A, B, K and L 50μm;D,E 50μm, A.i, B.i, D’, E’, K.i-L.i’, O, O’, R-S’ and U-V’ 10μm; H-I’’ 25μm; P, P’ 20μm.

### Pancreatic plexus formation is dependent on Merlin

Endocrine progenitors arise within the pancreatic plexus, and perturbation of this epithelial structure alters pancreatic differentiation.^4,6,19^ This plexus has been proposed to provide a unique niche that incubates endocrine progenitors.^25,26^ 3D WMIF for MUC1 at E14.5 revealed marked plexus defects in *Nf2^PancKO^* pancreata, including fewer lumen junctions and tips (**Fig. 2D-G**). These defects emerged by E12.5, when mutant pancreata formed significantly fewer microlumens (**Fig. 2H-J**).

To visualize epithelial dynamics, we performed live imaging of explanted pancreata.^4,27,28^ Explants expressed myristoylated-Tomato (*myr^Tom^*) to outline cell membranes and Crumbs3-GFP (*Crb3^GFP^*) to mark apical lumen surfaces. Crumbs3 is an apical transmembrane protein.^29^ After 72 hours (hrs) of culture, *Nf2^PancKO^;myr^Tom^;Crb3^GFP^* (termed here *Nf2^PancKOTomGFP^*) explants retained stratified epithelium and displayed dysmorphic lumens relative to controls (**Fig. 2K-N**). Although mutant and control explants grew at similar rates, mutants failed to destratify and showed branching defects by 3 days of culture (**Extended data 3. A-G**). Epithelial clefts, which presage pancreatic branches,^30^ formed at similar rates but were reduced by ∼40% in mutant explants at 48 and 72 hrs (**Extended data 3. H**).

### Merlin is required for epithelial lumen formation

MERLIN was enriched at the apical membrane of nascent lumens (**Extended data 4. A-A.i’’**). Because *Pdx1^Cre^* deletes *Nf2* around E11.5 (**Extended data 4. B-D.i’’**),^31,32^ after initial lumens have already formed, we used a pancreatosphere assay to test the direct requirement for MERLIN in *de novo* lumen formation.^4,33^ This assay allowed isolation of E12.5 pancreas control or mutant cells (with already deleted *Nf2*) and assessment of *de novo* lumen formation. In pancreatospheres, MERLIN was enriched at nascent apical lumen membranes (**Fig. 2O,O’**). Control spheres formed single central lumens by 96 hrs, while mutant spheres did not (**Fig. 2P-P’’**). These data show that Merlin is directly required for *de novo* lumen formation and indicate that the epithelial defects in *Nf2^PancKO^* pancreata arise after initial lumen and plexus formation. Moreover, they suggest that defective lumenogenesis underlies failure of epithelial remodeling in mutants.

### Disruption of epithelial architecture alters cell fate non-cell autonomously

Our previous studies showed that pancreatic cell fate depends on epithelial morphogenesis.^3,4,34^ To establish if Merlin regulates lineage allocation, we lineage-traced control and *Nf2^PancKO^* cells from the onset of Pdx1^Cre^ expression (∼E8.75) through E14.5 (**Fig. 2Q**). In controls, 38% of labeled cells adopted a ductal (DBA^+^) fate, whereas 53% became acinar (CPA1^+^) (**Fig. 2R**). In *Nf2^PancKO^* pancreata, 71% of labeled cells became ductal and only 19% became acinar, indicating a marked shift towards ductal identity.

To distinguish cell-autonomous from tissue-level functions of MERLIN, we generated mosaic Nf2 deletions and compared them with *Nf2^PancKO^* (**Fig. 2Q**). Mosaic deletion was induced at E9.5 in *Ptf1a^CreERT2^;Nf2^f/f^;myrTom^f/f^* mice (or *Nf2^iMosaicKO^*) using low-dose tamoxifen (tmx) labeling a subset of epithelial cells (Ptf1a is expressed broadly in epithelium at this stage^35^). This strategy labeled approximately 10% of epithelial cells. Importantly, lineage allocation was unchanged in *Nf2^iMosaicKO^* mutants, with similar proportions of Tom^+^ cells adopting ductal or acinar fates in control and mutant mosaics (∼60% became ductal, 25% became acinar). Thus, loss of MERLIN in individual cells was insufficient to alter fate. These data demonstrate that MERLIN regulates lineage allocation non-cell autonomously, indicating that disruption of epithelial architecture, or the cellular neighborhood, rather than loss of MERLIN in individual cells, drives the fate defects observed in *Nf2^PancKO^* pancreata.

### Merlin is expressed at the pre-lumen Apical Membrane Initiation Site (AMIS)

Our previous studies showed that E9.75-E12.5 pancreatic epithelium continuously generates *de novo* lumens that interconnect to form the ductal plexus (**Fig. 3A**).^4,6^ In MDCK cells and in developing kidney, *de novo* lumen formation begins at an Apical Membrane Initiation Site (AMIS) where Crb3, Par3, and Rab11^+^ vesicles promote apical membrane biogenesis (**Extended data 5.A**).^8,11,13,36^ These observations underscore the dependence of lumen formation on polarized membrane trafficking. Consistent with this, Crb3, Par3, Par6, Rab11, Merlin, and Muc1 were enriched at pancreatic AMISs and nascent lumens (**Fig. 3B-D’** and **Extended data 5**). These data indicate that the same AMIS machinery operates in developing pancreatic epithelium *in vivo*.

**Fig 3:**
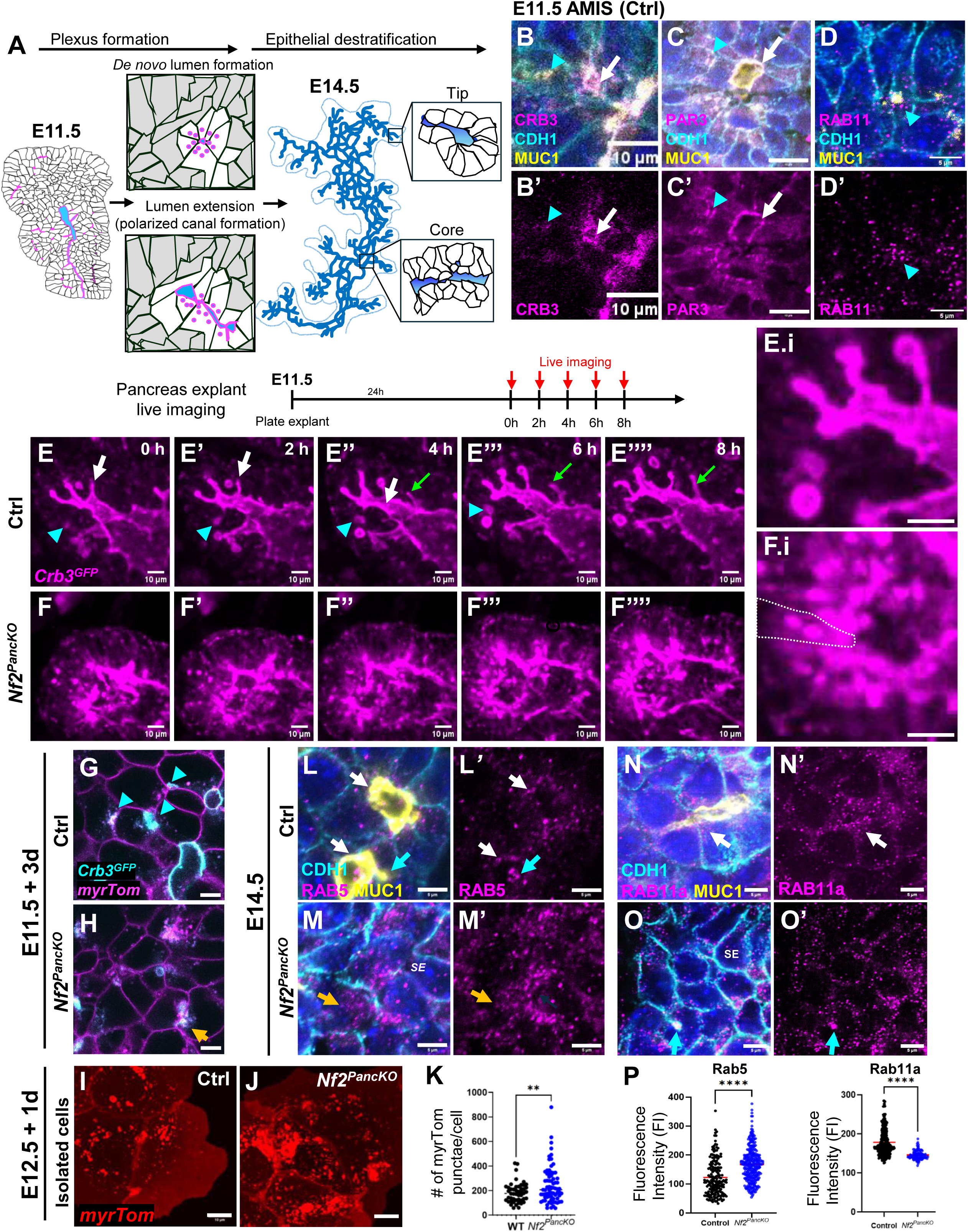
MERLIN coordinates polarized vesicular trafficking at the AMIS during pancreas lumenogenesis. **A)** Model of pancreatic lumenogenesis. At E11.5, the pancreatic epithelium is transiently stratified and undergoes *de novo* lumen formation and lumen extension via polarized vesicle trafficking. Progressive lumen growth and fusion drives destratification into an epithelial monolayer by E14.5. **B-D’)** IF showing localization of canonical AMIS markers Crb3, Par3, and Rab11 at nascent pancreatic lumens. **E-F’’’’)** Live imaging (0–8 h; MIPs; Videos 3–4) of control (E) and *Nf2^PancKO^* (F) pancreata. Control explants exhibit *de novo* microlumen formation (cyan arrowheads), lumen fusion (white arrows), and lumen extension (green arrows), whereas mutants exhibit lumen disruption and defective Crb3 trafficking. **E.i, F.i)** Higher magnification views showing Crb3 localization. **E)** In control explants, Crb3 is tightly restricted to the apical membrane, while **F)** mutants accumulate intracellular Crb3 aggregates. White dashed line indicates a single cell boundary extrapolated from myrTom labeling (see Supplementary videos). Representative images from n=3 controls and 6 mutants evaluated. Representative images from control **(G)** and mutant **(H)** explants. Mutant explants display a disorganized luminal network and intracellular Crb3. Representative images from n = at least 3 pancreata per genotype. **I, J)** Primary epithelial cells isolated from E12.5 control (I) and mutant (J) pancreata, cultured for 24 h. **K)** Quantification of intracellular myrTom+ structures. Mutant cells accumulate significantly more intracellular vesicles than controls (n = 48 control cells from 2 pancreata, and 73 mutant cells from 3 embryos. P=0.0044 by Welch’s t-test). **L-M’)** IF for RAB5 in E14.5 control and mutant pancreata. RAB5 is localized along the apical membrane in control pancreata (white arrows, L, L’). In mutant stratified epithelium (SE) regions (M, M’), RAB5 is accumulates at nascent lumens (cyan arrows) and in the cytoplasm (orange arrow). **N-O’)** In mutant SE at E14.5, RAB11 levels are decreased (representative images from n=2 embryos per genotype). **P)** Quantification of RAB5 and RAB11 fluorescence intensity/cell (N > 100 cells analyzed per genotype, p <0.0001 for both RAB5 and RAB11 by Welch’s t-test. Red line indicates mean). Scale bars: Panels B-F.i’ and I-J 10μm; G,H L-O’ 5μm.

### MERLIN is required for apical targeting of Crb3

To test whether Merlin is required for AMIS formation, we examined apical polarity proteins in control and *Nf2^PancKOTomGFP^* pancreatic explants cultured *ex vivo*. Loss of MERLIN disrupted both *de novo* lumen initiation (arrowheads) and lumen extension (arrows) (**Fig. 3E-F.i,** and **Supplementary Movies 5 and 6**). Moreover, Crb3 failed to accumulate at the apical membrane and instead remained intracellular (**Fig. 3G, H** and **Extended data 6. A-F.i**). In controls, Crb3 was tightly restricted to the apical surfaces of nascent and extending lumens, whereas mutant epithelia displayed diffuse intracellular and basolateral Crb3 accumulation. When microlumens formed in mutant epithelia, they were sometimes intracellular (**Supplementary Movies 7 and 8**). These findings indicate MERLIN is required for apical membrane assembly during pancreas lumenogenesis.

### Merlin is required for AMIS and de novo lumen formation, but not for maintenance of existing apical identity

In the persisting stratified epithelium of *Nf2^PancKO^*, Muc1^+^ lumens were absent (**Extended data 6. G-H.i’**). However, Par3 remained expressed and was scattered along the cell cortex. This suggests that Merlin is required to focus polarity components into discrete AMIS and initiate lumen formation.

Loss of Merlin produced two distinct epithelial phenotypes: 1. Perduring stratified epithelium (SE), and 2. Expanded lumens (ExL) (**Fig. 2A-C** and **Extended data 7**). These phenotypes likely reflect the timing of *Nf2* deletion by Pdx1^Cre^ (see Supplementary Fig. 4). Epithelial cells that lose MERLIN before AMIS establishment fail to initiate *de novo* lumen formation altogether, whereas microlumens that form before Merlin depletion expand. In expanded mutant lumens, MUC1^+^ apical cells retained normal localization of aPKC and ZO-1, whereas these markers were absent from Muc1-stratified cells (**Extended data 8. A-D.i’’, E-H.i’’**). We next evaluated cell polarity using Golgi positioning. In control epithelia, the Golgi (anti-Golgi matrix protein 130, GM130) localized beneath the apical membrane (**Extended data 8.I-I.i’’**). Similarly, mutant cells that had an apical surface also displayed normal subapical Golgi (**Extended data 8. J-J.i’’**), similar to ZO1 and aPKC.

### Loss of MERLIN uncouples intracellular polarity from apical domain assembly

Despite disrupted Par3 localization, stratified *Nf2^PancKO^* cells retained polarized Golgi positioning (**Extended data 8. I-L.ii**). In control epithelia, the Golgi localized to beneath AMIS-associated Par3, while in mutant stratified regions Golgi and Par3 positioning became uncoupled, with individual cells displaying multiple Par3 foci. By contrast, the apical markers ZO-1 and aPKC were absent from mutant stratified epithelium (**Extended data 8. E-L.i’**). These data indicate that loss of MERLIN disrupts assembly of a coherent apical domain without abolishing intracellular polarity (much like loss of Cdc42),^23^ leaving a polarized Golgi axis intact.

### MERLIN coordinates productive apical membrane trafficking

Because Crb3 transport was disrupted in *Nf2^PancKO^* epithelia (**Supplementary Movies 9 and 10**), we quantified intracellular vesicles in isolated pancreatic epithelial cells. E12.5 control and *Nf2^PancKOTomGFP^* pancreatic buds were dissociated and cultured on glass coverslips for vesicle analysis (**Fig. 3I-K**). Mutant cells exhibited a ∼33% increase in intracellular Tom^+^ vesicles, indicating accumulation of intracellular membrane compartments.

Crb3 is an early apical determinant required for lumen formation and is trafficked to nascent apical membranes during epithelial polarization.^36^ It is one of the first components recruited to forming lumens in Madin Darby Canine Kidney (MDCK) epithelial cysts.^8,37^

Live imaging revealed elevated vesicle trafficking near nascent lumens compared with established lumens, consistent with a role in lumen assembly (**Supplementary Video 11 and 12**).

Because Crb3 is known to be trafficked to the AMIS via Rab11^+^ vesicles, we examined endosomes by IF in control and *Nf2^PancKO^* pancreas. Loss of MERLIN resulted in marked increase in Rab5^+^ endosomes and a concomitant reduction in Rab11^+^ recycling endosomes at E14.5, both in intact pancreata and isolated epithelial cells (**Fig. 3L-P, Extended data 9.**). Moreover, both Rab5^+^ and Rab11^+^ vesicles lost their normal subapical enrichment and were dispersed throughout the cytoplasm. This redistribution was confirmed by altered apical:basal fluorescence intensity (FI) ratios in lumen-lining (**Extended data 9.H**). Similarly, the trafficking components Clathrin and Cdc42, which normally localize to nascent apical domains, became mislocalized in stratified regions of *Nf2^PancKO^* by E14.5 (**Extended data 9. I-J.i’**). These defects were more pronounced in central regions of the epithelium in mutants (**Extended data 10**). Together, these data support a model wherein Merlin promotes productive apical membrane trafficking by coordinating endocytic and recycling pathways during lumen initiation.

### Loss of MERLIN alters epithelial lineage allocation

To define the molecular consequences of *Nf2* loss, we performed single-nucleus RNA sequencing (snucRNAseq) of control and mutant pancreata. We first profiled control E14.5 pancreata (**Extended data 11. A-B**), when mutant defects are already evident (see Fig. 1G-M). UMAP analysis of control and mutant E14.5 pancreata (3-4 per genotype) revealed expansion of ductal cells, loss of acinar cells, and a modest increase in mature endocrine cells (**Extended data 11. A-C**).

Compared with controls, *Nf2^PancKO^* pancreata showed increased progenitor gene signatures (**Supplementary fig. 1**). Because pancreatic multipotent progenitors (MPs) are normally largely depleted by E14.5,^38^ they were not detected in our control dataset. To account for this developmental shift, we bioinformatically integrated data from both E11.5 and E14.5 control datasets for comparison (**Supplementary fig. 2**). Pseudotime analysis revealed two principal paths to endocrine cell fate, likely reflecting the first- and secondary-wave endocrine differentiation (**Supplementary fig. 1**). These analyses identified seven distinct epithelial cell populations at E14.5, including multipotent progenitors, acinar (Aci), ductal (Duc), endocrine progenitor (EP), and endocrine cells (End).

To examine differentiation dynamics, we performed lineage trajectory analysis (**Fig. 4A**). Diffusion maps resolved four main lineages: acinar, ductal, and two different endocrine trajectories. One endocrine trajectory likely corresponds to first-wave endocrine cells (the primarily glucagon^+^ cells that arise as clusters in the embryonic pancreatic bud between E10.0-E12.5), while the other represents secondary-wave endocrine cells (the singly delaminating endocrine cells that give rise to pre- and perinatal endocrine cells).^39^

**Fig 4.**
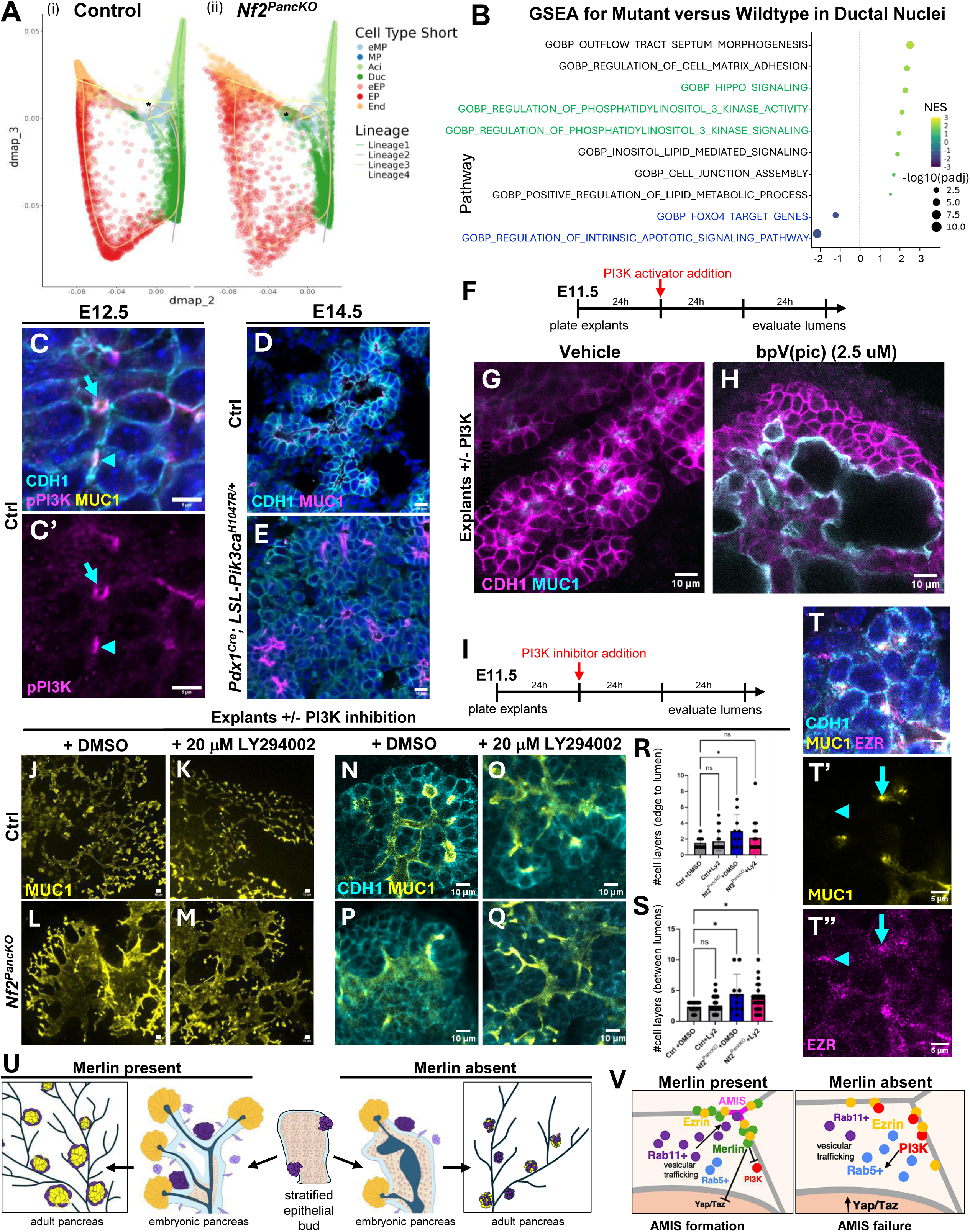
MERLIN restrains Hippo and PI3K signaling to coordinate polarized trafficking and epithelial morphogenesis. **A)** Lineage trajectory analysis of control and *Nf2^PancKO^* single-nucleus RNA-seq data (* indicates the inferred trajectory root). **B)** Gene set enrichment analysis (GSEA) of ductal nuclei reveals activation of Hippo and PI3K pathways (green) and repression of FOXO and apoptosis programs (blue) following loss of MERLIN. **C)** Phosphorylated PI3K (p85/p55) localizes to the apical membrane of nascent lumens (AMIS, cyan arrowhead; Opening lumen, cyan arrow). **D, E)** Representative E14.5 control (D) and Pik3ca-overexpression (*Pdx1^Cre^; LSL-Pik3caH1047R*) (E) pancreata (n=3 embryos/ genotype). Constitutive PI3K activation phenocopies epithelial stratification defects seen in mutants. **F-H)** E11.5 pancreatic explants cultured for 24 h and treated with vehicle or 2.5 µM BpV(pic) (PI3K activator) for an additional 48 h. PI3K activation blocks epithelial de-stratification and induces dysmorphic lumens. **I)** Experimental design for LY294002 rescue experiments. Explants treated with the PI3K inhibitor LY294002 at 24 h and analyzed at 72 h. **J-M)** MIPs of MUC1 immunofluorescence (IF) in control (J,K) and mutant (L, M) explants treated with DMSO or 20 µM LY294002. **N-Q)** High-magnification optical sections stained for MUC1 and CDH1. Quantification of epithelial layering from epithelial edge to nearest lumen **(R)** and between adjacent lumens **(S)**. PI3K inhibition partially rescues the stratification phenotype in mutants. R) Data analyzed by one-way ANOVA with Dunnett’s test (P-value for control-DMSO vs control-Ly2 = 0.93, control-DMSO vs *Nf2^PancKO^*-DMSO = 0.02, control-DMSO vs *Nf2^PancKO^*-Ly2 = 0.29). S) Data analyzed by ANOVA followed by Dunnet’s test. P-value for control-DMSO vs control-Ly2 = 0.95, control-DMSO vs *Nf2^PancKO^*-DMSO = 0.01, control-DMSO vs *Nf2^PancKO^*-Ly2 = 0.03). For both P and Q: n = 20 measurements from 2 control+DMSO explants, 30 from 3 control+Ly2, 10 from 1 *Nf2^PancKO^*+DMSO, and 25 from 3 Nf2PancKO+Ly2 explants. Error bars indicate standard deviation. **T)** IF showing EZRIN localization at nascent lumens (cyan arrowhead) in E14.5 pancreata. **U)** Working model illustrates how loss of MERLIN disrupts epithelial remodeling and results in reduced endocrine (purple/yellow), acinar (orange) and altered ductal (blue) lineages in the adult pancreas. **V)** Proposed mechanism. MERLIN functions as a cortical brake that spatially constrains Hippo and PI3K signaling, promoting polarized vesicular trafficking to the AMIS (pink), lumen formation, epithelial remodeling, and ultimately, lineage allocation. Scale bars: Panels C-C’ and T-T’’ 5μm, D, E, G, H, J-Q 10μm.

In *Nf2^PancKO^* pancreata, pseudotime analysis suggested a modest increase in MPs and impaired maturation along both endocrine trajectories (**Extended data 12. A-B’** and **Supplementary fig. 2E**). Quantification of endocrine cells at E14.5 identified by WMIF revealed a transient increase in mass that was abolished by P1 (**Extended data 12. C-K**).

By IF performed on tissue sections, however, endocrine progenitors (EP) (Neurog3^+^) and the α-to-β-cell ratio were relatively unchanged at midgestation, although there was a modest reduction of the latter (**Extended data 12. L-Q**). These data suggest that *Nf2* loss delays, rather than abolishes, endocrine differentiation, with the major deficit emerging postnatally.

By contrast, acinar loss was already pronounced in E14.5 *Nf2^PancKO^* pancreata. Expression of acinar genes, including *Amy2a1* and *Cpa1*, was markedly reduced (**Extended data 13. A**). Consistent with this, acinar mass was decreased ∼3 fold by IF staining for carboxypeptidase A1 (CPA1) and amylase (AMY, **Extended data 13. B-G**). Acinar defects persisted after birth, with co-expression of acinar and ductal markers at P1 (**Extended data 13.H-I)**. Key acinar transcription factors, including *Ptf1a*, *Foxa2* and *Gata4,* were also significantly downregulated at E14.5 (**Extended data 13. J**).

### MERLIN defines a developmental window for pancreatic cell fate determination

To determine whether Merlin is required after pancreas morphogenesis, we deleted *Nf2* following epithelial remodeling and lineage allocation, at mid-gestation. Nf2 was deleted in acinar cells at E14.5, using *Ptf1a^CreERT2^;myrTom^ff^;Nf2^ff^* (*Nf2^iAcinarKO^*) or *myrTom^ff^;Nf2^ff^* (*Cre-*negative control). Despite efficient recombination (myrTom^+^), most acinar cells exhibited no detectable acinar or ductal defects at E17.5 (**Extended data 14. A-D.i’’**). Likewise, deletion of *Nf2* in differentiated β-cells (*Ins^CreERT2^;myrTom^ff^;Nf2^ff^* pups induced at P2 and P3, *Nf2^iInsKO^*) did not affect insulin expression or islet architecture (**Extended data 14. E-F.i’’**). The evidence supports a model whereby Merlin is critically required during a restricted developmental window but is dispensable for maintenance of differentiated pancreatic cell fates.

### Hippo activation incompletely explains defective Nf2^PancKO^ morphogenesis

Gene Set Enrichment Analysis (GSEA) confirmed expected activation of Hippo pathway targets following loss of Merlin (**Fig. 4B**). Canonical Yap1/Taz targets, including *Cyr61(Cnn1)*, *Ctgf(Cnn2),* and *Ankrd1*, were significantly upregulated (**Extended data 15. A**). Increased expression was observed in both acinar and ductal populations by snucRNAseq. This suggests that pancreatic MERLIN suppresses Yap1/Taz signaling in the developing pancreas. However, combined heterozygous loss of Yap1 and Taz only partially rescued the lumen defects of *Nf2^PancKO^* pancreata (*Nf2^PancKO^;Yap1^f/+^;Taz ^f/+^;Pdx1^Cre^*), (**Extended data 16** and **Supplementary Videos 13-16**), suggesting that additional pathways contribute to the morphogenetic defects caused by MERLIN loss.

### Merlin restrains PI3K-driven trafficking programs

Transcriptional profiling further revealed activation of phosphoinositide 3-kinase (PI3K) and membrane trafficking pathways following loss of *Nf2* (**Fig. 4B** and **Extended data 17**).

Consistent with increased PI3K activity, FoxO transcription factors (*Foxo1* and *Foxo3*) and apoptosis-associated gene programs were significantly downregulated. These findings are consistent with previous studies identifying Merlin as a negative regulator of PI3K signaling in HEK293 cells.^40,41^ Genes encoding vesicle trafficking and endosomal transport, including Rab5, Rab8 and Exoc1, were also upregulated.

### PI3K activation phenocopies Nf2 loss

We next tested whether PI3K dysregulation contributes to the lumenogenesis defects observed in *Nf2^PancKO^* pancreata. We found PI3K was enriched at nascent lumens during early pancreas morphogenesis, at E12.5 (**Fig. 4C, C’**). To assess the consequences of elevated PI3K signaling, we genetically activated PI3K during pancreas lumen and plexus formation. Expression of constitutively activated Pik3ca (*LSL-Pik3caH1047R*)^42^ using the *Pdx1^Cre^* driver caused persistent epithelial stratification, closely phenocopying *Nf2^PancKO^* pancreata (**Fig. 4D, E**, and **Extended data 18**).

We next tested PI3K function pharmacologically in pancreatic explants. Indeed, activation of PI3K with the PI3K activator bpV(pic) induced dilated lumens, reduced plexus complexity, and perduring stratified epithelium, recapitulating the major architectural defects associated with loss of Merlin (**Fig. 4F-H)**. Conversely, treatment of *Nf2^PancKO^* explants with the PI3K inhibitor LY294002 partially rescued epithelial morphogenesis (**Fig. 4I-S**). Inhibitor-treated *Nf2^PancKO^* pancreata exhibited increased plexus complexity relative to vehicle-treated mutants. Together, these findings identify PI3K signaling as a key regulator of pancreatic epithelial morphogenesis and indicate that MERLIN promotes lumen formation and plexus remodeling, in part through suppression of PI3K activity.

### Merlin- and PI3K-docking protein Ezrin marks sites of lumen initiation

Because both Merlin and PI3K have been found to associate with the FERM domain-containing protein Ezrin,^43,44^ we asked whether Ezrin localizes to sites of lumen initiation, as it does in nascent lumen initiation in the kidney (PMID: 28860115). Ezrin was enriched at both AMIS and nascent microlumens in the developing pancreatic epithelium (**Fig. 4T’-T’’’**). This *in vivo* co-localization places Ezrin at sites where Merlin regulates lumen formation and interconnection of microlumens into a ductal plexus. Mechanistically, our data support a model in which Merlin coordinates apical membrane trafficking and epithelial morphogenesis by restraining Yap1/Taz and PI3K signaling at nascent lumens, potentially by competitively binding to Ezrin at the AMIS (**Fig. 4U, V**).

### Merlin couples mechanical cues and intracellular trafficking

Because Merlin functions as a mechanosensitive regulator in multiple tissues,^45,46^ we asked whether it responds to physical cues in pancreatic epithelial cells. Human pancreatic ductal epithelial cells (HPDECs) were cultured under sparse or confluent conditions to alter cell-cell contact and mechanical state. Under confluent conditions, MERLIN protein abundance was significantly increased (**Extended data 19**). Rab5^+^ endosomal compartments were similarly increased. These data indicate both Merlin and Merlin-associated intracellular trafficking respond to changes in the physical environment experienced by pancreatic epithelial cells. Together with our developmental analyses, these findings raise the possibility that MERLIN links mechanical cues to apical trafficking and epithelial morphogenesis during formation of the pancreas.

## DISCUSSION

This study identifies Merlin/Nf2 as a previously unrecognized mechanosensitive regulator of pancreas morphogenesis that couples epithelial architecture to lineage specification, including insulin-producing β-cells. Notably, Merlin is required during a restricted developmental window corresponding to lumen and plexus formation but is dispensable for early bud formation and maintenance of the pancreas. Loss of Merlin impairs *de novo* lumen formation and microlumen coalescence, resulting in defective epithelial remodeling, altered lineage allocation, and reduced endocrine mass. Rather than functioning solely through canonical Hippo signaling, Merlin coordinates a broader signaling network that integrates PI3K signaling, polarity-directed vesicular trafficking, and epithelial cell behavior. These findings support a model in which pancreatic differentiation emerges from a mechanically organized epithelial state and suggest that Merlin establishes the tissue architecture needed for faithful developmental cell fate decisions.

### Epithelial morphogenesis as an instructive regulator of lineage allocation

Across developmental systems, cell fate is determined not only by cell-intrinsic programs but also by the spatial organization of local niches, which impact cells non-autonomously. For instance in the Drosophila germarium, oocyte specification depends on the stereotypical geometry of a 16-cell cyst, with a single connected germ cell that adopts the oocyte fate.^47^ In mammalian skin, the position of a stem cell within the extended hair follicle niche predicts whether it remains quiescent or differentiates.^48^ We have shown that the pancreas represents a similar example, where the transient embryonic epithelial plexus serves as a morphogenetic niche and disruption of its lumens alters endocrine mass, including β-cells.^4,6,34,49^ When the plexus is blocked from resolving properly, as in the *Pdx1^Cre^;Afadin^f/f^;RhoA^f/f^* mutant pancreas, endocrine progenitors incubate in this niche days longer and endocrine mass is tripled.^4^ By contrast, in the *Nf2^PancKO^* pancreas, formation of the epithelial plexus is impaired, resulting in a persistent stratified epithelial state. In the early wildtype stratified pancreas, glucagon-positive cells can differentiate, but insulin-positive cells are rare. Consistent with this, we observe that glucagon-positive cells form relatively normally, whereas insulin-positive cells are markedly reduced in adult mutant pancreata. Our findings hence support the idea that tissue architecture itself is an instructive determinant of cell fate. Importantly, mosaic loss of Merlin does not alter lineage allocation, demonstrating that the differentiation defects observed in *Nf2^PancKO^* pancreata arise from disruption of tissue architecture rather than cell-autonomous changes in fate programs. These findings support a model in which the pancreatic plexus acts as an instructive morphogenetic niche whose organization governs lineage output.

A notable implication of our findings is that transient morphogenetic intermediates can exert lasting effects on organ composition. The pancreatic plexus exists only briefly during development, yet perturbations that prolong or prevent its formation profoundly alter endocrine output with consequences that persist into adulthood. These observations suggest that developmental architectures may function as temporal regulatory niches that control lineage allocation before disappearing from the mature organ. More broadly, transient morphogenetic structures in other developing tissues may similarly act as instructive intermediates that shape adult tissue composition and function.

This work also has implications for understanding how endocrine mass is established. Current models emphasize transcriptional and growth factor signaling as primary determinants of endocrine differentiation. Our data suggest that tissue architecture constitutes an additional layer of developmental control: that of an instructive cellular neighborhood. Because endocrine cell mass that is established during development influences adult glucose homeostasis, transient defects in epithelial organization may have lasting physiological consequences long after morphogenesis has ended.

### Merlin functions beyond Hippo to restrain PI3K in pancreas

Our findings support the idea that Merlin regulates signaling pathways beyond the Mst/Lats/Yap1/Taz axis. The incomplete rescue obtained by Yap1/Taz reduction further suggests that key functions of Merlin are separate from canonical Hippo signaling. Instead, our data supports a model in which Merlin regulates apical membrane biogenesis by spatially constraining PI3K-dependent trafficking pathways during epithelial lumen formation. Merlin localizes to membrane-associated polarity and cytoskeletal complexes that include Par3, Crb3 and Ezrin, positioning it to spatially coordinate signaling at the apical cortex. Prior studies have demonstrated that Merlin can directly suppress PI3K activity through interaction with the PI3K activator PIKE-L,^40^ while Ezrin and related cortex scaffolds are known to promote PI3K/Akt signaling at the plasma membrane.^43^ Because PI3K signaling strongly influences phosphoinositide composition, membrane identity and vesicle docking,^50^ aberrant PI3K activation following Merlin loss could disrupt the polarized trafficking machinery required for apical membrane biogenesis. Consistent with this model, loss of Merlin disrupts localization of Crb3-, Rab11- and Rab5-associated trafficking machinery and impairs lumen formation. We speculate that Merlin normally constrains PI3K signaling within apical cortical domains to ensure productive targeting and fusion of apical vesicles during lumenogenesis. Loss of this spatial control may lead to inefficient or ectopic vesicle delivery, impaired AMIS assembly and defective epithelial organization during pancreas development.

### Merlin interprets dynamic cell tension environments in pancreatic cells

Changes in MERLIN expression and subcellular localization with epithelial cell density support emerging models positioning Merlin as a sensor of tissue mechanics.^46^ Recent studies showing assembly of Merlin and Hippo pathway components into membrane-associated cortical signaling domains provide a framework for understanding how mechanical information is translated into morphogenetic behaviors. The density-dependent changes we observe suggest that Merlin functions as part of a mechanically responsive cortical signaling network that integrates dynamic epithelial architecture with polarity, trafficking, and developmental signaling. In the growing pancreas, Merlin may dynamically interpret local epithelial cell density and tissue geometry to spatially coordinate lumen formation and lineage allocation during organ growth.

Recent studies provide a mechanistic framework for these observations by demonstrating assembly of Merlin and Hippo pathway components at the cell cortex.^51^ Rather than functioning solely as an upstream regulator of Hippo signaling, Merlin may organize spatially restricted signaling domains that integrate mechanics, polarity, vesicular trafficking, and growth control. Such a model may explain why polarity proteins, Rab-dependent trafficking pathways, PI3K signaling components, and Hippo regulators converge during pancreatic lumenogenesis. We speculate that Merlin-dependent condensates may act as mechanically responsive organizing centers that coordinate localization of Crb3-, Rab11-, Ezrin-, and PI3K-associated complexes, thereby spatially constraining signaling and membrane trafficking during AMIS formation and lumen fusion. Disruption of these signaling domains could impair apical trafficking and impair lumen organization, altering pancreatic lineage induction within the epithelial niche and ultimately reducing beta cell mass. More broadly, our findings support the concept that transient developmental architectures function as instructive intermediates that determine the cellular composition of mature organs.

## METHODS

### Ethics statement

All animal experiments were performed in accordance with the Guide for the Care and Use of Laboratory Animals and the Animal Welfare Act, as well as protocols approved by the University of Texas Southwestern Medical Center Institutional Animal Care and Use Committee (IACUC; approval number 2017–102243, approval date 26 September 2017). All animals were observed daily, and appropriate care was provided by the veterinary staff of the UTSW Animal Resource Center, which is fully accredited by the Association for Assessment and Accreditation of Laboratory Care, International (Unit Number 000673) and by the NIH Office of Laboratory Animal Welfare (Assurance Number D16-00296).

Dams were euthanized via IACUC-approved humane methods, using carbon dioxide asphyxiation and secondary cervical dislocation.

### Mice and embryo handling

CD1 mice were used as wild type controls. *Nf2^ff^*, *Pdx1^Cre-early^*, and *R26-tdTomato^+/+^* were used for experiments in this study. For mosaic deletion experiments, Inducible deletion of Merlin was carried out by gavage of tamoxifen (tmx) (Sigma Aldrich, St. Louis, MO) at a dose of 75 mg/kg body weight to pregnant dams between E8.5 and E9.5. E11.5–E18.5 embryos were collected and dissected in PBS buffer.

Pancreata from mutant embryos were compared to WT or heterozygous tissue from littermate embryos. Genotypes were determined by PCR after O/N digestion using DirectPCR (Tail) Lysis buffer (Viagen, Los Angeles, CA) per manufacturer’s instructions. Briefly, inducible simultaneous deletions were obtained by mating *Nf2^f/f^* females with *Nf2^f/+^;Pdx1^Cre^* males. To obtain mosaic *Nf2*-deficient clones, pregnant dams were gavaged with 0.75 mg/kg tmx at E9.5. Tamoxifen induction was performed as previously described.^34^

### Generation of myrTom mice

N-myristoylated tdTomato Lox-Stop-Lox (Myr-tdTom-LSL) mice were generated by homologous recombination using the genomic ROSA26 locus. N-myristoylated tdTomato was generated by PCR from pCA-mTmG (a gift of Liqun Luo, Addgene # 26123) using primers that added 5’ and 3’ Mlu1 sites. The PCR product was ligated in Ai9 (a gift from Hongkui Zeng, Addgene #22799) using Mlu-1 site. The targeting vector was linearized by Kpn1 and then transfected into mouse embryonic stem cell line SM-1 using the UTSW Transgenic Core facility. A target allele containing the Neo-resistance cassette (Neo^r^) were selected with geneticin (G418) and diphtheria toxin A (DTA). Targeted clones were screened by long-range PCR of extracted genomic DNA, with correctly-targeted clones producing a 3.2 kb product from the 5’ arm and a 4.6kb from the downstream 3’arm. The primer sets utilized are the following: 5’arm PCR: 5’-GACTAGGGCTGCGTGAGTCTC-3’/5’-AGAAGAAGGCATGAACATGGTTAG-3’; and 3’arm PCR: 5’-CAGCAGCCTCTGTTCCACATAC-3’/5’-TGCAGTGTTGAGGGCAATCTG-3’.

For further verification, PCR was also performed for Neo^r^, loxP, and Myr-tdTomato. Three correctly-targeted ES cells (3B2, 4E11, 4H3) were injected into blastocysts obtained from C57/Bl6 females. The 4H3 clone was used for all experiments.

### Sectioning

Post dissection, pancreata were fixed in 4% PFA at 4°C overnight. For paraffin sectioning, tissues were gradually dehydrated to 100% ethanol, then rinsed twice in 100% ethanol, twice in xylene for 30 min at room temperature, a mixture of 1:1 paraplast:xylene for 10 min at 60°C, and then a series of 100% paraplast at 60°C (McCormick Scientific). The tissues were then embedded in paraplast and sectioned at 10 µm with a Biocut 2030 microtome. SuperfrostPlus glass slides (Fisher) were used. For cryosectioning, tissues were rinsed in PBS and incubated in 30 % sucrose overnight at 4°C for cryoprotection. The next day, the tissues were rinsed in OCT twice for 30 min each at room temperature. The tissues were embedded in OCT, snap-frozen on dry ice, and sectioned at 30 µm using a Leica CM-3050S cryostat. SuperfrostPlus glass slides (Fisher) were used.

### Immunostaining on sections

Paraffin sections were deparaffinized with xylene, rehydrated through ethanol wash series into PBS, permeabilized in PBS + 0.3% Triton X-100 for 10 min, and then treated with R buffer A (nuclear antigens) or R buffer B (cytoplasmic antigens) in a 2100 Retriever (Electron Microscopy Sciences). Slides were incubated in CAS Block (Invitrogen) in PBS for at least 1 hour and then in primary antibody overnight at 4°C, washed in PBS, and then incubated in secondary antibody for 2 h at room temperature. Slides were then washed again in PBS and mounted using Prolong Gold (with or without DAPI). For nuclear staining, slides were also incubated in DAPI in PBS for 10 min at room temperature prior to mounting. For cryosections, slides were baked for 30 min at 60°C, rinsed in PBS, and treated for antigen retrieval as above. For cryosections, all washes were performed in PBS. Slides were permeabilized in 0.3% Triton X-100 in PBS for 10 min, blocked in CAS-Block, and incubated in primary antibody overnight at 4°C. The next day, slides were washed, incubated in secondary antibody for 2 h at room temperature, and washed again. Mounting and nuclear staining were performed as above.

For both paraffin and cryosections, images were acquired either on the Nikon CSU-W1 SoRa confocal microscope at the Quantitative Light Microscopy Core, or the Nixon-AXR or Nikon A1R confocal provided by the UT-Southwestern Molecular Biology department.

### Whole-mount immunostaining

For embryonic tissues, immunostaining was carried out as described before.^49^ For postnatal pancreata, tissues were washed in PBS following fixation and stored in PBS overnight at 4°C. Next, tissues were dehydrated into 50% methanol, incubated in 50% methanol for 1 h at room temperature, rehydrated back into PBS, and permeabilized in PBS + 1% Triton X-100 for 2 h at room temperature. Tissues were blocked in CAS-Block for at least 2 h and incubated in primary antibody overnight at 4°C. Tissues were then washed in PBS six times for 1 h each, incubated in secondary antibody overnight at 4°C, and washed in PBS five times for 30 min each. Next, tissues were dehydrated to 100% methanol, washed three times, and mounted in BABB (one-third benzyl alcohol and two-thirds benzyl benzoate) on slides using coverslip spacers. Images were acquired either on a Zeiss LSM710 Meta confocal, or Nikon1-AXR provided by the UT-Southwestern Molecular biology department.

### Light sheet microscopy

Pancreata for light sheet microscopy were processed as above. After secondary antibody washes, pancreata were mounted in low-melting agarose and then gradually dehydrated to 100% methanol. At least 3 washes with methanol were carried out to ensure complete dehydration of agarose block and then blocks were cleared with BABB. Agarose blocks were secured to glass slides and imaged using cleared-tissue axially swept light-sheet microscopy (ctASLM), as previously described.^52^ Briefly, samples were imaged on a custom ctASLM microscope equipped with matched cleared-tissue illumination and detection objectives (54-10-12, Applied Scientific Instrumentation), four laser excitation lines (LX 405-100C, LX 488-50C, LS 561-50 and LX 637-140C, Coherent), and sCMOS camera detection (ORCA-Flash 4.0 v2, Hamamatsu). Images were acquired with a 737 × 737 µm field of view, with raw lateral and axial resolutions of 0.83 µm and 0.94 µm, respectively, and deconvolved resolutions of 0.62 µm and 0.72 µm, respectively.

### Vibratome sectioning and immunostaining

After dissection and fixation, tissue was embedded in 2.5–5% low melting point agarose (Invitrogen). 200 µm sections were cut using a Leica vibratome. Floating sections were permeabilized with Triton X-100 in PBS (1% for 1.5 h at room temperature. Sections were blocked in CAS-Block for 1 h and incubated in primary antibody mixture overnight at 4° with gentle nutation. Sections were washed in PBS (5 × 1 h) and were incubated in secondary antibody mixture overnight at 4°. Sections were washed in PBS (5 × 30 min).

Sections were placed on Superfrost Plus glass slides and cleared in a bubble of Rapiclear 1.47 (Cedarlane Labs) for at least 10 min. Slides were mounted in Rapiclear and imaged as in previous sections.

### Glucose Tolerance Test

Intraperitoneal glucose tolerance tests were performed in mice fasted overnight. Baseline blood glucose levels were measured from tail vein blood using a handheld TRUEbalance glucometer. Glucose (2 g/kg body weight) was administered by intraperitoneal injection, and blood glucose levels were measured at 15, 30, 60, 90, and 120 min post-injection.

Data were analyzed by 2-way ANOVA. Area under the curve (AUC) was calculated using GraphPad Prism.

### Dry Weight of Pancreas

Adult mice (∼18 weeks of age) were euthanized and pancreata were dissected. Adipose tissue was removed, and pancreata were briefly blotted to remove surface moisture prior to weighing on an analytical balance. Pancreatic weights were recorded in Microsoft Excel. **qPCR** Real-time qPCR was performed as described previously.^49^ Briefly, 500ng of total RNA was isolated from individual E14.5 mouse pancreata (dissected on ice) using RNeasy Microkit (Qiagen). cDNA was synthesized using SuperScript III (Invitrogen). 1μl of cDNA in Power SYBR Green Master mix (Applied Biosystems) was used for real-time qPCR analysis (QuantStudio3, ThermoFisher) of gene expression. Gene expression levels were normalized to *GAPDH* of littermate control, and the ΔΔC_t_ method was used to calculate fold change. Data were collected from individual embryos (*n* = 5 per genotype), and samples were analyzed in triplicate. Data are presented as mean (SD).

### Single nuclear RNA sequencing

#### Nuclear Isolation

For E11.5 samples, pancreata were dissected from five litters, and all pancreatic buds collected between E11.25 and E11.75 were pooled, yielding approximately 40 pancreatic buds in total. Samples were snap-frozen in liquid nitrogen and stored at −80°C until processing. For E14.5 samples, individual pancreata were collected from three control (Cre-negative, littermate) and three *Nf2^PancKO^* embryos, snap-frozen in liquid nitrogen, and stored at −80°C until processing. Nuclei were isolated from frozen tissue using the 10x Genomics Chromium Nuclei Isolation Kit according to the manufacturer’s protocol. Tissues were lysed on ice for 5 minutes prior to nuclei purification.

For E14.5 samples, nuclei from pancreata of the same genotype were pooled before library preparation. Single-nucleus RNA-sequencing libraries were generated using the 10x Genomics Next GEM Single Cell 3′ Reagent Kit v3.1 and sequenced at the UT Southwestern Next Generation Sequencing Core on an Illumina NextSeq 500 High Output Flow Cell using v2.5 chemistry. Low-quality nuclei and potential doublets were removed during quality control.

#### RNAseq data analysis

Single-nucleus RNA-seq data were processed using standard quality-control, normalization, clustering, trajectory inference, and differential expression workflows as described in the Supplementary Methods.

#### Data and Code Availability

Custom software used for diffusion map analysis, time-embedded diffusion integration, geometric harmonics projection, EIGEN marker identification, and downstream analyses is available at: https://github.com/cpchaney/pdx1-cre_nf2

### Pancreatic sphere assay and immunostaining

#### Sphere formation

Sphere assays were carried out by isolating E12.5 pancreata, dissociating pancreata into single cells and culturing in Matrigel as previously described.^33^ Dispase treatment and subsequent mesenchyme removal in PBS were each performed in a small drop of liquid in a regular dissection dish. The pancreata were transferred into 100 µL of TrypLE using forceps, and 500 µL of DMEM + FBS was used to stop trypsinization. Cell clumps were broken by pipetting up and down, and cells were centrifuged at 300g for 5 min. Cells were then resuspended in an appropriate volume of culture medium described in the original protocol. The Matrigel–cell mix was seeded on coverslips in coverglass-bottom plates. Spheres were fixed after 3 days in culture. Spheres were cultured in PEM media (DMEM F12, 10% FBS, 64ng/mL FGF2, and 10 μM Y-27632, gentamycin).

#### Immunostaining

Immunostaining was performed as follows: Spheres were rinsed with PBS on ice and fixed in 4% PFA in PBS for 10 min at room temperature. Next, spheres were washed in PBS + 0.1% NP40 (PBSN) and permeabilized in PBSN for 15 min at room temperature. Spheres were blocked in CAS-Block for 30 min, incubated in primary antibody overnight at 4°C, washed in PBS, incubated in secondary antibody for 1 h, washed in PBS again, and then imaged.

### Explant Culture

Pancreatic explants were performed as previously described.^4^ In brief, E11.5-E12.0 pancreata were dissected from embryos, and placed on a fibronectin-coated, coverglass bottom 24 well plate. DMEM/F12, 10%FBS, +gentamycin media was utilized to culture pancreata. Media was changed every other day, unless otherwise noted for drug treatment. Explants were maintained at 37°C, 5% CO_2_.

### Drug treatment

Treatment with bpV(pic) and LY294002 was performed as previously described.^53^ For LY294002 experiments, dimethyl sulfoxide (DMSO) was used as the vehicle control. For bpV(pic) experiments, sterile distilled water was used as the vehicle control. Drugs were diluted to the appropriate concentrations in explant culture medium immediately prior to use. To administer treatments, existing medium was aspirated under sterile conditions and replaced with medium containing either vehicle or drug. For all drug treatment experiments, fresh treatment medium was prepared daily and replaced every 24 hours.

#### Immunofluorescence on explants

Explants were fixed in 4% paraformaldehyde (PFA) for 10 minutes at room temperature and subsequently washed three times with PBS. Following fixation, explants were gently detached from the culture dish using forceps and transferred to glass vials. Explants were permeabilized in PBST (PBS containing 1% Triton X-100) for 1 hour, blocked in CAS-Block for 30 minutes to 1 hour, and incubated with primary antibodies overnight at 4°C. The following day, explants were washed three times with PBS, incubated with secondary antibodies for at least 1 hour at room temperature, and washed three additional times with PBS. Explants were mounted on glass slides using iSpacer tape (SunJin Lab Co.) and cleared in RapiClear prior to imaging.

#### Live imaging

All live imaging was performed on a Nikon CSU-W1 SoRa confocal microscope at the Quantitative Light Microscopy Core at UT Southwestern Medical Center. For overnight imaging experiments, images were acquired at intervals of no greater than 10 minutes for a minimum imaging duration of 8 hours, depending on microscope availability. All dual-reporter-positive explants from each litter were imaged.

Because littermate controls were not always available (for example, a litter may contain a *Pdx1^Cre^; Nf2^ff^, myrTom^f/+^; Crb3^GFP+^* mutant but no corresponding cre+ control embryos), stage-matched control explants from independent litters were used for comparison. For analysis of vesicular trafficking events, live imaging was performed at interval of 1 frame per 25 seconds (one z-stack), where a frame is defined as a complete z-stack at 0.5um intervals utilizing the 100x objective. Imaging was performed for approximately 10 minutes. All live imaging was done at 37°C with 5% CO_2_.

### 2D cell culture

#### Primary cell isolation and culture

Pancreata were dissected at E12.5, mesenchyme was manually removed by dissections with forceps post treatment with Dispase. Pancreatic buds were transferred to individual sterilized microcentrifuge tubes and dissociated using 100uL TrypLE. 500uL of DMEM/10% FBS was utilized to neutralize Trypsin. Cells were plated on a fibronectin-coated coverglass bottom plate and cultured in PEM media in standard tissue culture conditions (37°C with 5% CO_2_).

#### Human Pancreatic Ductal Epithelial Cell Culture

A vial of human pancreatic ductal epithelial cells (HDPE-H6c7)^54^ were obtained from Kerafast and cultured in Keratinocyte Serum Free Media supplemented with bovine pituitary extract and EGF. Cells were cultured on gelatin-coated chamber slides at tissue culture conditions.

#### Immunofluorescence on cells

Cells were fixed in 4% PFA for 10 minutes at room temperature. Cells were then washed three times in PBS, permeabilized for 10 minutes in PBSN (0.1% NP-40), and blocked in CAS block for at least 30 minutes. Cells were then incubated in primary antibodies overnight with gentle nutation at 4° C. The next day, cells were washed three times in PBS and incubated in secondary antibody for at least 1 hour. Cells were washed three times and mounted in either Fluoromount (HDPE cells) or imaged directly in the well.

### Image Quantification

Quantification of lineage volume, epithelial plexus architecture, cell death, proliferation, and differentiation was performed using established image analysis pipelines;^3,4,6,34,49,55^ detailed methodologies and parameter settings are provided in the Supplementary Methods. Image analysis was performed blinded to genotype whenever feasible.

### Statistical analysis

Statistical analyses were performed using GraphPad Prism version 10.5.0 (774; GraphPad Software, Inc.). Unless otherwise noted, statistical significance was assessed using Welch’s t-test. P-values are reported in the figure legends. Formal power analysis was not performed prior to data collection; however, sample sizes were chosen to be consistent with those reported in previously published studies in the field. No samples or animals were excluded from analysis unless predefined technical failure criteria were met.

## ACKNOWLEDGEMENTS

We thank DJ Pan for the *Nf2^flox/flox^* mice, and Mike Dellinger for the *LSL-Pik3ca^H1047R^* mice. We additionally thank all the members of the Cleaver Lab, Afshan Fathima Nawas, Victor Varner, Anne Grapin-Botton, DJ Pan and Elizabeth Chen for scientific discussion and critical feedback.

In addition, we would like to thank Stephen Johnson for his expert computational assistance and overall daily support. We thank the UTSW quantitative Light Microscopy Core (qMLC), for support with microscopy. We are particularly grateful for the critical support provided by members of the UTSW Next Generation Sequencing Core, including Ashwani Kumar and Chao Xing.

## FUNDING

This work was supported by DK079862 and DK106743 to OC; NIH NIGMS RM1GM145399 and CPRIT RP250571 to KMD; DK116622, DK118032, and DK141873 to DKM; and K99DK140516 and Breakthrough T1D 3-PDF-2023-1327-A-N to NA. The qMLC is funded by NIH 1S10OD028630-01 to Dr. Kate Luby-Phelps.

## AUTHOR CONTRIBUTION STATEMENT

NA and OC conceptualized the study. NA carried out most experiments, with assistance from TB, TP, AM, PML, and MAC. NA performed cell and tissue imaging and analysis, with assistance from JL and KMD. Critical reporter lines were provided by JT, DKM (*myrTom*) and TJC (*Crb3^GFP^*), and provided essential feedback and data interpretation. CC performed bioinformatic analyses. NA and OC curated data. OC provided resources. NA generated figures. NA and OC wrote the original draft of the manuscript, which was reviewed and edited by DKM, TP, AM, PL, and OC. NA and OC acquired funding.

## COMPETING INTEREST STATEMENT

The authors declare no competing interests.

## EXTENDED DATA

**Extended data. 1.**
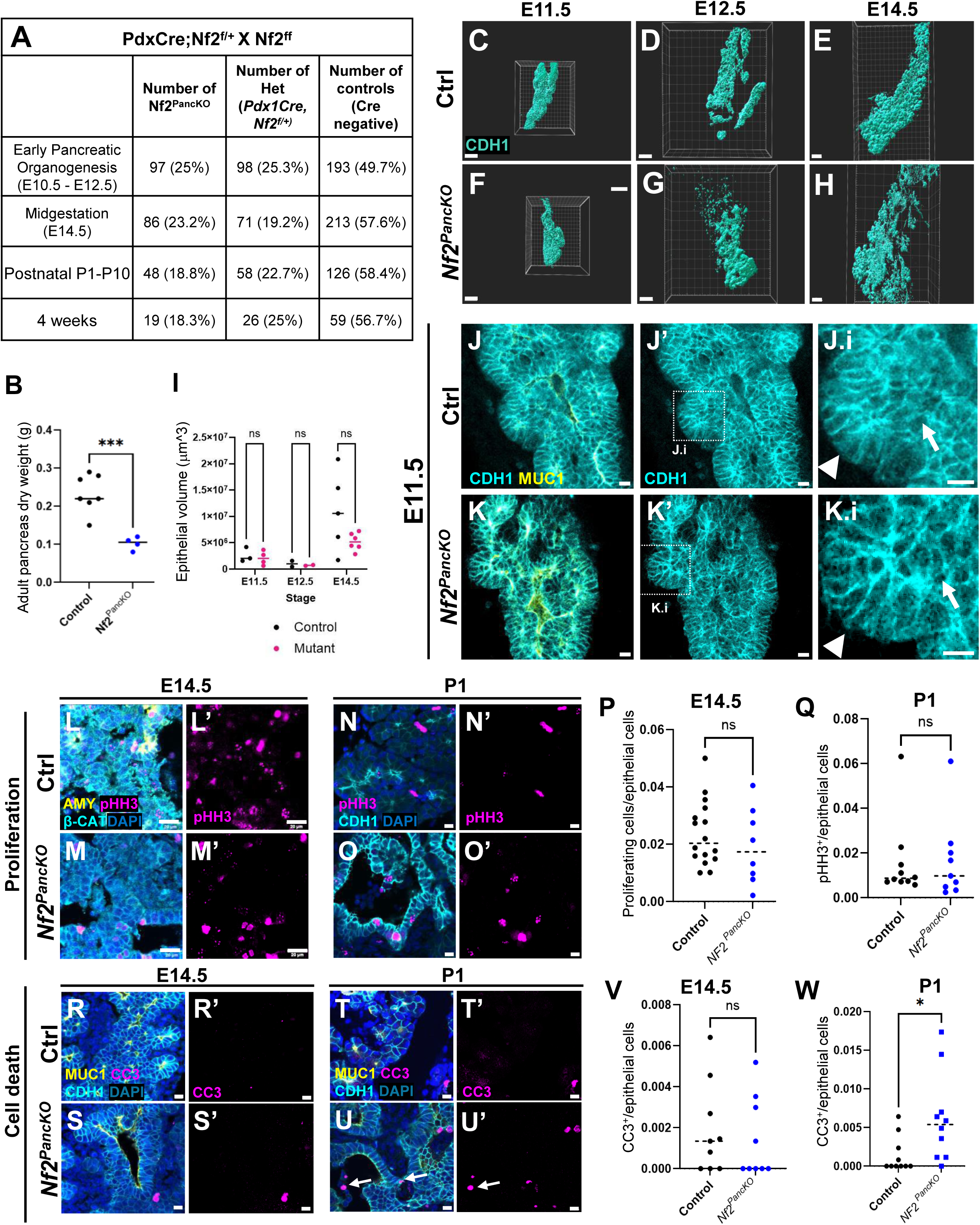
Epithelial volume and cell number are unchanged during early pancreatic morphogenesis. **A)** Number of embryos/animals per developmental stage and genotype. Expected Mendelian ratios are 25% *Nf2^PancKO^*, 25% heterozygous, and 50% Cre-negative control. **B)** Dry weight of adult pancreata. Each dot represents an individual animal, p=0.0002 by Welch’s t test. **C-H)** Imaris surface rendering of CDH1^+^ control (C-E), and mutant (F-H) pancreata from E11.5 to E14.5. **I)** Epithelial volume is unchanged during early to mid-pancreatic development in *Nf2^PancKO^*. Each dot represents one pancreas. **J-K.i’)** Representative optical section of E11.5 control and mutant pancreata. White arrowheads, cap cell at periphery of pancreas; white arrows, cuboidal body cell. IF for pHH3, β-catenin and AMY in control **(L, L’)**, and **(M, M’)** mutant pancreata at E14.5, and **(N-O’)** P1**. P)** Quantification of proliferating cells/epithelial cells at E14.5 and **Q)** P1. p=0.40, 0.909, respectively. IF for CC3, CDH1 and MUC1 in control **(R, R’)** and mutant **(S, S’)** pancreata at E14.5 and **(T-U’)** P1**. V)** Quantification of apoptotic cells/CDH1^+^ epithelial cells at E14.5, and **W)** P1. p=0.6448, 0.02, respectively. For panels P,Q,V,W: Each dot represents one FOV analyzed. N=3 ctrl and n=3 *Nf2^PancKO^* animals analyzed at each embryonic stage. Scale bars: Panels C and F 70μm, D and G 80μm, E and H 700μm, L-O’ and R-S’ 20μm, and N-O’ and T-U’ 20μm. For all graphs, line represents median of data.

**Extended data 2.**
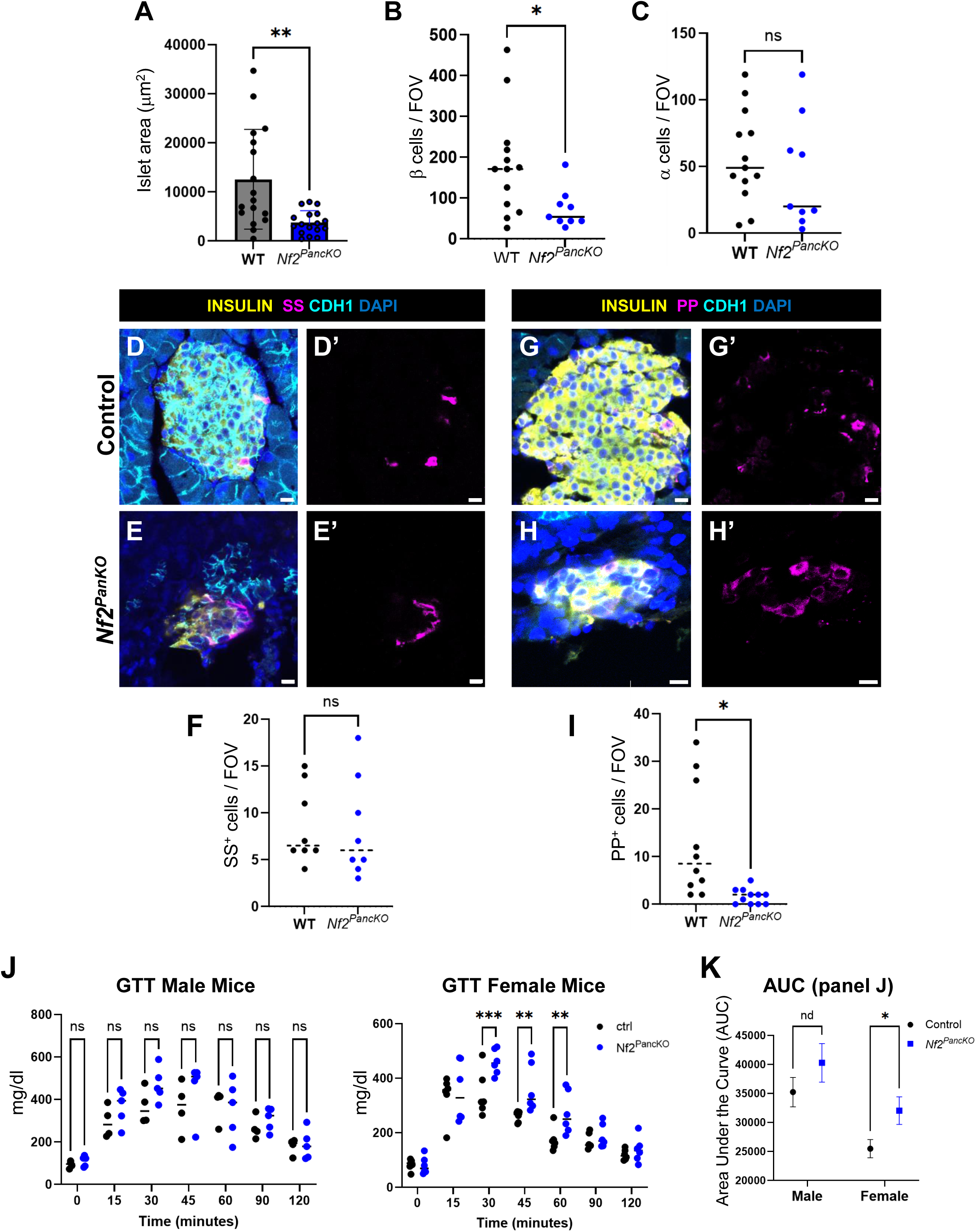
Loss of Merlin impairs isletogenesis and endocrine function. **A)** Islet area measured in pancreas sections from control and mutant pancreata as measured on section. P= 0.0015, analyzed by unpaired t-test. Each dot represents one islet. N=17 control, n=17 mutant islets were evaluated from n=3 ctrl and n=3 mutant animals (16-18 weeks). Number of β-cells **(B)** and α-cells **(C)** per field of view (FOV) in adult control and *Nf2^PancKO^*. P = 0.024, 0.438 respectively, as analyzed by unpaired t-test. For B and C, each dot represents one FOV (13 FOV analyzed from controls, 9 from *Nf2^PancKO^*, from n=3 ctrl and n=3 mutant pancreata). Representative control **(D, D’)** and mutant **(E, E’)** tissue stained for insulin, CDH1, and somatostatin (SS). **F)** Somatostatin^+^ cell numbers are not affected by loss of Merlin (p=0.8776). Representative control **(G, G’)** and mutant **(H, H’)** tissue stained for Insulin, CDH1, and Pancreatic Polypeptide (PP). **I)** PP^+^ cell number is reduced in *Nf2^PancKO^* adult tissue (p=0.0146). For D-H’, each dot represents an FOV, n=3 animals from each genotype, analyzed by Welch’s t-test. **J)** Glucose Tolerance (GTT) data in Fig. 1H, separated by sex. Glucose tolerance was unchanged in males, but significantly impaired in females. **K)** Area under the curve (AUC) analysis of glucose tolerance in male and female mice, p = 0.083 for male, 0.00043 for female, analyzed by multiple t tests with Bonferroni-Dunn correction. Scale bars: Panels D-E’ and G-H’ 10μm. For A-C, F,I,K – line represents median. For K, error bars represent standard deviation.

**Extended data 3.**
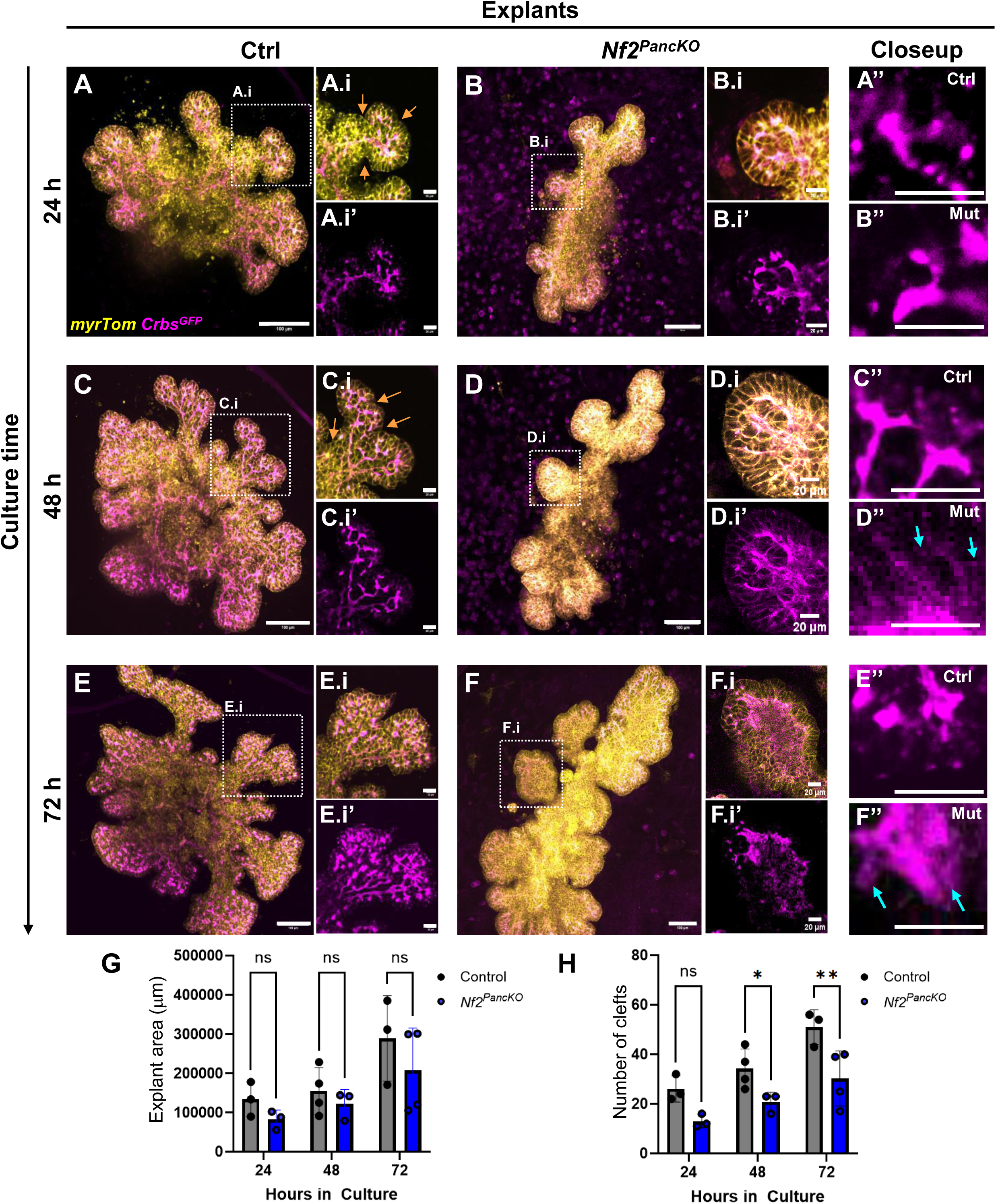
*Nf2^PancKO^* E11.5 pancreatic explants exhibit defective epithelial remodeling and branching despite normal growth. MIPs of control **(A,C,E)** and *Nf2^PancKO^* **(B,D,F)** *myrTom;Crb3^GFP^* explants at 24, 48 and 72 h of culture, respectively. **A.i-F.i’)** Single optical sections corresponding to the regions outlined in white dashed boxes in A-F. *Crb3^GFP^* single channel view of A’-F’. **A’’-F’’)** High magnification view of *Crb3^GFP^* at 24-72 h of culture. Note accumulation of intracellular *Crb3^GFP^* (blue arrows), and loss of branching clefts (orange arrows). **G)** Explant size was comparable between control and *Nf2^PancKO^* cultures, through the 72-h imaging period (p=0.26, 0.39, 0.15 at 24, 48, and 72 h respectively, using two-way Anova with Tukey’s multiple comparison test). **H)** *Nf2^PancKO^* explants display defects in branching as measured by number of clefts (p=0.051, 0.038, 0.0070 at 24, 48, and 72 hours respectively, analyzed by 2-way Anova followed by Tukey’s multiple comparison test). Scale bars: Panels A-F 100μm, and A.i-F’’ 20μm.

**Extended data 4.**
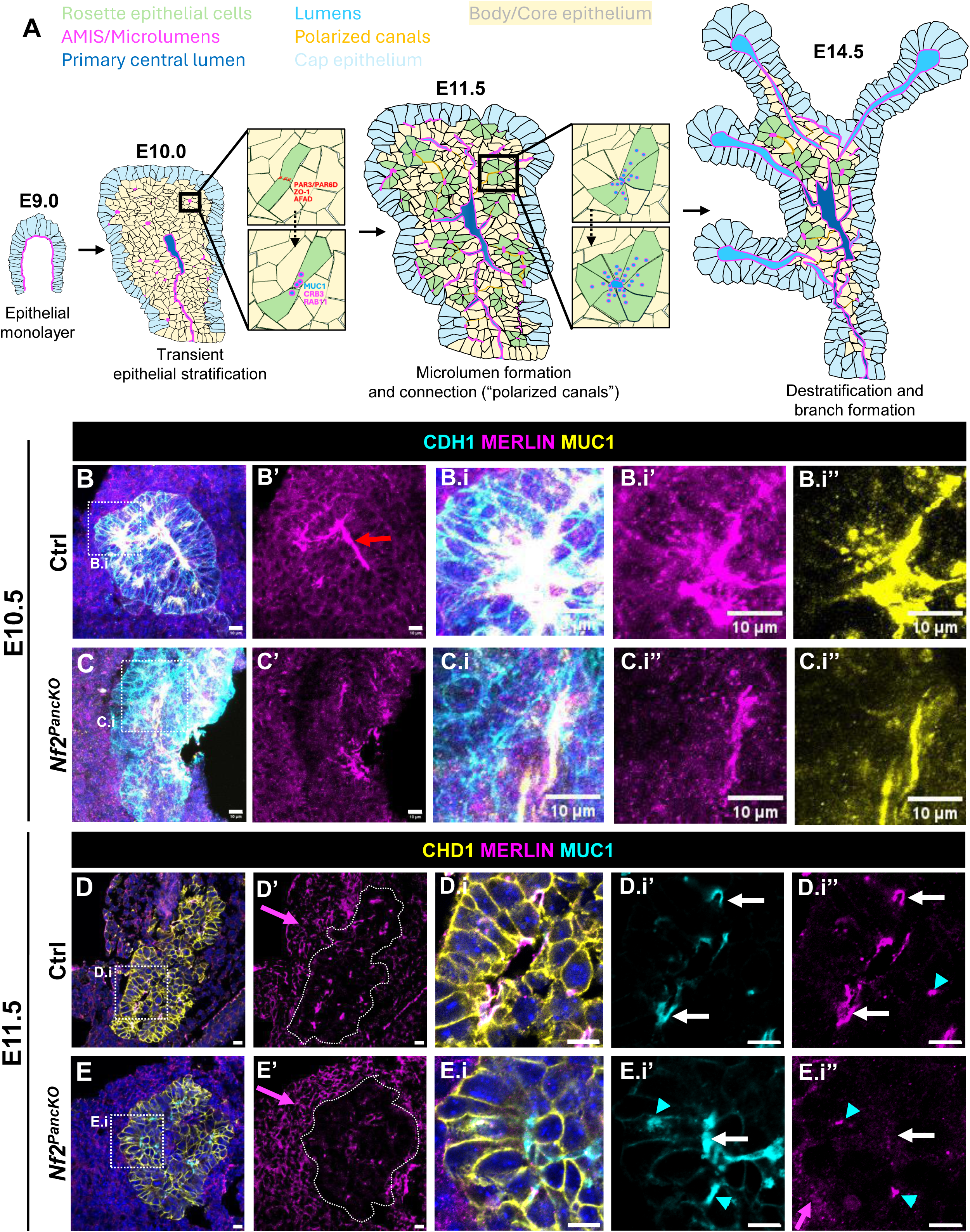
Pdx1^Cre^ deletes MERLIN in pancreatic epithelium by E11.5, but not E10.5. **A)** Schematic of pancreatic morphogenesis showing transient epithelial stratification followed by destratification and branching. Early endocrine cells arise as a first-wave of glucagon^+^ clusters, while second-wave endocrine cells delaminate individually from core epithelium. IF for CDH1, MERLIN and MUC1 in E10.5 control **(B, B’)** and *Nf2^PancKO^* **(C, C’)** pancreata (paraffin sections). Dotted lines in A’ and B’ outline the pancreatic epithelium. Red arrows indicate mesenchymal tissue. **B.i-C.i’’)** Higher magnification insets shown by boxed regions in B, C, respectively. IF for CDH1, MERLIN and MUC1 in E11.5 control **(D, D’)** and *Nf2^PancKO^* **(E, E’)** pancreata (paraffin sections). Dotted line in D’ and E’ is tracing of CDH1, indicating pancreatic epithelium**. D.i-E.i’’)** Higher magnification insets shown by boxed regions in D, E, respectively. White arrows indicate lumens; cyan arrowheads, AMIS. Magenta arrowhead indicates mesenchymal tissue expressing MERLIN. All scale bars are 10μm.

**Extended data 5.**
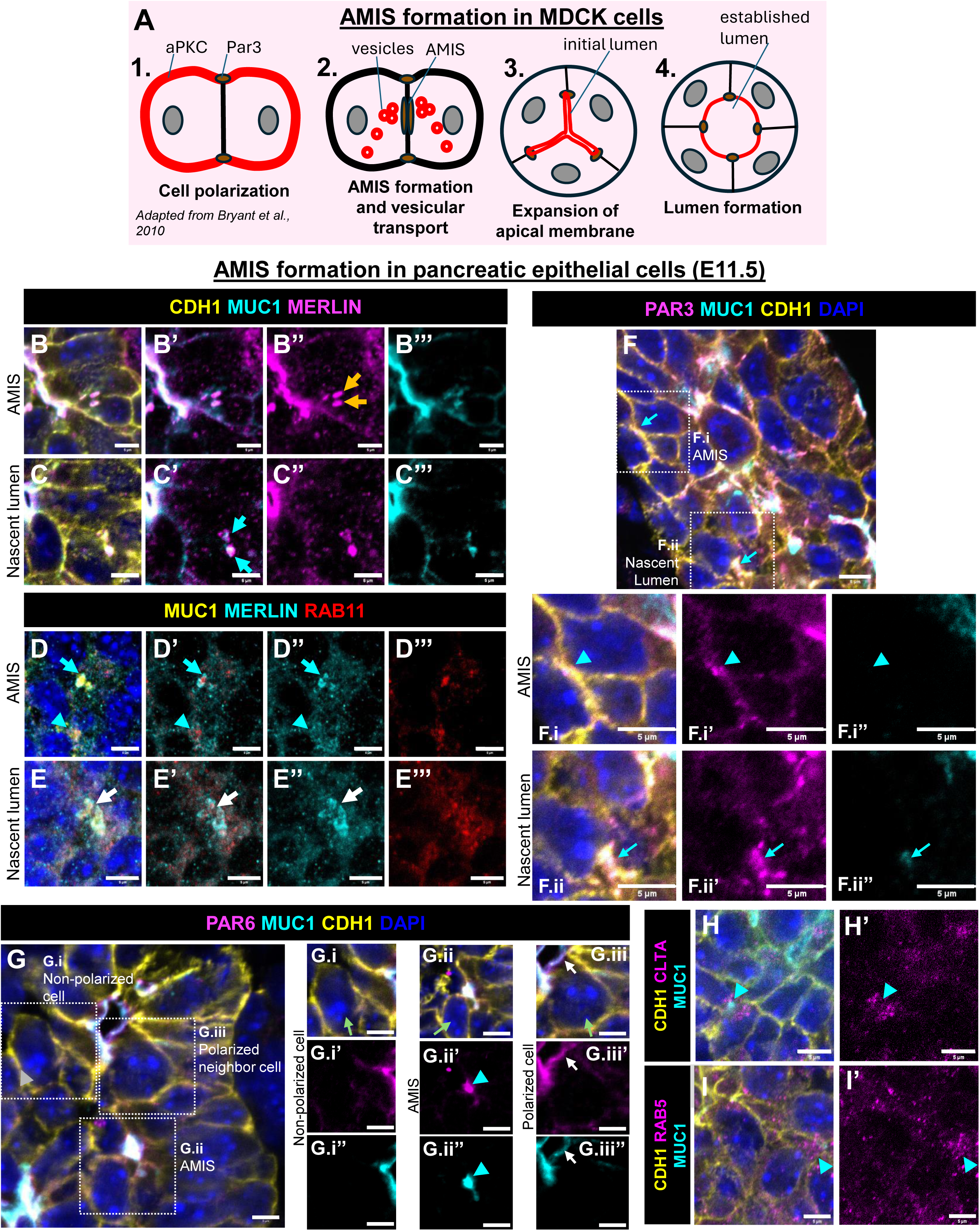
Cell polarity and AMIS components localize to nascent pancreatic microlumens. **A)** Schematic of Apical Membrane Initiation Site (AMIS) formation based on studies in MDCK epithelial cysts, illustrating RAB11-dependent delivery of apical determinants including Crb3 and Par3 during *de novo* lumen formation. **B-C’’’)** MERLIN localizes to the subapical cortex of developing lumens and partially overlaps with MUC1 in E11.5 wild-type pancreatic epithelium. Orange arrows indicate intracellular MERLIN co-localizing with MUC1^+^ vesicles. Blue arrowheads denote nascent lumens. **D-E’’’)** MERLIN and RAB11 localize to the AMIS, nascent lumens, and mature (open) lumens (cyan arrowhead, cyan arrow, and white arrow, respectively). **F)** IF for PAR3, MUC1 and CDH1 in wildtype E11.5 pancreas. **F.i-F.i’’)** PAR3 is enriched at cell-cell boundaries of developing lumens prior to detectable MUC1 accumulation, indicating the AMIS (cyan arrowheads). **F.ii-F.ii’’)** PAR3 remains enriched at the apical surface of nascent lumens MUC1^+^ lumens (cyan arrows). **G)** IF for PAR6D, MUC1 and CDH1 in wildtype E11.5 pancreas. **G.i)** PAR6D is absent from stratified epithelial (SE) cells that have not initiated lumen formation. PAR6D becomes apically enriched at the AMIS (cyan arrowheads) **(G.ii),** and polarized lumen-lining cells (white arrows) **(G.iii)**. Green arrows, cells of interest. **H-I’)** IF of wildtype E11.5 showing enrichment of CLTA and RAB5 at nascent lumens. All scale bars 5μm.

**Extended data 6.**
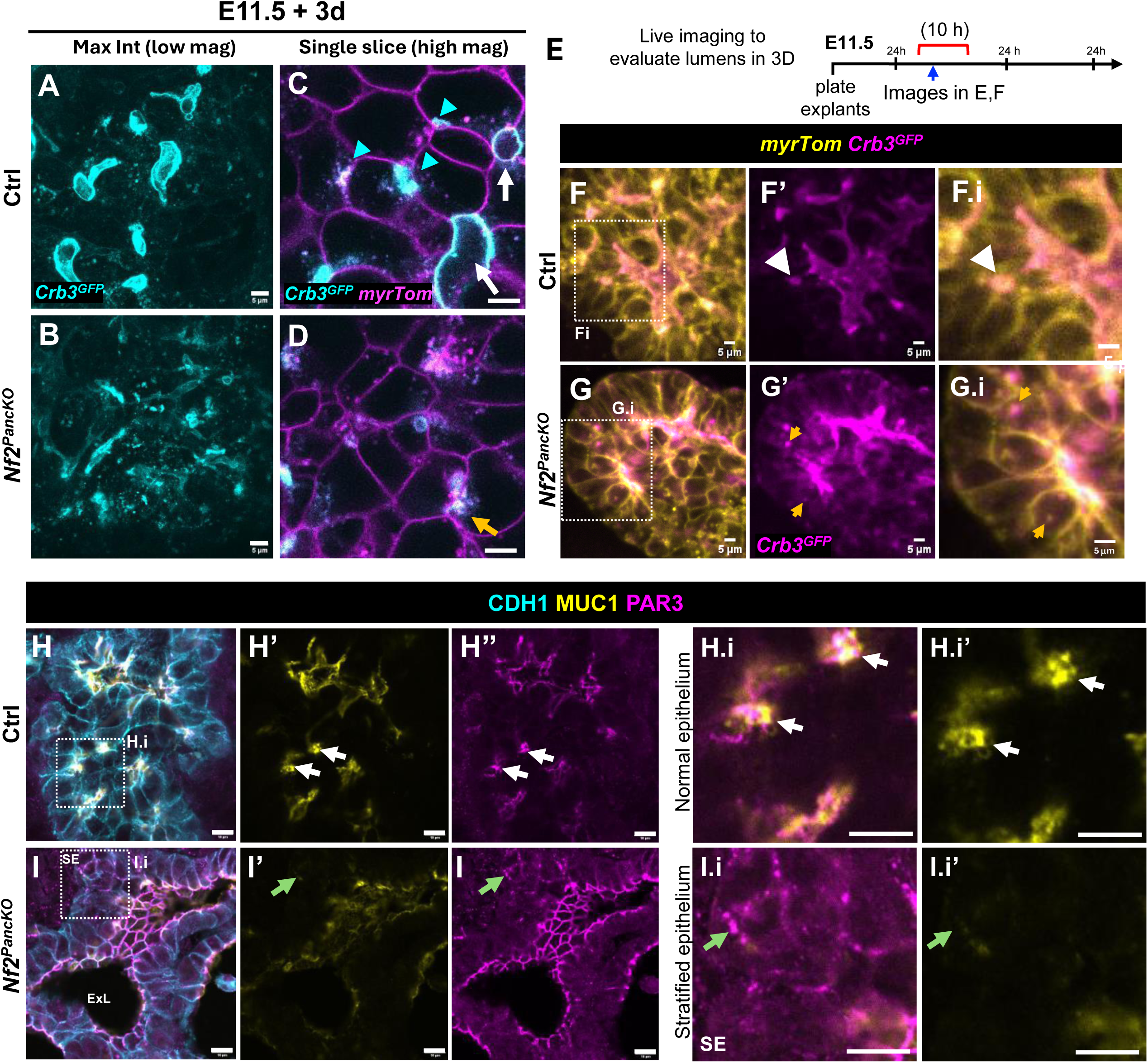
Merlin is required for pancreatic AMIS formation and polarized localization of PAR3. MIPs of control **(A)** and mutant **(B)** explants at 3 days of culture. Images in B and C were taken from approximately 2 hours of live imaging time. Panels **C** and **D** are the same as Fig. 3G-H. MIPs of *Crb3^GFP^* included here to provide context. **E)** Experimental schematic. Pancreata were explanted at E11.5 and live-imaged between 24-34 h of culture. **F)** In control epithelium, *Crb3^GFP^* localizes exclusively between adjacent cells at nascent lumens (white arrowheads). **G)** In *Nf2^PancKO^*, Crb3 accumulates in intracellular punctae, rather than at cell surfaces (orange arrows). **H)** In E14.5 control epithelium, PAR3 is tightly restricted to the apico-junctional region of lumen-lining cells (white arrows), co-localizing with MUC1. **I)** In stratified regions of *Nf2^PancKO^,* PAR3 remains detectable (green arrows), but is dispersed along the cell cortex and is not associated with MUC1, indicating failure to assemble into discrete AMIS. Scale bars: Panels A-D and H-I.i’ 10μm, F-G.i. 5μm.

**Extended data 7.**
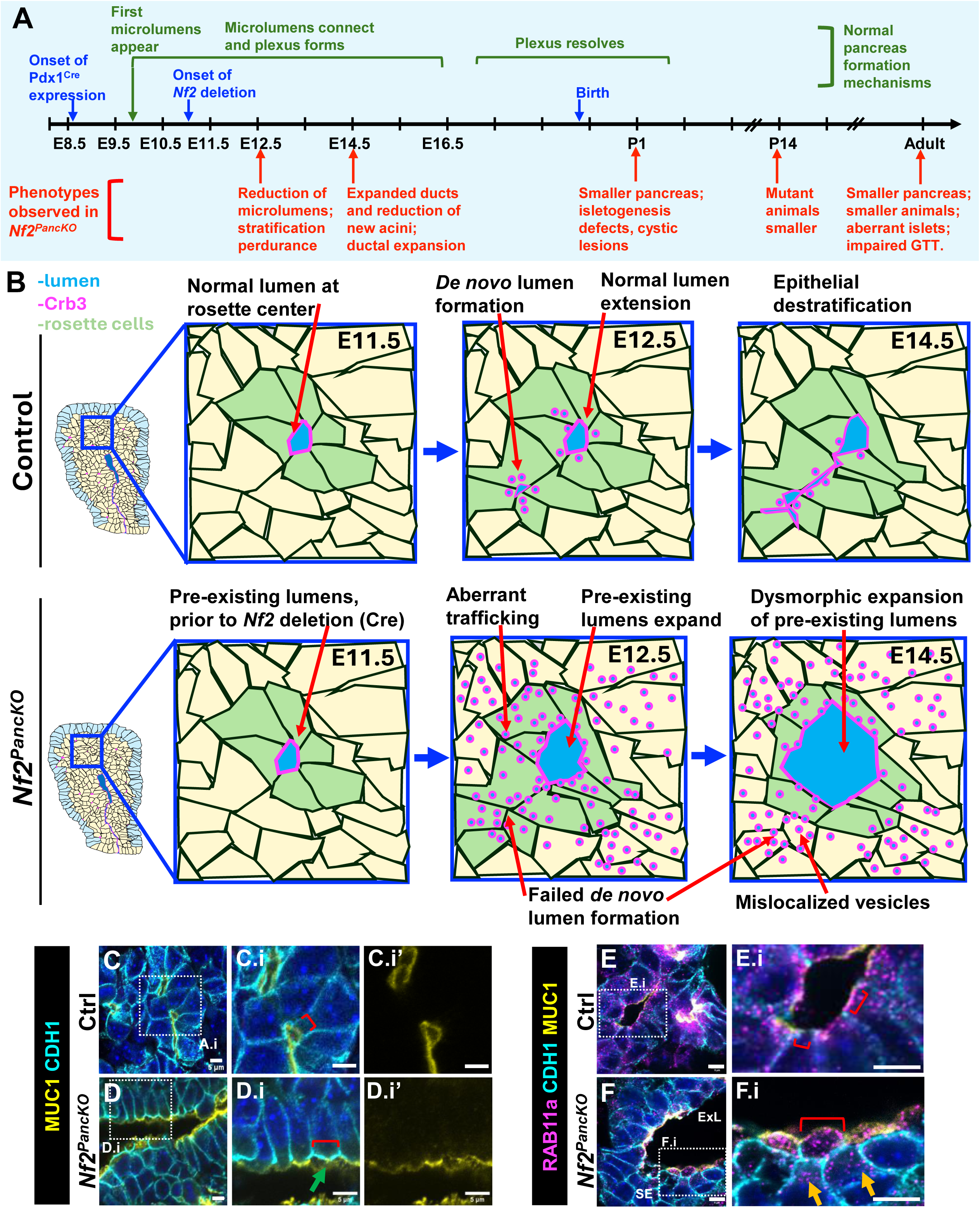
Epithelial phenotype summary: *De novo* lumens fail to form in *Nf2^PancKO^*, while pre-existing lumens expand, likely due to dysregulated vesicular trafficking. **A)** Timeline of phenotypes present in *Nf2^PancKO^*. Pdx1^Cre^ expression begins at approximately E8.5, microlumen formation at approximately E10.0, and MERLIN deletion at approximately E11.5. First morphological defects are evident by E12.5. By E14.5, ductal defects are robust, and acinar numbers are reduced. By P1, pancreata are visibly smaller, and contain cystic lesions. Adult animals display reduced body size and impaired endocrine function. **B)** Models of epithelial processes in both control and mutant pancreata, between E11.5 and E14.5. In both genotypes, select cells have already undergone microlumen formation at E11.5. In controls, *de novo* lumens form, leading to lumen growth and epithelial destratification. In mutant pancreata, cells lining pre-existing lumens maintain polarity and continue to expand; however, cells that have not initiated lumens prior to Merlin deletion fail to assemble an AMIS and remain stratified. Mislocalization of vesicular trafficking components in these cells suggest that Merlin is essential for coordinating membrane trafficking during lumenogenesis. **C-D.i’)** IF for CDH1 and MUC1 in control and mutant E14.5 tissue (15μm sections). In expanded lumens of mutant pancreata, the apical domain (red brackets) often appears expanded, and MUC1 extends into the lumens space along these rounded protrusions. Representative images are shown from >5 embryos per genotype. Rab11 IF on control **(E)** and mutant **(F)** pancreata at E14.5. Note accumulation of RAB11^+^ vesicles at the apical domain (red bracket) and increased cytosolic Rab11^+^ structures (orange arrows). All scale bars are 5μm.

**Extended data 8.**
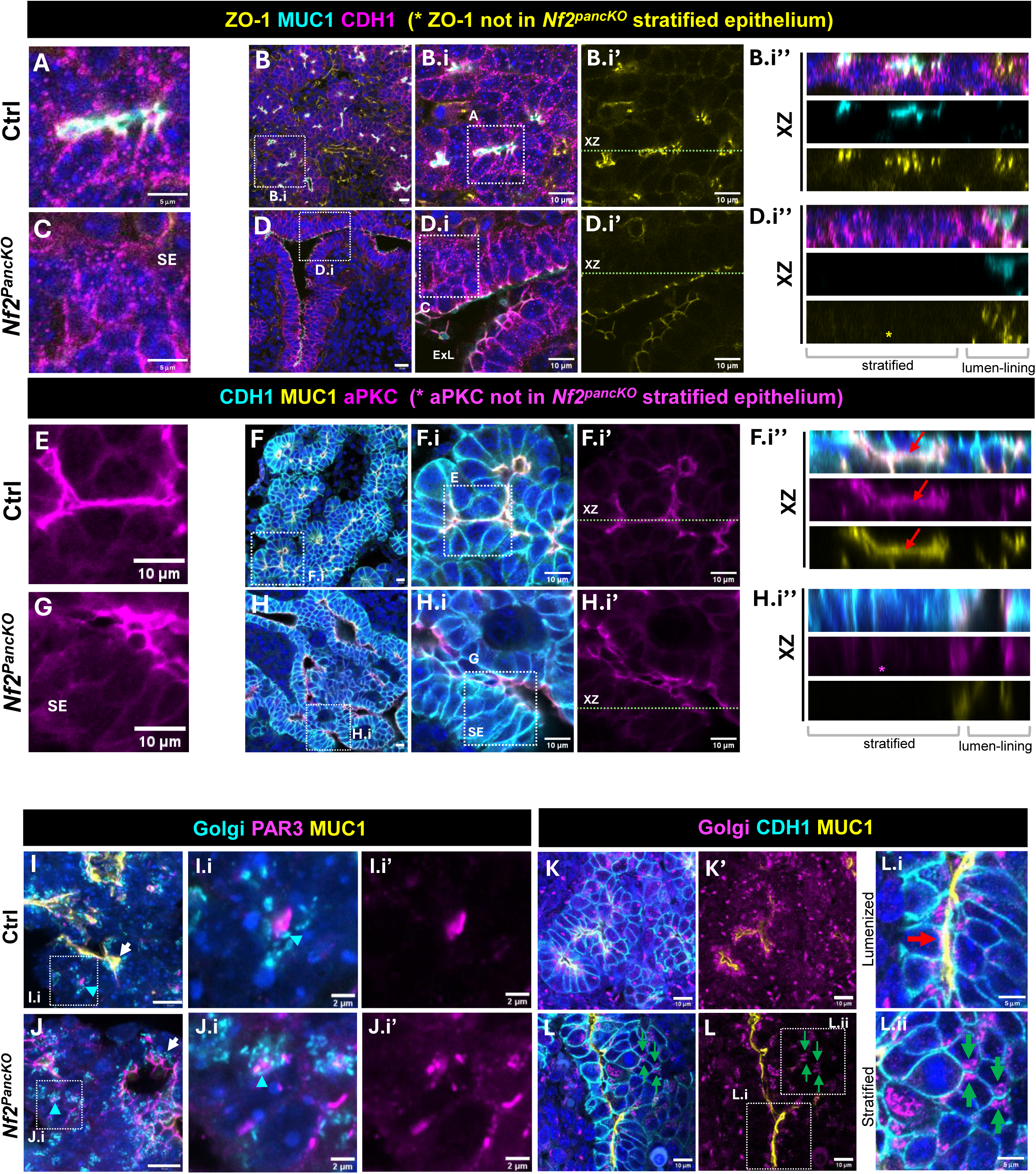
Persistence of partially polarized and apolar cells in E14.5 *Nf2^PancKO^* pancreata. **A-D.i’’)** Immunofluorescence (IF) for ZO-1, MUC1, and CDH1 in control and mutant pancreata. High-magnification views of central pancreatic epithelium in control **(A)** and mutant **(C)** tissue from panels B.i and D.i, respectively. SE indicates stratified epithelium. Dashed box indicates regions shown in B and D, respectively. **B.i-B.i’’)** ZO-1 is restricted to the apico-junctional regions of lumenized epithelium in control tissue. **D-D.i’’)** In mutant tissue, ZO-1 is correctly localized at expanded lumens, but is absent in stratified regions, in both XY plane and orthogonal (XZ) projections through the 10µm stack (**D.i’’).** Dashed green line in B.i’ and D.i’ indicates the planes used for orthogonal reconstructions. **E-H.i’’)** IF for aPKC, CDH1, and MUC1 in E14.5 control and mutant pancreata. aPKC is present at approximately the same levels in control and mutant tissue in lumen-lining cells. **E)** aPKC is tightly localized to the apical membrane in control tissue. **G)** In lumen lining cells of *Nf2^PancKO^*, aPKC is present along the apical membrane. However, in SE regions, aPKC immunoreactivity was undetectable. Dashed green lines in F.i’ and H.i’ indicate plane used for orthogonal projections in **F.i’’** and **H.i’’**. **I-J.i’)** IF for Golgi (GM130) and MUC1. In control pancreata (**I.i)**, the Golgi is localized subapically, under Par3, at both mature lumens (white arrow) and at developing AMIS (cyan arrowhead). In mutant AMIS **(J.i)**, Golgi positioning is maintained (cyan arrowhead), but no longer aligns with PAR3 localization at the AMIS. Golgi positions normally at expanded lumens (white arrow). **K-L.ii)** IF for Golgi (GM130), CDH1, and MUC1. **K, K’)** In control pancreata, Golgi are localized to the subapical regions of lumen-lining epithelial cells (red arrow). **(L, L.i)** In *Nf2^PancKO^*, lumen-lining cells display subapical Golgi, while stratified epithelial cells frequently oriend paired Golgi towards one another **(L.ii)**, indicating persistence of a polarized intracellular axis despite failure to establish an apical domain. For A-L, representative images from N ≥ 3 embryos evaluated per genotype. Scale bars are 10μm in all panels, except 5μm in A, C, L.i, L.ii and 2μm in I.i, I.i’, J.i, J.i’.

**Extended data 9.**
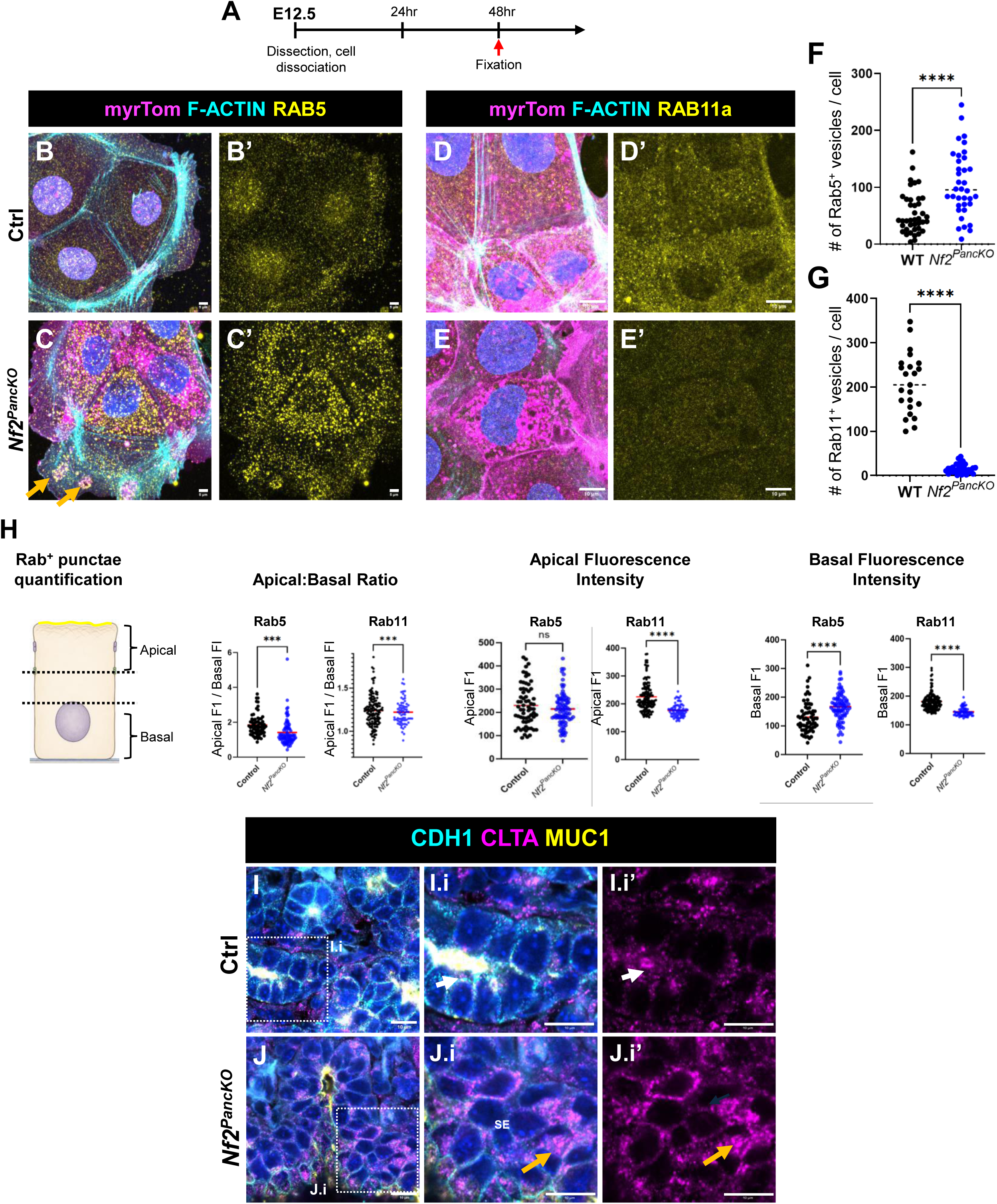
Loss of MERLIN disrupts the balance and spatial organization of RAB5^+^ and RAB11^+^ vesicle populations. **A)** Experimental schematic for dissociated epithelial cell analysis. E12.5 pancreata were dissociated into single cells, cultured for 48 h, fixed, and processed for IF. **B-C’)** IF for RAB5 in cultured control and *Nf2^PancKO^* cells. Mutant cells exhibit increased numbers of RAB5^+^ vesicles. Many intracellular myrTOM^+^ structures are RAB5^+^ (orange arrows). **D-E’)** IF for RAB11 in cultured control and *Nf2^PancKO^* cells. RAB11^+^ vesicles are reduced in *Nf2^PancKO^* cells. Quantification of number of Rab5^+^ **(F)** and RAB11^+^ **(G)** intracellular structures shows an overall increase of Rab5 ^+^ and decrease of Rab11 ^+^ vesicles. (For J, n= 40 control, and 36 mutant cells evaluated. P <0.0001 by Welch’s t.test. For K, n=23 control and 35 mutant cells evaluated. P <0.0001 by Welch’s t.test). **H)** Quantification of Rab5/Rab11 localization in lumen-lining MUC1^+^ epithelial cells at E14.5. Cells were divided into apical and basal compartments, and fluorescence intensity (FI) was measured in each region. Note alteration of RAB5^+^ and RAB11^+^ vesicle distribution, with shift from Rab11-mediated recycling towards Rab5-endosomal compartments. (Mean gray value was utilized as a measure of FI for each portion of the cell. Data analyzed by Welch’s t-test. P values = Rab5 Apical FI/Basal FI – 0.0001, Rab11 Apical FI/Basal FI – 0.0001, Rab5 Apical FI – 0.27, Rab11 Apical FI -- <0.0001, Rab5 Basal FI -- <0.0001, Rab11 Basal FI -- <0.0001). **I-J.i’)** IF for clathrin heavy chain (CLTA) on control (I-I.i’) and mutant (J-J.i’) pancreata at E14.5. I) Normally, CTLA accumulated at forming lumens that express MUC1. J) In mutant epithelia, CTLA is ectopically localized in the cytoplasm (orange arrows), consistent with disrupted membrane trafficking. Scale bars: Panels B-C’ 5μm, D-E’ and I –J.i’ 10μm. For all graphs, midline represents median.

**Extended data 10.**
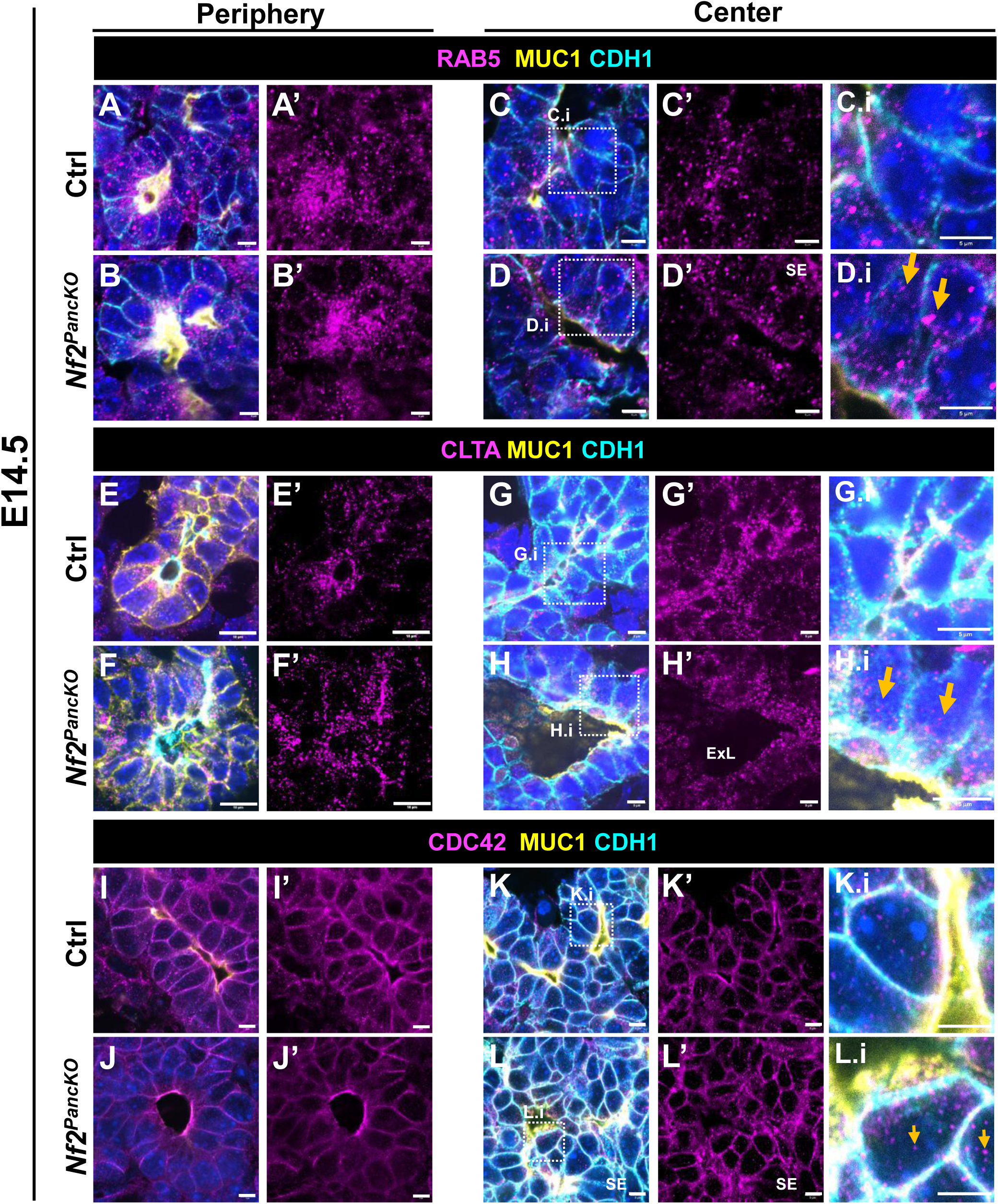
Loss of MERLIN preferentially disrupts trafficking protein localization in the core stratified epithelium. **A-L.i’)** IF for trafficking components in E14.5 control and *Nf2^PancKO^* pancreata. Peripheral (A-B’, E-F’, J-K’) and central (C-D.i, G-H.i, L-M.i) epithelial regions are shown. **A-D.i)** IF for RAB5 in E14.5 control (A, C) and *Nf2^PancKO^* (B, D) pancreata. **C.i, D.i)** Dotted lines indicate region of high magnification views. RAB5 is enriched along the MUC1^+^ apical membrane, particularly in peripheral epithelium of both control and mutant cells, but accumulates in ectopic cytoplasmic punctae in the core epithelium of mutants (orange arrows). **E-H.i)** IF for CTLA in E14.5 control (E, G) and *Nf2^PancKO^* (F, H) pancreata. **G.i, H.i)** High magnification views. CTLA is apically localized in both control and mutant tissue, but shows increased cytoplasmic localization in mutant cells (orange arrows). **I-L.i)** IF for CDC42 in E14.5 control (I, K) and *Nf2^PancKO^* (J, L) pancreata. **K.i, L.i)** High magnification views. CDC42 is enriched along the MUC1^+^ apical membrane, in both controls and mutant cells, exhibits increased cytoplasmic localization in core epithelial mutant cells. In all three cases, trafficking defects are more pronounced in the central (core) region of the pancreas. MERLIN-dependent trafficking defects first appear in the epithelial population that fails to form new lumens following Cre-mediated deletion. Representative images from N ≥ 2 embryos evaluated per genotype. Scale bars: Panels A-D.i, G-L.i’ 5μm, and E-F’ 10μm.

**Extended data 11.**
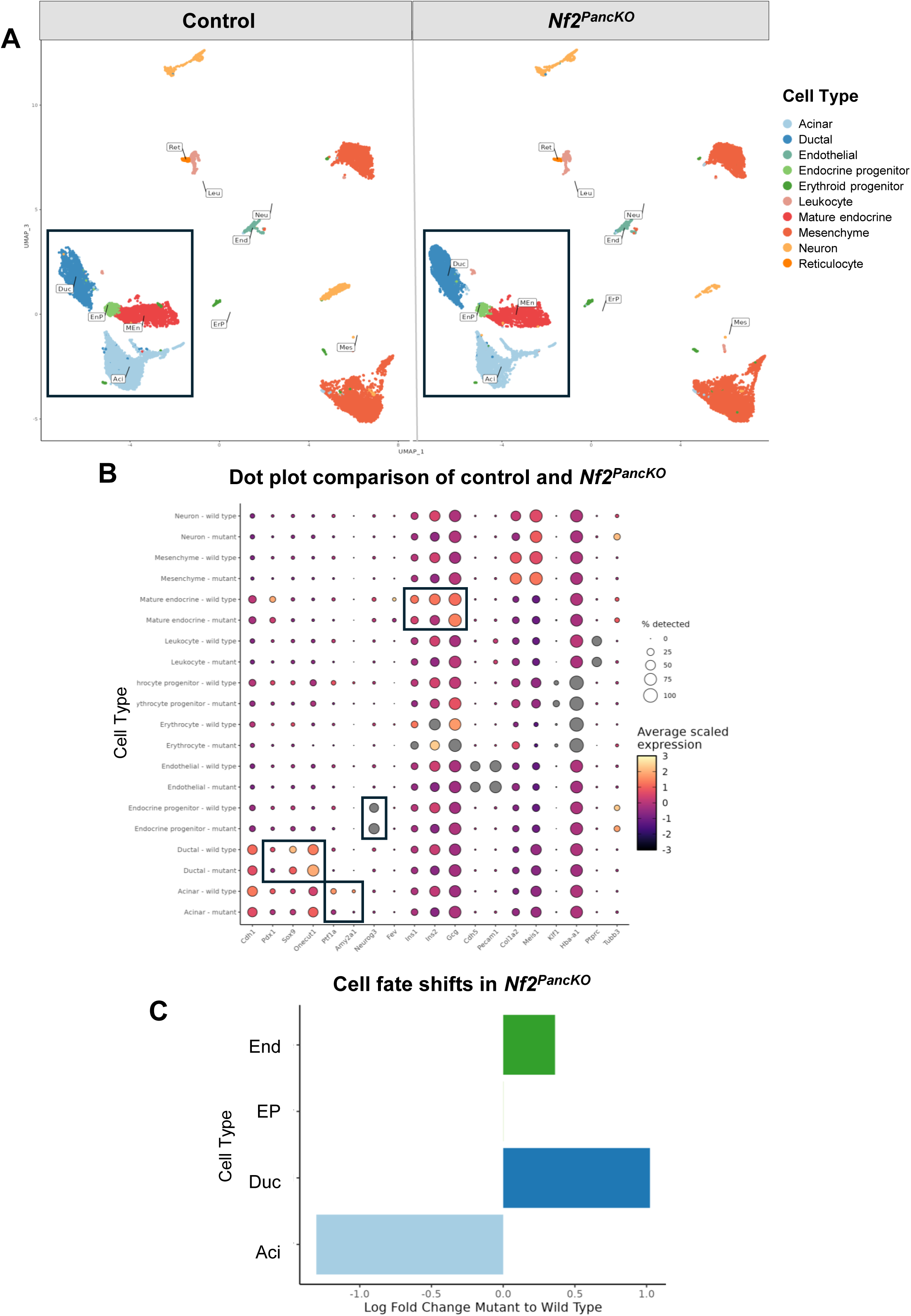
Loss of MERLIN alters lineage composition at E14.5. Transcriptomic comparison of control and *Nf2^PancKO^* epithelium at E14.5, using only E14.5 dataset. **A)** UMAP representation of single nuclear sequencing (snRNA-seq) from E14.5 control and mutant pancreata. Boxes highlight the major epithelial populations. A total of 11,511 epithelial nuclei were analyzed from control pancreata, and 8,020 epithelial nuclei were obtained from *Nf2^PancKO^*. Acinar, ductal, endocrine progenitor, and endocrine clusters were identified. **B)** Canonical marker genes were used to identify epithelial cell populations and for cluster annotations. Black boxes denote markers of interest. **C)** Relative abundance of epithelial cell populations in control and mutant pancreata. Loss of MERLIN results in expansion of ductal cells, a modest increase in endocrine cells, and a corresponding reduction in acinar cells. Control epithelial nuclei contained: acini –49%, ductal --26% endocrine progenitor – 14%, endocrine – 7%; whereas *Nf2^PancKO^* epithelial nuclei contained: acini –20.4%, ductal –50.2% endocrine progenitor – 14%, endocrine – 12%.

**Extended data 12.**
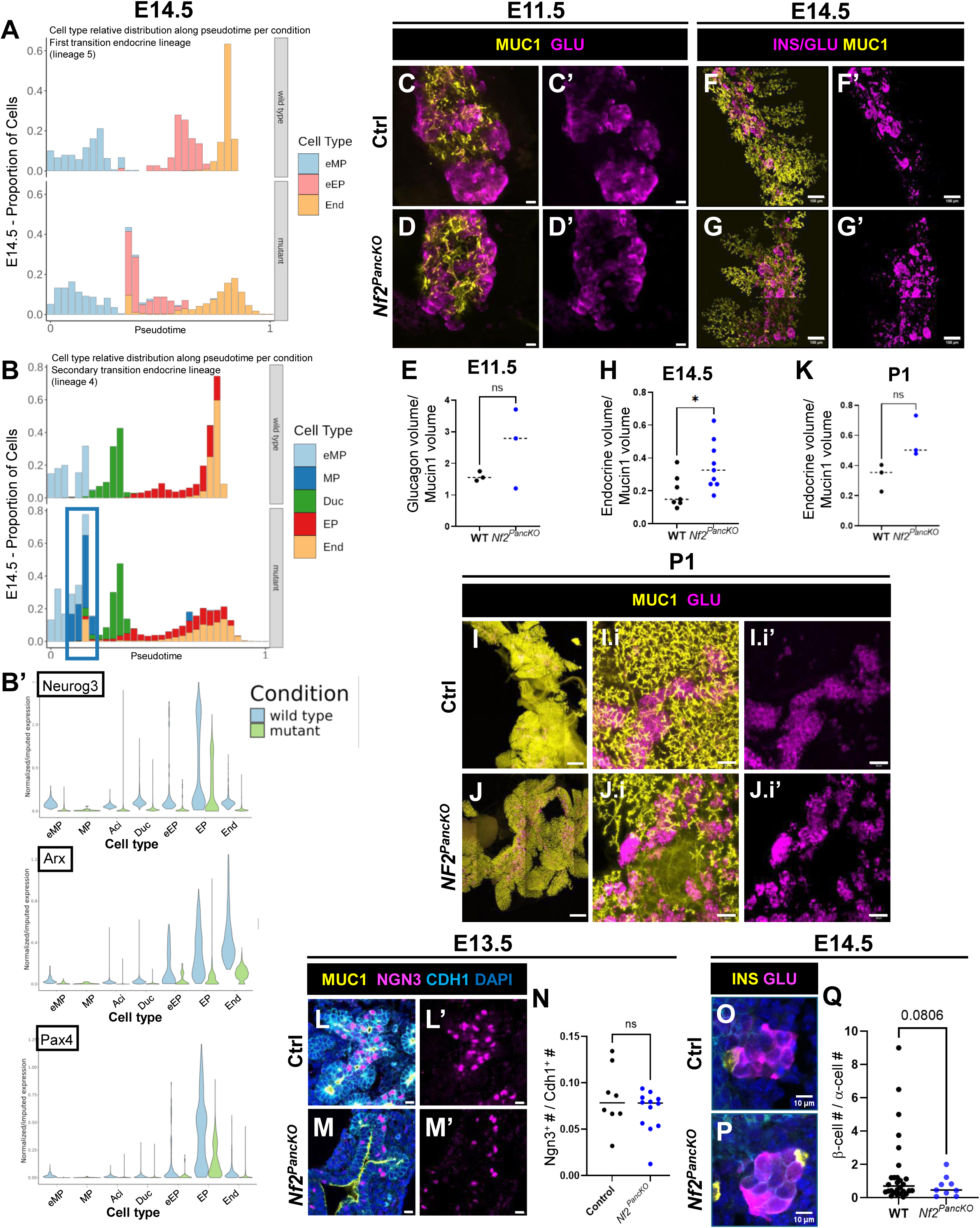
Endocrine differentiation is delayed in *Nf2^PancKO^* pancreata, resulting in reduced postnatal endocrine mass. Endocrine mass transiently increases in *Nf2^PancKO^*, and subsequent isletogenesis is disrupted. **A, B)** Pseudotime distributions (histograms) of cells progressing along the early (A) and late (B) endocrine differentiation trajectories reconstructed from combined E11.5 and E14.5 single nucleus RNA-seq datasets. Mutant pancreas exhibit delayed progression along both endocrine trajectories. **B’)** Expression of key endocrine differentiation markers (*Neurog3, Arx, and Pax4*) in from E14.5 data using clustering from combined E11.5 and E14.5 datasets. **C-J.i’)** WMIF of control **(C, F, I)** and *Nf2^PancKO^* **(D, G, J)** pancreata at E11.5 **(C, D)**, E14.5 **(F, G)**, and P1 **(I-J.i**’) for MUC1 and GLU **(C, D)**, INS/GLU and MUC1 **(F, G)** and MUC1 and GLU **(I, J)**. E11.5 pancreata is stained for MUC1 and glucagon only, for other time points pancreata are stained for MUC1 and combined antibodies against insulin and glucagon. **E,H,K)** Quantification of endocrine volume at E11.5 **(E)**, E14.5 **(H)**, and P1 **(K)**, respectively. Each dot represents an individual animal analyzed. (P= 0.25, 0.02, 0.06 by unpaired t.test, respectively). Section IF of E13.5 control **(L, L’)** and *Nf2^PancKO^* **(M, M’)** for MUC1, NGN3, and CDH1 at E13.5. **N)** Quantification of NGN3+ cells shows no significant difference between genotypes (p=0.26 n=3 ctrl, 3 mutant animals analyzed). **O-Q)** Section IF for insulin and glucagon in E14.5 control (O) and mutant (P) pancreata. **Q)** Quantification of the ratio of β-cells:α-cells at E14.5 shows no significant difference between genotypes (p=0.08 by Welch’ t-test, n=4 mutants, 6 ctrl animals evaluated). Each dot represents one FOV. Scale bars: Panels C-D’ 10μm; F-G’ 100μm; I-J 250 μm; I.i-J.i’ 50μm’ L-M’ 10μm, O, P 10μm. For all graphs, midline indicates median.

**Extended data 13.**
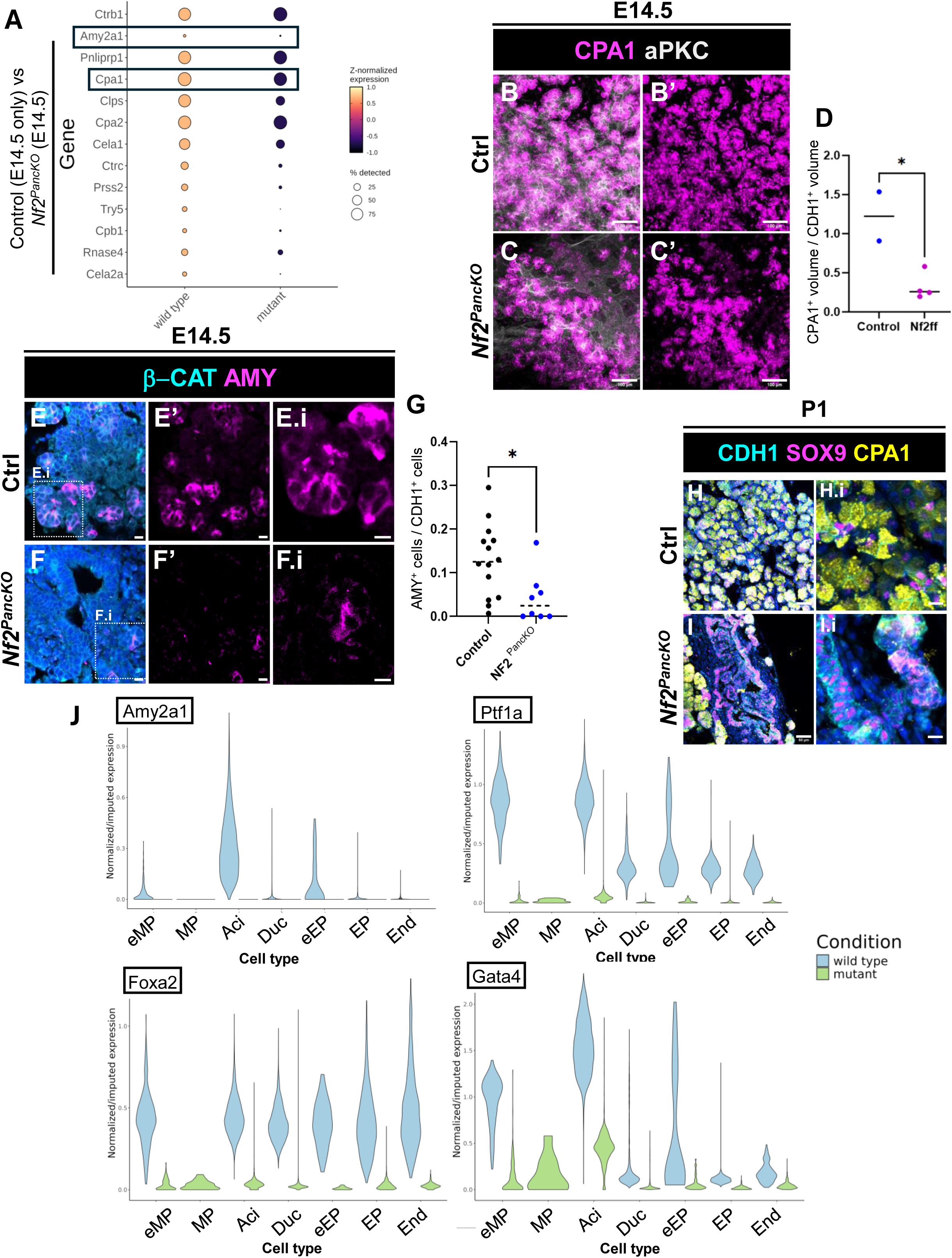
Acinar cells are poorly differentiated in *Nf2^Panc^^KO^*. **A)** Dot plot showing reduced expression of key acinar secretory genes from E14.5 mutant acinar nuclei relative to E14.5 controls, using the clustering from the integrated single nucleus dataset. WMIF of E14.5 control **(B, B’)** and *Nf2^PancKO^* **(C, C’)** pancreata stained for aPKC and CPA1. **D)** Quantification of CPA1^+^ volume. Each dot represents one pancreas, p = 0.018 by unpaired t-test. Section IF of E14.5 control **(E-E.i)** and *Nf2^PancKO^* **(F-F.i)** stained for β-catenin (β-CAT) and amylase (AMY). White boxes indicate region magnified in **E.i-F.i. G)** Quantification of AMY^+^ epithelial cells per FOV. Each dot represents a single FOV analyzed, from n = 3 ctrl and mutant pancreata each. AMY^+^ epithelial cells are reduced in *Nf2^PancKO^*. P =0.0044 by unpaired t-test. **H-I.i)** IF of control (H) and mutant (I) pancreata for CDH1, SOX9, and CPA1 at P1. **J)** Violin plots of *Amy2a*, *Ptf1a*, *Foxa2*, and *Gata4* from integrated dataset, comparing E14.5 control and mutant nuclei. Scale bars: Panels B-C’ 100μm; E-F.i, H.i, I.i 10μm; and H,I 50μm. For D-G, midline indicates median.

**Extended data 14.**
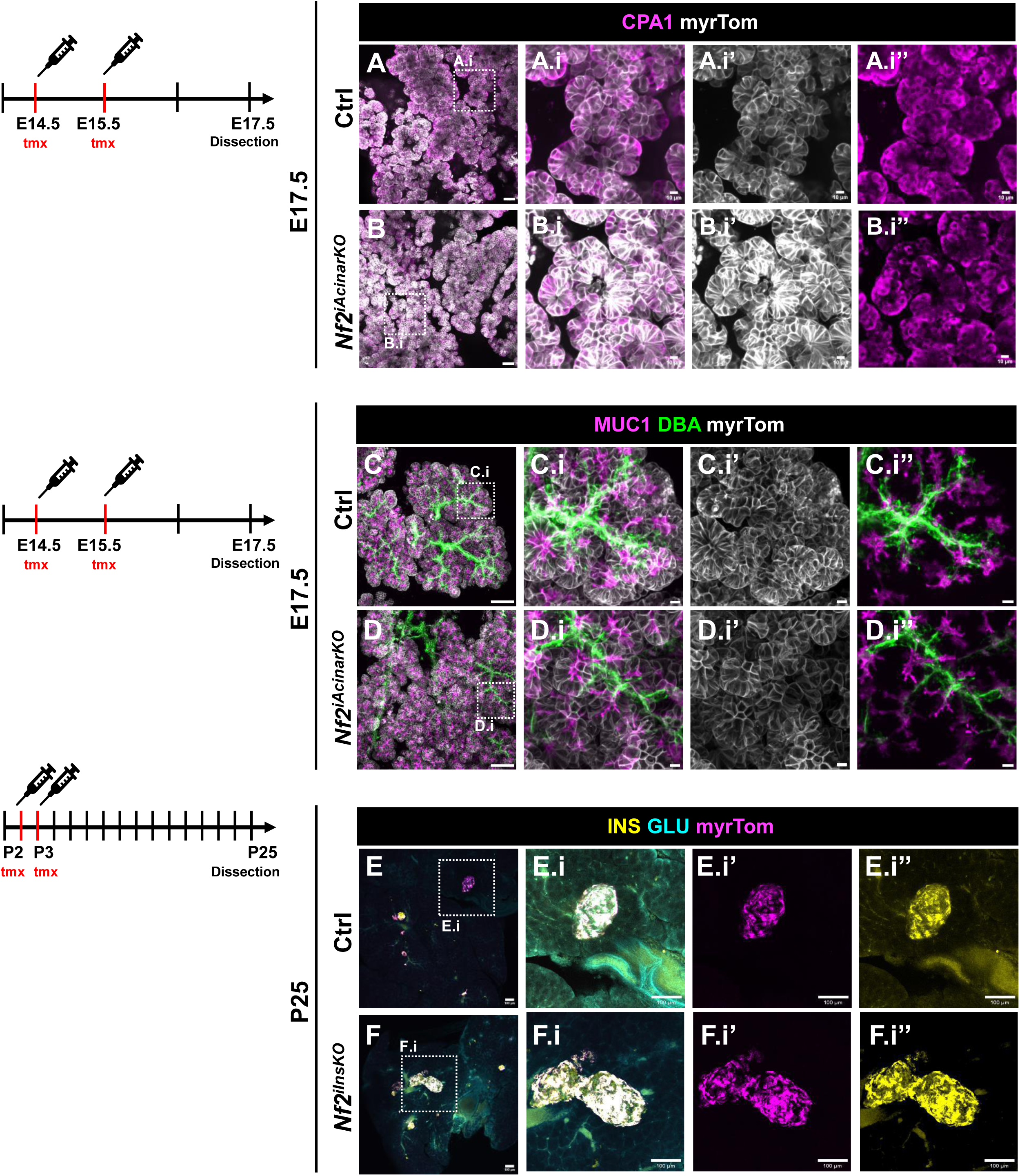
MERLIN is required during pancreatic lineage establishment but not lineage maintenance. **A-B.i’’)** Representative 200μm vibratome section of E17.5 control (A) and *Nf2^iAcinarKO^* (B) pancreata stained for CPA1 and myrTom. Robust myrTom expression confirms efficient Cre-mediated recombination following tamoxifen (tmx) administration at E14.5 and E15.5. Dotted lines indicate region of high magnification views in A.i and B.i. CPA1 expression was normal in mutant acinar cells. **C-D.i’)** Representative vibratome sections of E17.5 control (C) and *Nf2^iAcinarKO^* (D) pancreata stained for MUC1 and DBA, demonstrating normal lumenal morphology following post-morphogenetic deletion of MERLIN. **E-F.i’’)** Representative section IF of P25 control (E) and *Nf2^iInsKO^* (F) pancreata following tmx induction at P2 and P3. *Nf2^iInsKO^* animals display robust insulin expression and normal islet architecture following postnatal deletion of MERLIN in differentiated β-cells. These data show that MERLIN is required during a narrow developmental window but not for maintenance of differentiated fates. Representative images from n ≥2 animals per genotype. Scale bars: Panels A, B 50μm, A.i-B.i’’ 10μm, C, D 100μm, C.i-D.i’’ 10μm, E-F.i’’ 100μm.

**Extended data. 15.**
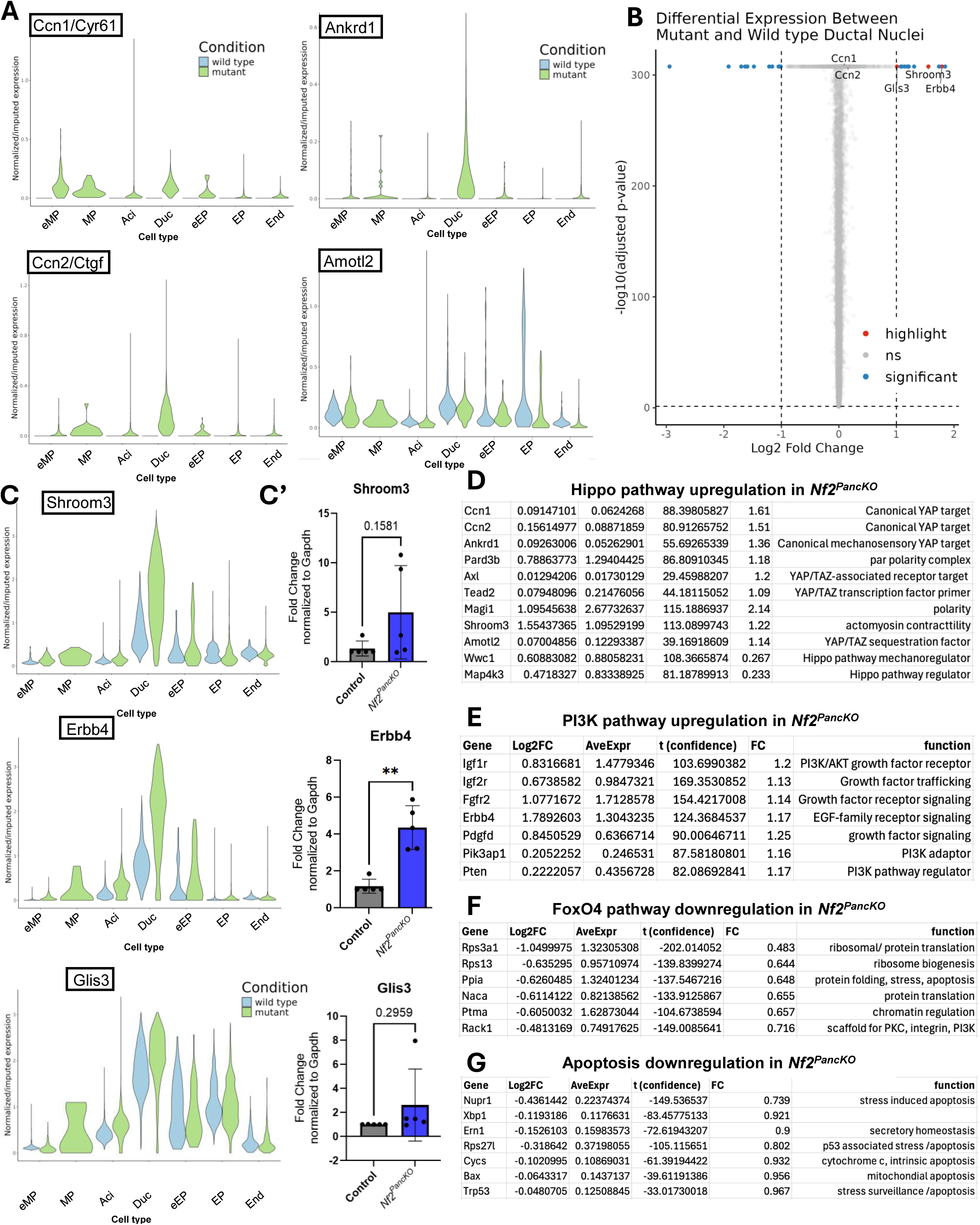
Loss of MERLIN alters Hippo- and PI3K-associated transcriptional programs in E14.5 pancreatic epithelium. **A)** Violin plots showing upregulation of canonical Hippo pathway target genes, *cyr61, ankrd1, ctgf, and amotl2* in E14.5 in *Nf2^PancKO^* pancreata. Violin plots made from integrated dataset (with only E14.5 control and mutant nuclei compared). **B-C)** Ductal nuclei from integrated dataset show significant upregulation of *glis3*, *shroom3*, and *erbb4*. **C’)** qPCR validation of *shroom3*, *erbb4*, and *gli3s* in E14.5 control and mutant pancreata. Each dot represents one pancreas analyzed. Statistics by Welch’s t-test, for Erbb4 p=0.002. Error bars indicate standard deviation. **D-G)** Gene set enrichment analysis (GSEA) of ductal nuclei shows activation of Hippo and PI3K-associated programs and repression of FoxO4 and apoptosis-associated programs in in *Nf2^PancKO^* pancreata, as per the E14.5-only dataset (clustering as in Fig. S11A).

**Extended data 16.**
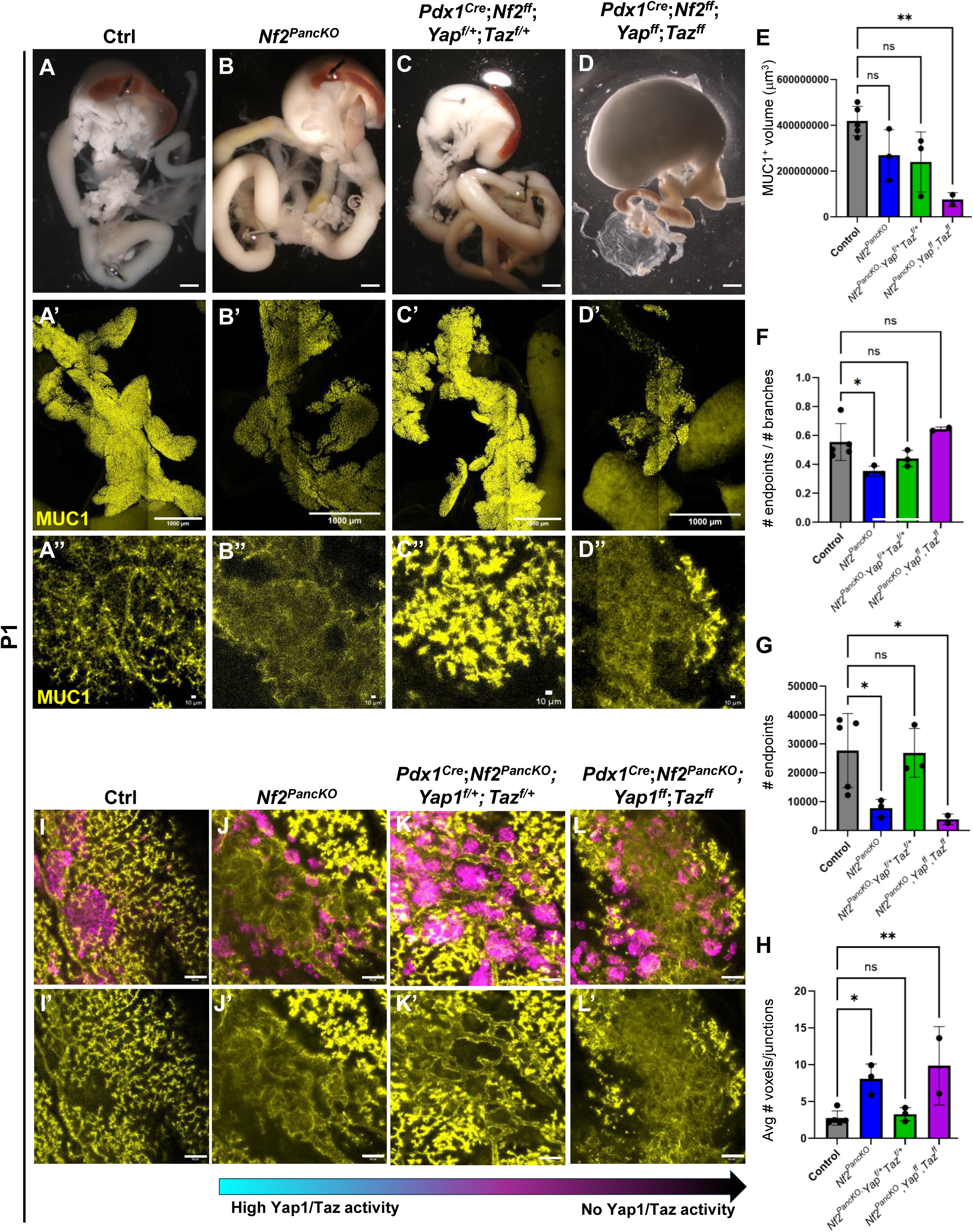
Reduced YAP/TAZ dosage partially rescues pancreatic morphogenesis in *Nf2^PancKO^* mice. **A-D)** Gross morphology of digestive tracts from control (A), *Nf2^PancKO^* (B), *Nf2^PancKO^Yap/Taz ^Haploinsufficiency^* (C), and *Nf2^PancKO^Yap/Taz^PancKO^* (D) mice at P1. **A’-D’’)** WMIF of MUC1 showing the pancreatic ductal network at P1 for each genotype, as indicated (A’-D’, lower magnification. A’’-D’’, higher magnification). **E-H)** Quantification of various aspects of ductal lumen morphology. Each dot represents one pancreas evaluated. Data was analyzed by one-way ANOVA, followed by Dunnet’s test for multiple comparisons. **E)** Total MUC1^+^ volume per pancreas. (P values: control vs *Nf2^PancKO^* – 0.13, control vs *Nf2^PancKO^Yap/Taz ^Haploinsufficiency^* _–_ 0.07, control vs *Nf2^PancKO^Yap/Taz^PancKO^* _–_ 0.005). **F)** Number of terminal endpoints/total branches normalized to total branch number. (P values: control vs *Nf2^PancKO^* – 0.03, control vs *Nf2^PancKO^Yap/Taz ^Haploinsufficiency^* _–_ 0.28, control vs *Nf2^PancKO^Yap/Taz^PancKO^* _–_ 0.55). **G)** Total number of terminal endpoints/pancreas. (P values: control vs *Nf2^PancKO^* – 0.049, control vs *Nf2^PancKO^Yap/Taz ^Haploinsufficiency^* _–_ 0.99, control vs *Nf2^PancKO^Yap/Taz^PancKO^* _–_ 0.04). **H)** Mean junction size. (P values: control vs *Nf2^PancKO^* – 0.02, control vs *Nf2^PancKO^Yap/Taz ^Haploinsufficiency^* _–_ 0.97, control vs *Nf2^PancKO^Yap/Taz^PancKO^* _–_ 0.0096). Reduction of YAP/TAZ dosage partially restores ductal architecture in mutant pancreata. **I-L)’** Higher magnification images of the central epithelium from each genotype stained for MUC1 together with insulin (INS) and glucagon (GLU). Scale bars: Panels A-D 500μm, A’-D’ 1000μm, A’’-D’’ 10μm, I-L’ 50μm. For all graphs, error bars indicate standard deviation.

**Extended data 17.**
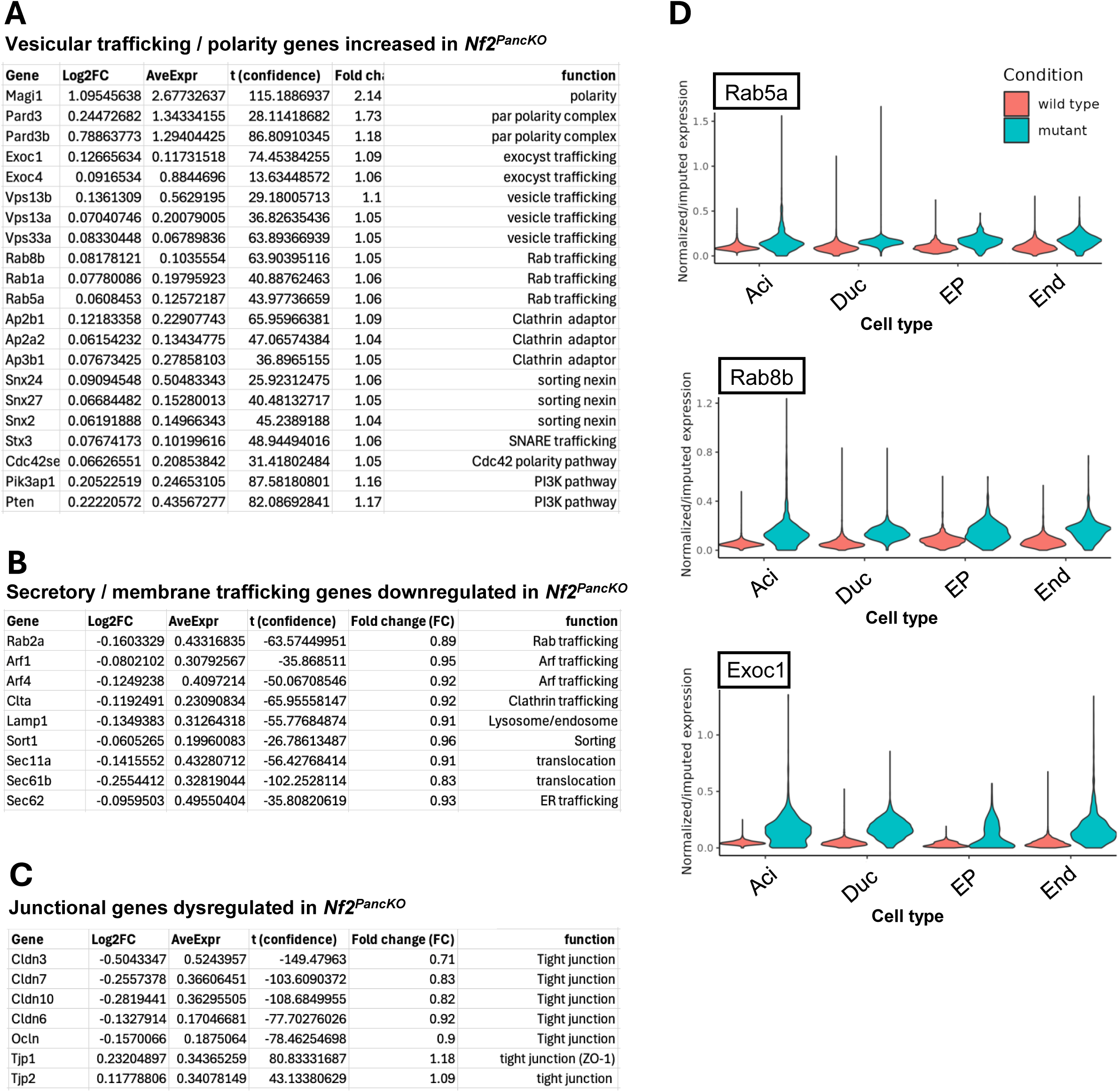
Loss of MERLIN alters transcriptional programs regulating polarized membrane trafficking. **A-C)** Key trafficking, secretory, and junctional genes are dysregulated in *Nf2^PancKO^*, in E14.5 ductal nuclei (clustering as in Fig. S10). **D)** Violin plots showing expression of selected vesicle trafficking genes in E14.5 control and mutant ductal nuclei (E14.5-only data set, clustering as in Fig S11A).

**Extended data 18.**
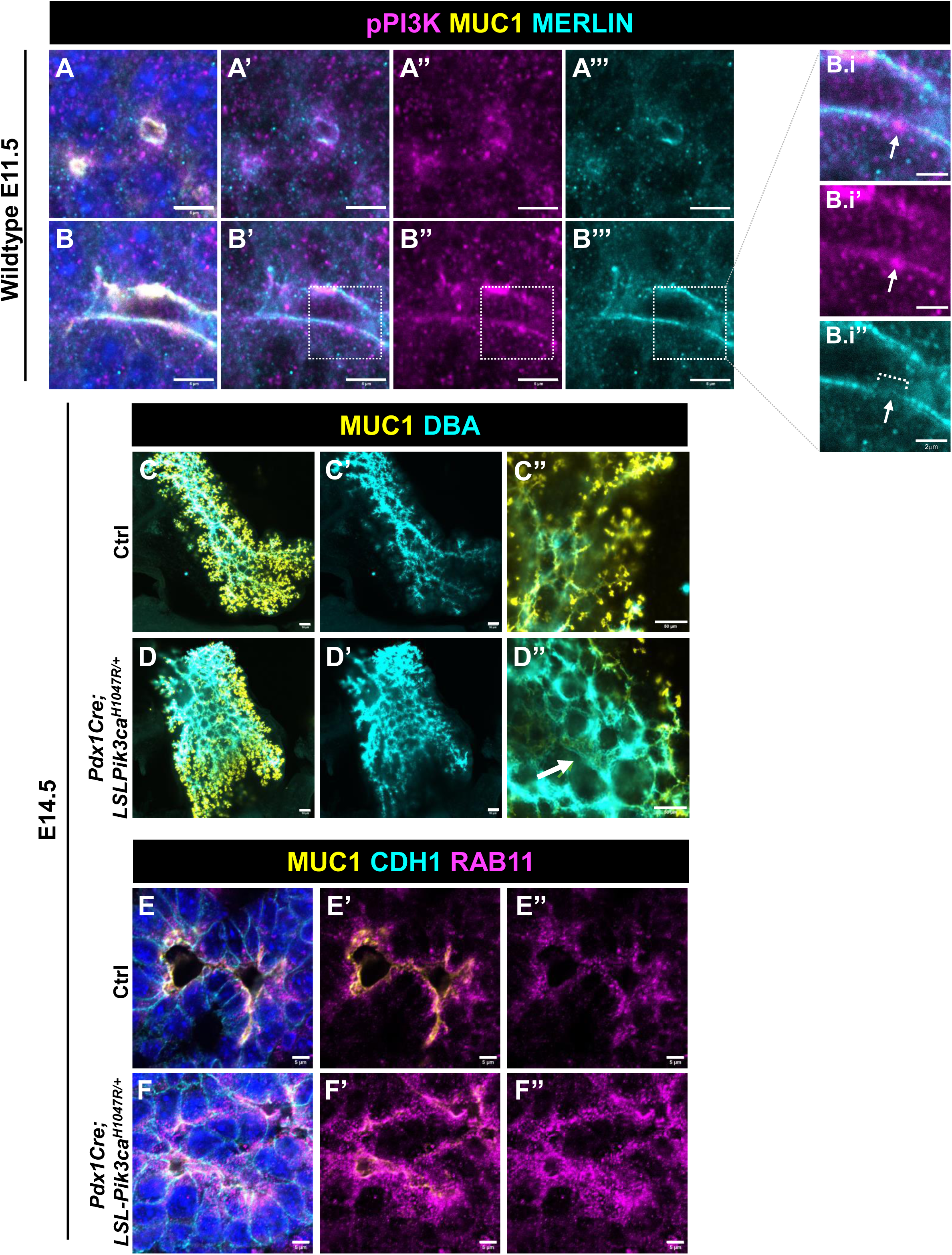
Genetic hyperactivation of PI3K signaling phenocopies the epithelial remodeling defects of *Nf2^PancKO^* pancreata. **A-B’’’)** IF for phosphorylated PI3K (antibody against phospho-p85/p55), MERLIN, and MUC1 in wildtype E11.5 pancreata. Both pPI3K and MERLIN are apically localized at **(A-A’’’)** nascent and **(B-B’’’)** open/mature lumens. **B.i-B.i’’)** High magnification insets of open lumen in B-B’’’. White arrow indicates region of high pPI3K immunoreactivity, and relatively low MERLIN signal. **C-D’)** WMIF for DBA and MUC1 showing MIPs of 100μm substacks through the central region of control **(C)** and *Pdx1^Cre^;LSL-Pik3ca^H1047R/+^* **(D)** pancreata. **C’’, D’’)** Optical section at high magnification. Constitutive PI3K activation results in enlarged, dysmorphic ducts (white arrows), closely phenocopying the epithelial defects observed in mutants. Representative images from n=2 embryos. **E-F’’)** IF for RAB11, CDH1, AND MUC1 in E14.5 control (E) and *Pdx1^Cre^;LSL-Pik3ca^H1047R/+^* (F) pancreata. Constitutive PI3K activation disrupts normal RAB11 localization, consistent with impaired polarized membrane trafficking. Scale bars: Panels A-B’’’ 5μm, B.i-B.i’’ 2μm, C-D’’ 50μm, E-F’’ 5μm.

**Supplementary Fig. 19.**
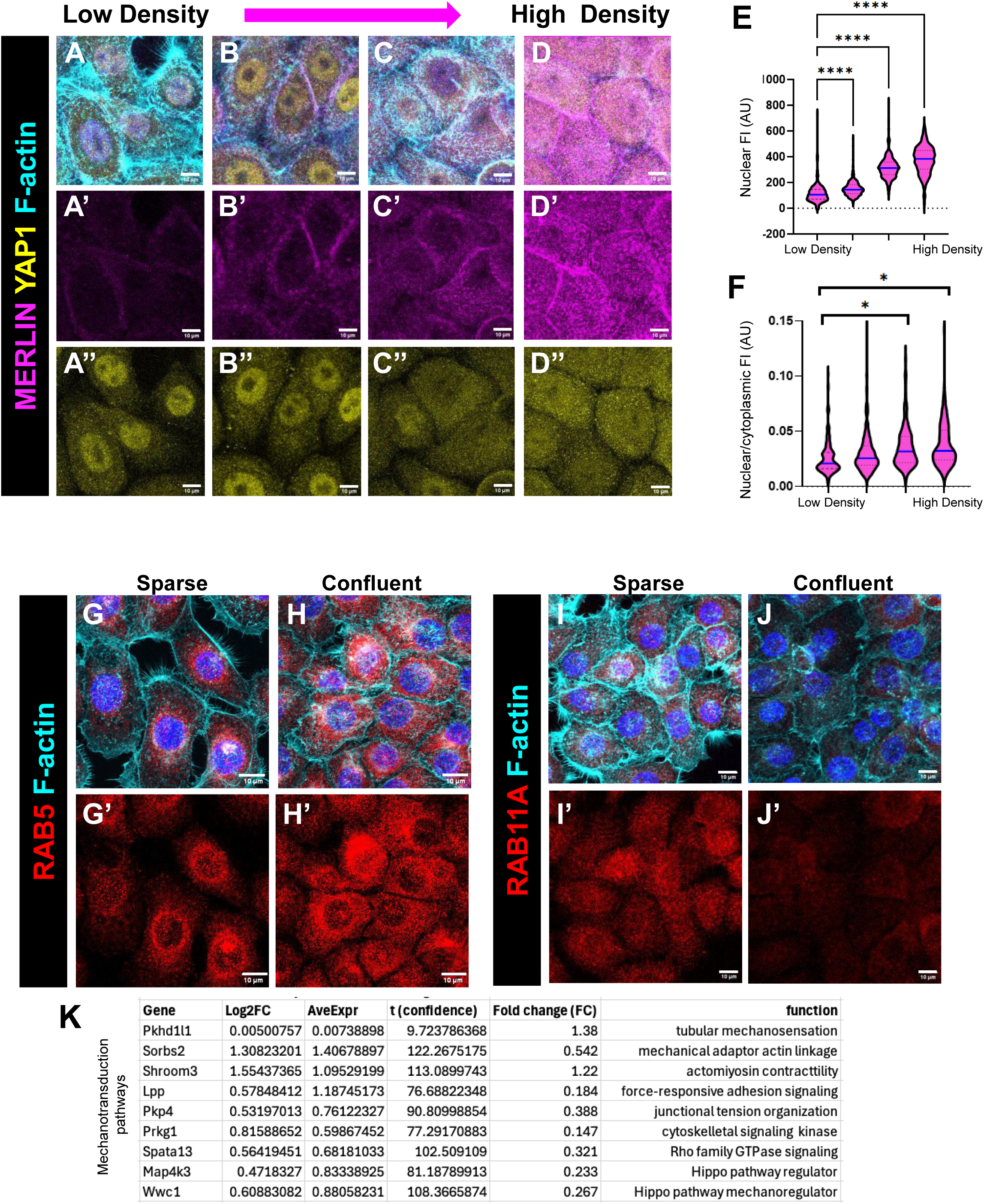
MERLIN and mechanotransduction-associated pathways respond to epithelial cell density. **A-D)** Human pancreatic ductal epithelial (HDPE) cells cultured at 10,000 cells/cm^2^ (A), 20,000 cells/cm^2^ (B), 30,000 cells/cm^2^ (C), or 50,000cells/cm^2^ (D) and stained for MERLIN, YAP1, and F-actin. Increasing cell density (increased intracellular tension) results in elevated nuclear MERLIN localization. **E)** Quantification of nuclear MERLIN fluorescence intensity (FI). (Data analyzed by Anova followed by Dunnett’s test for multiple comparisons. P <0.001 for all comparisons). **F)** Quantification of nuclear/cytoplasmic MERLIN fluorescence ratio. (Data analyzed by Anova followed by Dunnett’s test for multiple comparisons. P = 0.10 for seeding density of 10,000cells/cm^2^ vs 20,000cells/cm^2^, p = 0.0002 for 10,000cells/cm^2^ vs 30,000cells/cm^2^, p <0.0001 for 10,000cells/cm^2^ vs 50,000cells/cm^2^. For E-F, n >100 cells analyzed per condition). For E-F, blue line indicated median of data. **G, H)** IF for RAB5 in sparse (G) or confluent (H) HDPE cultures. **I)** Quantification of MERLIN expression FI in G, H. **J-K’)** IF for RAB5 in sparse (J) or confluent (K) HDPE cultures. **L-M’)** IF for RAB11 in sparse (G) or confluent (H) HDPE cultures. Cell density alters intracellular localization of both RAB5 and RAB11, consistent with density-dependent regulation of membrane trafficking. **N)** Selected mechanotransduction-associated genes are differentially expressed in E14.5 *Nf2^PancKO^* ductal nuclei relative to controls (clustering as in Fig S11). All scale bars 10μm.

## SUPPLEMENTARY INFORMATION

**Supplementary Fig. 1.**
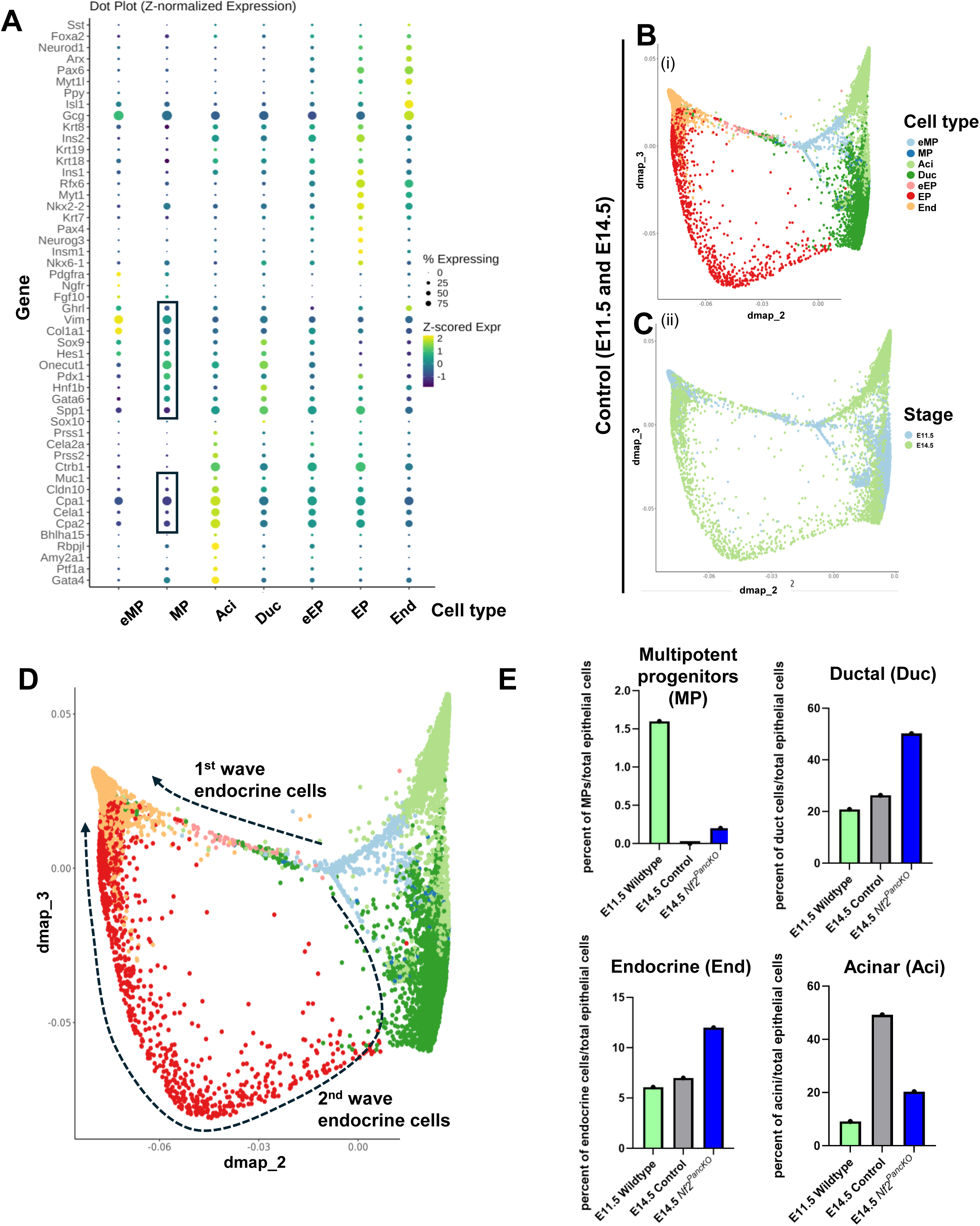
Transcriptomic analysis of control embryonic pancreas using integrated datasets from E11.5 and E14.5. **A)** Markers used to assign cluster IDs are of integrated E11.5 and E14.5 data. **B)** Diffusion map of combined control E11.5 and E14.5 data. **C)** E11.5 cells are shown in light blue, and E14.5 cells are shown in green. **D)** 1^st^ and 2^nd^ wave endocrine cells are annotated. First wave endocrine cells are on the arm which is composed of E11.5 and E14.5 cells, while the second wave endocrine cells are on the arm which is predominately E14.5 cells. E) Percentage of epithelial cells assigned to each selected cell type across conditions.

**Supplementary Fig. 2.**
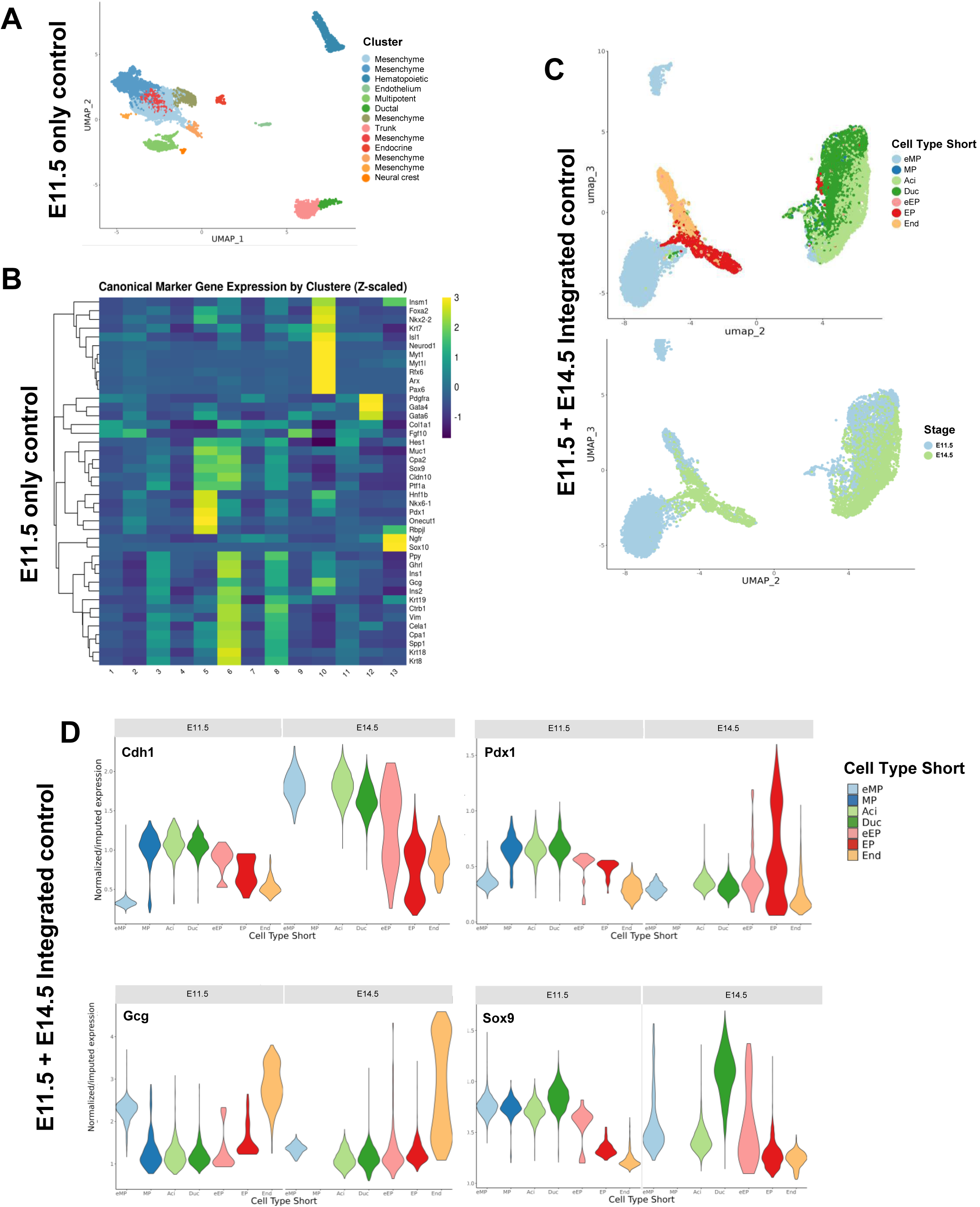
Integration of E11.5 and E14.5 data sets. **A)** UMAP of E11.5 wildtype pancreata –12,834 epithelial cells from E11.5 pancreata were obtained. **B)** Markers used to assign cluster IDs are of integrated E11.5 data **C)** Umap of combined control E11.5 and E14.5 data. **C’)** E11.5 cells are shown in light blue, and E14.5 cells are shown in green. **D)** Expression of key marker genes in integrated data set, with clusters split by stage.

## Supplemental Movie Legends

**Supplemental Movie 1:** Control E14.5 pancreata, stained with CDH1 (cyan), MUC1 (yellow), and laminin (magenta). Movie shows sequential optical sections from z-stack acquired on a light sheet microscope.

**Supplemental Movie 2:** *Nf2^PancKO^* E14.5 pancreata, stained with CDH1 (cyan), MUC1 (yellow), and laminin (magenta). Movie shows sequential optical sections from z-stack acquired on a light sheet microscope.

**Supplemental Movie 3:** Projection of control E14.5 pancreata, stained with CDH1 (gray), MUC1 (blue), and laminin (green). Movie begins on a single optical section through the light sheet, red outline indicates a cell as identified by the IMARIS cell function. Cell is in direct contact with MUC1+ lumen.

**Supplemental Movie 4:** Projection of *Nf2^PancKO^* E14.5 pancreata, stained with CDH1 (gray), MUC1 (blue), and laminin (green). Movie begins on a single optical section through the light sheet, green and purple outline indicates cells as identified by the IMARIS cell function. Green cell does not in contact a lumen

**Supplemental Movie 5:** Live imaging of a control pancreatic explant expressing the *Crb3^GFP^* reporter, acquired at 1 frame every 10 minutes. The movie is shown as a maximum intensity projection.

**Supplemental Movie 6:** Live imaging of a *Nf2^PancKO^* pancreatic explant expressing the *Crb3^GFP^* reporter, acquired at 1 frame every 10 minutes. The movie is shown as a maximum intensity projection.

**Supplemental Movie 7:** 3D projection of a *Nf2^PancKO^* pancreatic explant cultured for 72 hours. *Nf2^pancKO^* pancreata containing *myrTom* (magenta) and *Crb3^GFP^* (cyan) reporters showing a large vesicle observed within an epithelial cell.

**Supplemental Movie 8:** 3D projection of e14.5 *Nf2^PancKO^* pancreatic section (15µm), stained for CDH1 (green) and MUC1 (magenta). Note large intracellular vesicle.

**Supplemental Movie 9:** Live imaging of a control pancreatic explant expressing the *Crb3^GFP^* and *myrTom* reporters, acquired at 1 frame every 25 seconds over a period of 10 minutes.

**Supplemental Movie 10:** Live imaging of a *Nf2^PancKO^* pancreatic explant expressing the *Crb3^GFP^* and *myrTom* reporters, acquired at 1 frame every 25 seconds over a period of 10 minutes.

**Supplemental Movie 11:** Live imaging of a control pancreatic explant expressing the *Crb3^GFP^* and *myrTom* reporters, acquired at 1 frame every 25 seconds. Green boxes indicate mature lumens, and red boxes indicate nascent lumens.

**Supplemental Movie 12:** Live imaging of a control pancreatic explant expressing the *Crb3^GFP^*, acquired at 1 frame every 25 seconds.

**Supplemental Movie 13:** Control P1 pancreata, stained with MUC1 (yellow). Movie shows sequential optical sections from z-stack.

**Supplemental Movie 14:** *Nf2^PancKO^* P1 pancreata, stained with MUC1 (yellow). Movie shows sequential optical sections from z-stack.

**Supplemental Movie 15:** *Pdx1-Cre*;*Nf2^ff^*; *Yap^f/+^*;*Taz****^f/+^*** P1 pancreata, stained with MUC1 (yellow). Movie shows sequential optical sections from z-stack.

**Supplemental Movie 16:** *Pdx1-Cre*;*Nf2^ff^*; *Yap^ff^*;*Taz^ff^* P1 pancreata, stained with MUC1 (yellow). Movie shows sequential optical sections from z-stack.

## Supplemental Tables

**Supplementary Table 1.**
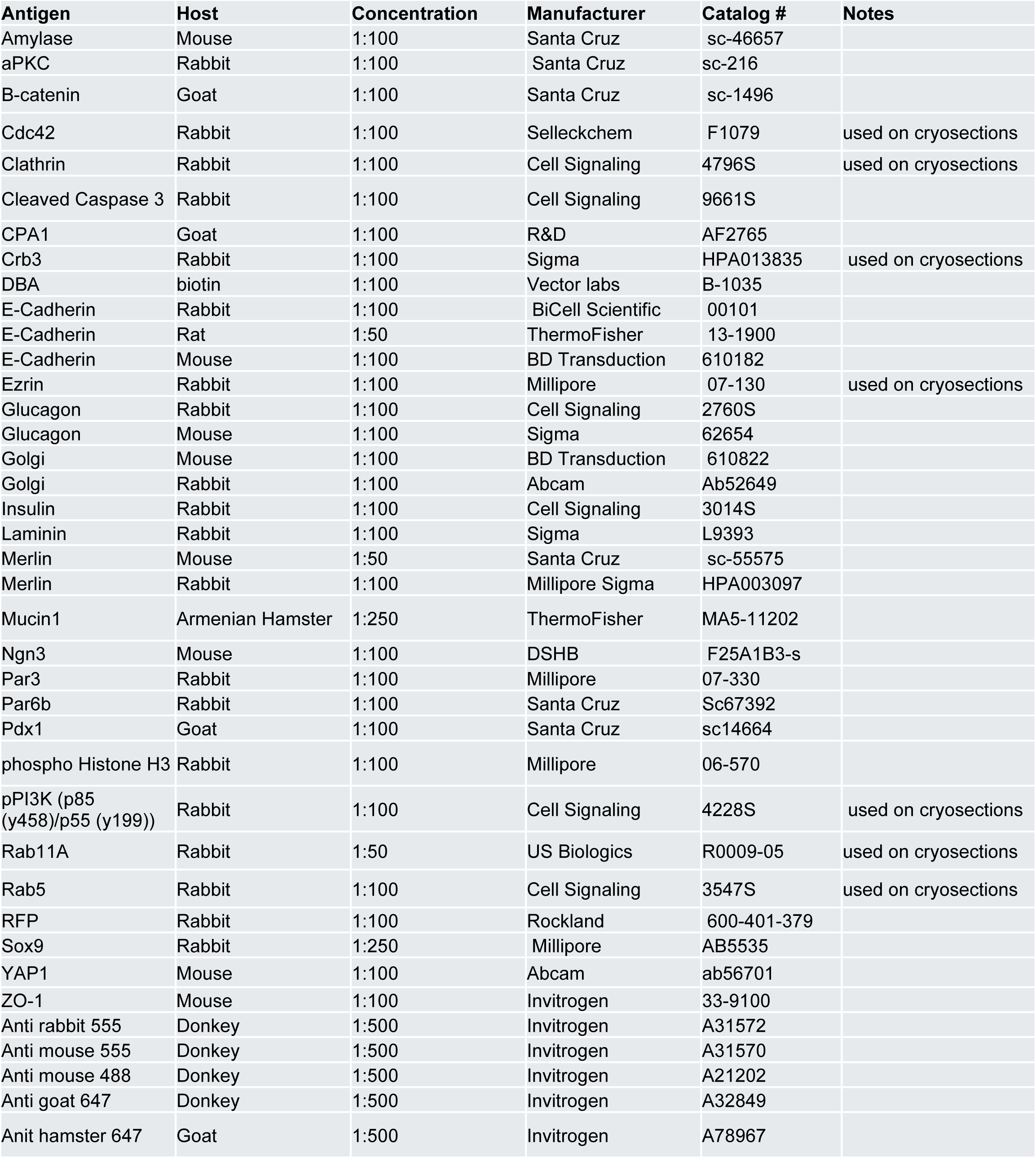
List of antibodies used in this study.

**Supplementary Table 2.**
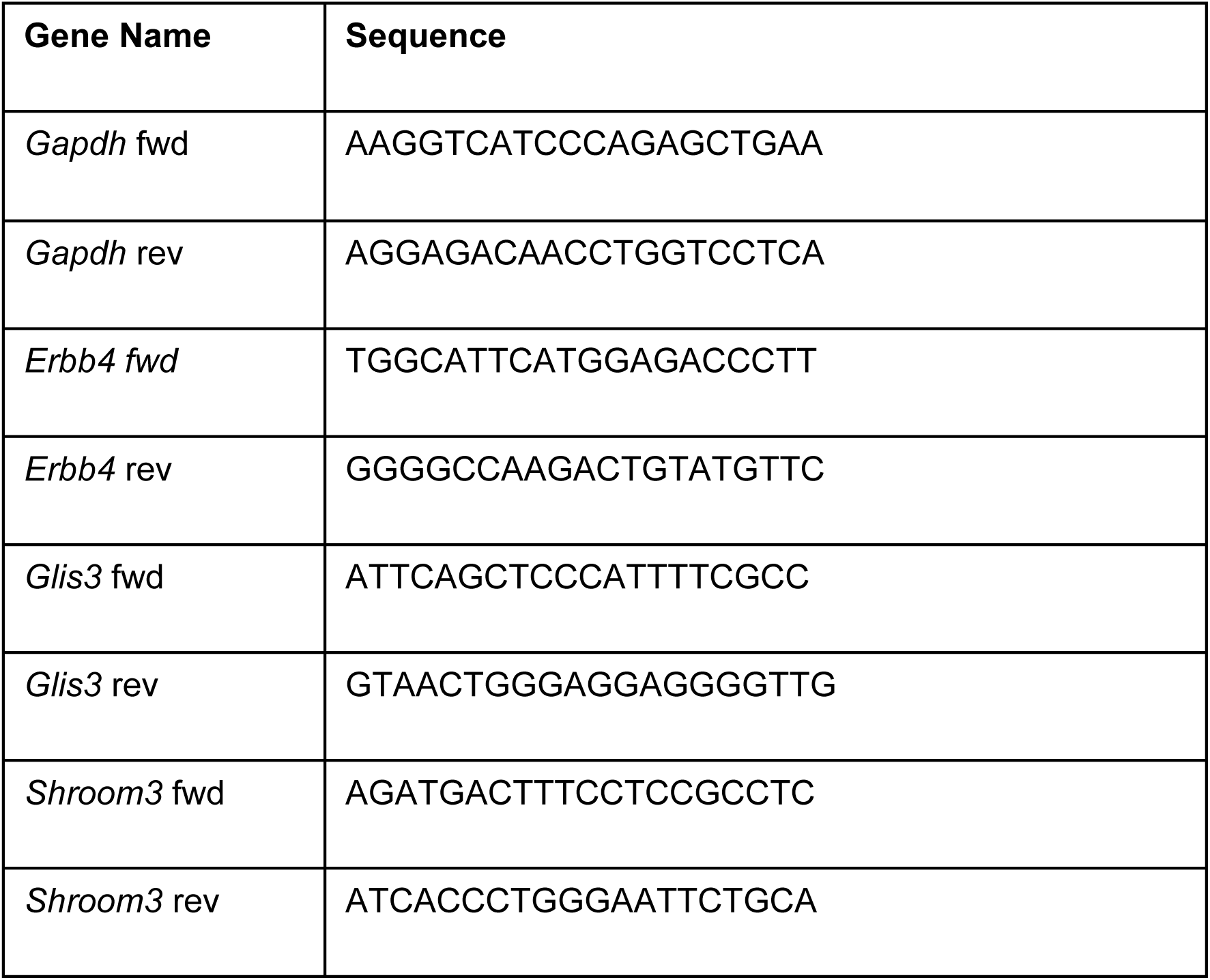
qPCR primers.

## Supplemental Methods

### Image Quantification

#### Pancreatic lineage volume quantifications

Whole-mount immunofluorescence images of pancreata stained for MUC1 and lineage markers (endocrine, CPA1, or DBA) were acquired and analyzed using IMARIS (Bitplane). For each sample, a region of interest (ROI) encompassing the pancreatic epithelium was manually defined based on MUC1 or CDH1 signal. For endocrine and CPA1 quantifications, a three-dimensional MUC1 surface was generated within the ROI to define total luminal volume. For DBA quantifications, a CDH1 surface was generated to define total epithelial volume. Lineage marker-positive volumes were quantified by generating a second surface within the same ROI using a manually adjusted intensity threshold optimized for each sample to accurately delineate tissue morphology while minimizing background signal. Surface volume measurements were recorded and exported to Excel. Lineage-positive volumes were normalized to total luminal volume (MUC1) or total epithelial volume (CDH1) for each pancreas. Normalized values were analyzed and plotted using GraphPad Prism.

#### Plexus quantitative assessment

To gain quantitative insight into these defects, we utilized the Skeletonize plugin in ImageJ (Fiji) as previously described.32 Confocal image stacks were first processed with a Gaussian blur (sigma = 2.0) and then converted into binary masks using Huang thresholding. Due to differences in staining intensity between control and mutant samples, the threshold was manually adjusted for each image to accurately capture epithelial morphology while minimizing background signal. Skeletonization was performed using the “Skeletonize (2D/3D)” plugin. Network morphology was quantified using the “Analyze Skeleton” plugin to extract parameters including branch number, junctions, endpoints, and total branch length. Plexus junctions were defined as the sum of triple and quadruple branch points detected per pancreas. For P1 data, as a measure of cyst formation, we quantified the number of junction voxels was summed for each pancreas and divided by the total number of junctions.

#### Cell death

Images were analyzed as maximum-intensity projections CC3+ epithelial cells were defined as CDH1+ cells containing a discrete CC3 signal. CC3+ cells lacking CDH1 were excluded. Apoptotic indices were calculated by normalizing CC3+ cells to total CDH1+ cells. FOVs from *Nf2^PancKO^* were selected such that each FOV contained regions of dysmorphic epithelium (cysts present, stratified epithelium).

#### Cell proliferation quantification

E14.5 and P1 sections were stained for CDH1, AMY, and phospho-histone H3 (pHH3). Maximum-intensity projections were analyzed. Proliferating epithelial cells (pHH3+) were normalized to total CDH1+ cells. FOVs from *Nf2^PancKO^* were selected such that each FOV contained regions of dysmorphic epithelium.

#### Adult islet characterization and surface area analysis

Adult sections stained for CDH1, glucagon, and insulin were imaged at 20× or 40× and analyzed as Z-stacks or maximum projections. Islets were manually classified as mixed, mantled, glucagon-only, or insulin-only. Islet surface area was measured in ImageJ using the freehand selection tool.

#### Adult islet cell quantification

Adult pancreatic sections were immunostained for E-cadherin, insulin, and either somatostatin (SS) or pancreatic polypeptide (PP) and imaged at 40× magnification.

Individual islets within each field of view were analyzed as z-stacks in ImageJ. The number of SS+ or PP+ cells was quantified for each islet and normalized to the number of insulin-positive (Ins+) cells.

#### Amylase-positive epithelial cell quantification

Maximum-intensity projections were generated for all images prior to analysis. For each field of view (FOV), the total number of epithelial cells and AMY+cells was quantified, and amylase epithelial density was calculated as the ratio of Amylase+ cells to total epithelial cells.

#### NGN3 quantification

E13.5 sections were stained for CDH1 and NGN3 and imaged at 40× magnification. NGN3+ cells were quantified and normalized to the total number of CDH1–positive epithelial cells per field of view.

#### Stratification Analysis (E14.5 pancreatic tissue)

Light sheet microscopy data was utilized to assess stratification of E14.5 pancreatic tissue. A single optical section from the middle of the z stack was assessed, and the maximum number of cell layers between the basal (LAMA+) and apical (MUC1+) markers was quantified. Between 2-4 FOVs were analyzed per pancreata.

#### Stratification Analysis (Explant)

Stratification in explants was performed as previously described.^1^ In brief, two metrics were quantified – the number of cell layers from the edge of the pancreas to the nearest lumens, and the number of cell layers between lumens. A single optical section from the middle of each pancreas was analyzed, and ten measurements were taken per analysis.

#### Explant morphology

For explant area and clefting analysis, maximum-intensity projections were used. For area measurements, the epithelial region (myrTom or CHD1-positive signal) of each explant was manually outlined, and total area was quantified in FIJI (ImageJ). Clefts were defined as clear invaginations or indentations of the epithelial boundary extending inward from the outer epithelial contour, visible in maximum-intensity projections. The number of clefts was quantified per explant.

#### Number of microlumens

MUC1 staining was performed on E11.5 and 12.5 pancreata, and confocal z-stacks through the entire dorsal bud were obtained. The number of disconnected, isolated MUC1 puncta were manually counted by scrolling through the stack.

### Mosaic and Nf2^PancKO^ cell fate analysis

Confocal z-stacks were obtained through thick vibratome sections for each genotype. Each myrTom positive cell was scored as either positive of CPA1, DBA, or neither based on overlap of myrTom signal and other markers. Cell counter plugin in FIJI was utilized to score each cell. Data was analyzed in GraphPad. For both mosaic and full deletion, heterozygotes (*ptf1aCreERT2,Nf2^f/+^*, *myrTom* ^f/^*^+^*^or ff^ and *pdx1Cre, Nf2^f/+^*, *myrTom* ^f/^*^+^*^or ff^, respectively) were utilized as controls.

#### Intracellular myrTom puncta quantification

Confocal z-stacks were acquired and converted into maximum intensity projections (MIPs) for analysis. Images were processed in FIJI (ImageJ) by applying background subtraction using a rolling ball radius of 20 pixels. Processed images were then thresholded using the Otsu algorithm to generate binary masks of intracellular myrTom signal, and watershedding was applied.

myrTom-positive puncta were quantified using the “Analyze Particles” function in FIJI, with consistent size and circularity (>0.25) parameters applied across all samples. The number of puncta per cell was recorded and used as a measure of intracellular myrTom accumulation. Quantitative data were exported to GraphPad Prism for statistical analysis.

#### Intracellular Rab5/Rab11 puncta quantification (cells)

Confocal z-stacks were acquired and converted into maximum intensity projections (MIPs) for analysis. Images were processed in FIJI (ImageJ) by applying background subtraction using a rolling ball radius of 10 pixels. Processed images were then thresholded using the Otsu algorithm to generate binary masks of intracellular Rab5/Rab11. Puncta were quantified using the “Analyze Particles” function in FIJI, with consistent size (>0.1um) parameters applied across all samples. The number of puncta per cell was recorded and exported to GraphPad Prism for statistical analysis.

#### Rab5/Rab11 total fluorescence intensity quantification (pancreatic tissue)

A single optical section per z-stack, corresponding to the plane with maximal CDH1 signal, was selected for analysis. Individual CDH1⁺ epithelial cells were manually outlined, and mean gray value of Rab5 or Rab11 fluorescence within each cell was measured in FIJI (ImageJ).

#### Rab5/Rab11 apical/basal intensity ratio (pancreatic tissue)

A single optical section per z-stack corresponding to the plane with maximal CDH1 signal was selected for analysis. CDH1⁺ epithelial cells lining MUC1⁺ lumens, defined by a clearly identifiable apical MUC1-positive surface, were analyzed. The apical third of each cell was manually delineated, and mean gray value of the relevant channel was measured in apical and basal regions. The apical-to-basal intensity ratio was calculated for each cell.

### Single-nucleus RNA-seq data analysis

#### Preprocessing and quality control

Raw gene-by-barcode count matrices generated by 10x Genomics Cell Ranger in HDF5 format were imported into R using DropletUtils. ^2^ Gene identifiers were mapped to unique feature names, and cell barcodes were assigned as column identifiers. Unless otherwise noted, preprocessing and downstream analyses were performed using Bioconductor workflows, including scuttle, scater, scran, and bluster.^3,4^

Sequencing reads were processed using Cell Ranger v5.0.1 and aligned to the mm10-2020-A mouse reference genome. Computational analyses were performed using R v4.5.3, Bioconductor v3.21, Python v3.10.18, Scanpy v1.11.5, and scVelo v0.3.3.

Droplet quality was assessed using barcode rank distributions with barcodeRanks. A lower UMI threshold of 1,024 was used to distinguish cell-containing droplets from background. Per-cell quality control metrics were computed using addPerCellQC, including total UMI counts, number of detected genes, and the proportion of mitochondrial transcripts. Mitochondrial genes were identified from Ensembl annotation release 101. Cells were flagged as outliers using median absolute deviation-based criteria for low library size, low feature detection, and high mitochondrial content, with a minimum difference threshold of 0.5 applied to mitochondrial proportions. Cells failing any criterion were excluded.

Putative doublets were identified using scDblFinder with cluster-based information.^5^ In datasets with discrete doublet-enriched clusters, those clusters were removed. In other datasets, clusters with greater than 50% predicted doublets were excluded.

#### Normalization, feature selection, and dimensionality reduction

Normalization was performed using deconvolution-based size factor estimation in scran. ^4^ Cells were pre-clustered using quickCluster, size factors were estimated using computeSumFactors with a minimum mean expression threshold of 0.1, and log-transformed normalized expression values were calculated using logNormCounts.

Highly variable genes were identified using a Poisson-based variance model with modelGeneVarByPoisson. Either the top 10% of genes ranked by biological variance or a fixed set of 4,096 highly variable genes was retained, depending on the analysis. Principal component analysis was performed on highly variable genes using scater.^3^

Low-dimensional embeddings for visualization were generated using UMAP, ^6^ typically with 8 neighbors, 3 components, minimum distance of 0.0, and spread of 1.5. For some analyses, reference-based UMAP was used, in which the embedding was learned on a reference population and query cells were projected into the same coordinate space.

#### Clustering and cell type annotation

Cells were clustered using graph-based approaches. Shared nearest neighbor graphs were constructed from PCA or diffusion map embeddings, followed by community detection using Louvain or Walktrap algorithms. In selected analyses, density-based clustering with HDBSCAN was applied to UMAP embeddings.^7^

Clusters were annotated post hoc based on differential marker gene expression, established pancreatic lineage markers, and comparison with expected embryonic pancreatic cell states. Both full and abbreviated cell type labels were assigned and used for downstream analyses.

### Diffusion map analysis and MAGIC-based imputation

Continuous transcriptional structure was analyzed using diffusion maps.^8,9^ Diffusion maps were computed using a custom R wrapper, calculateDiffusion, implementing the Coifman diffusion map construction. Pairwise distances were computed in PCA space, converted to a locally scaled affinity kernel, sparsified by retaining extended nearest-neighbor neighborhoods, and normalized using anisotropic density correction. Diffusion coordinates were obtained from the generalized eigenvalue problem defined by the graph Laplacian and degree matrix.

For some analyses, calculateDiffusion also generated diffusion-smoothed expression values using an implementation conceptually based on MAGIC.^10^ A Markov transition matrix was iteratively applied to log-transformed expression values until convergence criteria were met or a maximum of 8 diffusion steps was reached. Imputed values were rescaled by gene-specific 99th-percentile expression.

Imputed expression values were used for visualization, AUCell gene set activity scoring, and SCENIC regulon activity analysis, where diffusion smoothing improved detection of coordinated low-abundance transcriptional programs. Differential expression testing, marker gene identification, trajectory inference, and trajectory-based differential expression analyses were performed exclusively using observed expression values.

#### Marker gene identification and gene set analysis

Cluster-specific marker genes were identified using EIGEN, an ensemble marker detection framework that combines Welch’s t-tests, Wilcoxon rank-sum tests, binomial detection tests, and gene set–based enrichment statistics into a consensus ranking using a Borda-style vote aggregation strategy.^11^ For computational efficiency, analyses were limited to a maximum of 8,192 cells when required. Gene set enrichment analysis was performed using MSigDB collections, including KEGG, Reactome, WikiPathways, BioCarta, transcription factor target sets from GTRD, and Gene Ontology biological process terms.^12,13^ Gene sets containing fewer than 15 genes were excluded.

Per-cell gene set activity was quantified using AUCell applied to diffusion-imputed expression values. ^14^ Gene set activity scores were summarized at the cluster or cell type level using eigenGeneSets, which applies the EIGEN vote-based marker-ranking framework to AUCell score matrices rather than gene expression matrices. Regulatory network activity was assessed using pySCENIC. ^14,15^ Gene regulatory networks were inferred de novo using GRNBoost2, followed by motif enrichment–based pruning to identify regulons. Regulon activity was quantified on a per-cell basis using AUCell applied to diffusion-imputed expression values and subsequently summarized at the cluster and cell type levels.

#### Differential expression analysis

Differential expression analyses were performed using the MAST framework ^16^. Genes expressed in fewer than 5% of cells within the tested population were excluded. For cell type-restricted analyses, hurdle models included condition as the primary explanatory variable. For analyses performed across multiple cell types, models included terms for cell type, condition, and their interaction. Differential expression was assessed using Wald or likelihood ratio tests, and p-values were adjusted using the Benjamini-Hochberg false discovery rate procedure. Cell type-restricted analyses were performed by subsetting specific populations prior to model fitting.

#### RNA velocity analysis

RNA velocity was estimated independently for each dataset. Spliced and unspliced count matrices were generated from aligned BAM files using velocyto ^17^ and analyzed with scVelo ^18^. Genes were filtered using a minimum shared-count threshold of 10. Transcriptional dynamics were inferred using the dynamical model implemented in scVelo. Velocity vectors and transition graphs were computed and projected onto low-dimensional embeddings, including diffusion map space.

### Time-embedded diffusion integration across developmental stages

To integrate wild-type E11.5 and E14.5 nuclei across developmental time, a custom time-embedded diffusion framework was implemented in Python. Highly variable genes were selected using the Seurat v3 method with developmental stage treated as a batch variable, and PCA was performed in a shared feature space ^19^.

Pairwise distances were computed in PCA space. Within-stage distances were retained for each developmental stage, whereas between-stage distances were computed only for adjacent developmental timepoints, yielding a block-structured distance matrix that enforces temporal locality. Restricting inter-stage connections to adjacent developmental stages prevents biologically implausible shortcuts between distant developmental states while preserving continuous developmental trajectories and maintaining the temporal ordering of developmental progression.

Distances were converted to an anisotropic diffusion kernel using locally adaptive bandwidths estimated from k-nearest neighbors, with k = 8 and epsilon = 1.0. The graph was sparsified using separate neighbor budgets for within-stage and between-stage edges, typically k_intra = 64 and k_inter = 32. After alpha-normalization to reduce sampling density bias, the resulting kernel was row-normalized to define a Markov diffusion operator.

Diffusion components were obtained from the generalized eigenvalue problem defined by the graph Laplacian and degree matrix. The resulting 50-component time-embedded diffusion map represents a temporally constrained developmental manifold that preserves local transcriptional similarity while preventing spurious long-range connections between non-adjacent stages.

#### Reference-based manifold projection

To compare mutant E14.5 nuclei with wild-type developmental structure, mutant cells were projected into a wild-type reference diffusion manifold using geometric harmonics ^20^. Wild-type cells were used to define the reference manifold to avoid distortion of developmental trajectories by mutant-specific transcriptional programs.

A diffusion map was constructed using wild-type cells in PCA space with top_k = 32, alpha = 1.0, t = 1, and 50 nontrivial diffusion components. A ridge-regularized geometric harmonics model was then fit to map PCA coordinates to wild-type diffusion coordinates, using tau = 10^-3 and an automatically estimated kernel bandwidth. Model fit was assessed by reconstructing wild-type diffusion coordinates from the fitted geometric harmonics model and calculating root mean squared reconstruction error prior to projection of mutant cells. Mutant cells were then projected into the wild-type diffusion coordinate system as out-of-sample extensions.

### Integration using mutual nearest neighbors

As a complementary integration strategy, datasets were integrated using mutual nearest neighbors with fastMNN from batchelor ^21^. Integration was performed using 50 dimensions and k = 8 neighbors. The corrected embedding was used for UMAP visualization and clustering, providing an independent comparison to diffusion-based integration.

#### Trajectory inference and pseudotime analysis

Developmental trajectories were inferred using slingshot on diffusion-based embeddings.^22^ Lineages were defined using cluster labels, with specified root and terminal states, and simultaneous principal curves were fit to model developmental trajectories.

Mutant cells were projected onto wild-type trajectories by nearest-point projection in diffusion space. In integrated analyses, generalized additive models were used to map diffusion coordinates to pseudotime, enabling consistent pseudotime assignment across conditions. Lineage assignment was quantified using both hard assignments based on minimum distance to lineage curves and soft probabilistic assignments derived from distance-based weights.

#### Trajectory-based differential expression and gene dynamics

Gene expression dynamics along developmental trajectories were analyzed using tradeSeq.^23^ Generalized additive models were fit as smooth functions of pseudotime, typically using 6 knots. Genes associated with trajectory progression were identified using association tests, and differences between early and late trajectory states were assessed using start-versus-end tests.

Genes were grouped into modules based on shared expression dynamics using clusterExpressionPatterns, followed by hierarchical merging of related modules. Temporal activation patterns were further analyzed by identifying points of maximal expression change along pseudotime. Stage-specific comparisons were performed by identifying representative pseudotime positions, including density peaks within each stage, and testing for differential expression between these positions.

#### Fate probability and compositional analysis

Lineage commitment was quantified by calculating distances from cells to lineage curves in diffusion space. Probabilistic lineage assignments were derived using distance-based weighting, and entropy was used to quantify uncertainty in fate assignment.

Cell type composition was analyzed by aggregating nuclei across developmental stage, condition, and annotated cell type. Both absolute counts and relative abundances were calculated.

## Notes

### Competing Interest Statement

The authors have declared no competing interest.

